# Molecular insight into the enzymatic macrocyclization of multiply backbone N-methylated peptides

**DOI:** 10.1101/2022.07.21.500988

**Authors:** Emmanuel Matabaro, Haigang Song, Lukas Sonderegger, Fabio Gherlone, Andrew Giltrap, Sam Liver, Alvar Gossert, Markus Künzler, James H Naismith

**Author notes:** correspondence to (MK for protein and peptide production and function, JHN for protein structure and molecular mechanism). equal contribution (Emmanuel Matabaro, Haigang Song).

## Abstract

The enzyme OphP is essential for the biosynthesis of the macrocyclic peptide omphalotin A, a dodecamer with 9 backbone N-methylations produced by the wood-degrading fungus *Omphalotus olearius*. Heterologous expression of OphP and the peptide-precursor protein OphMA in yeast, yields omphalotin A. Thus, Oph P was hypothesized to have a dual function; catalyzing both endoproteolytic release of a peptide intermediate from OphMA, and macrocyclization of the multiply α-N-methylated core peptide with concomitant release of a C-terminal follower peptide. In our *in vitro* activity assays, OphP showed robust endoproteolytic and macrocyclase activity on α-N-methylated peptides but was unable to cleave OphMA. The enzyme had a strong preference for hydrophobic, highly α-N-methylated peptides and an α-N-methylated glycine residue at the P1 site. OphP adopts a canonical prolyl oligopeptidase (POP) fold with a predominantly hydrophobic substrate binding cleft, and a small and hydrophobic P1 binding pocket. We demonstrate that OphP is a POP-type macrocyclase with a specificity and a substrate route to the active site different from other members of the family. These results could be exploited for the biotechnological production of macrocyclic peptides with multiple backbone N-methylations, which are interesting due to their favorable pharmacological properties.

## Introduction

Despite violating Lipinski’s ‘rule of five’ (Lipinski et al., 2001), some peptide macrocycles show potent biological activity. This is because the macrocyclization restricts the bond rotations of the peptide backbone and, thus, the favourable conformation of the peptide has a much lower entropic penalty for target engagement (Marsault & Peterson, 2011; Marti-Centelles et al., 2015). In addition, peptide macrocycles are metabolically more stable as they are intrinsically resistant to proteases, and generally show an improved membrane permeability and oral availability when compared to linear peptides (Giordanetto & Kihlberg, 2014). Macrocycles are in clinical use against G-protein coupled receptors and protein-protein interfaces, latter being traditionally challenging targets for small (Lipinski compliant) molecules (Driggers et al., 2008; Surade & Blundell, 2012). Peptide macrocycles are particularly attractive for drug development due to their modular structure and potential chemical diversity arising from the flexible choice of building blocks (natural and unnatural amino acids, α-N-methylated amino acids, D-amino acids, β-amino acids, heterocyclised amino acids and thioamides). Macrocyclization of peptides can be accomplished either by forming a peptide bond between the N- and the C-terminus or by linking two or more functional groups forming amide, lactone, ether, thioether, disulfide or even C-C bonds (Arnison et al., 2013; Koehnke et al., 2017; Montalbán-López et al., 2021). There are powerful routes to synthetic peptide macrocycles with simple amino acids, but syntheses with more complex building blocks, that can significantly improve the potency and pharmacokinetics, are challenging. As a consequence, natural products remain the main sources of marketed peptide macrocycle drugs (Lee & van der Donk, 2022).

Naturally occurring peptide macrocycles are produced by two broad biosynthetic routes, resulting in non-ribosomal peptides (NRPs) and ribosomally synthesized and post-translationally modified peptides (RiPPs). In the case of NRPs, multidomain megaenzymes, non-ribosomal peptide synthetases (NRPS), catalyse the synthesis of the linear peptides from proteinogenic and non-proteinogenic amino acids using linearly arranged, amino acid specific modules in an assembly line fashion. The macrocylization of NRPs is mediated by a final thioesterase domain of the NRPS in bacteria or a condensation-like domain in fungi (Gao et al., 2012; Kohli et al., 2001; Zhang et al., 2016). Several enzyme classes catalyse macrocyclization of RiPPs, including PatG in patellamide biosynthesis (Donia et al., 2008; McIntosh et al., 2010), GmPOPB in fungal amatoxin (Czekster et al., 2017; Luo et al., 2014), and PCY1 in plant orbitide (Barber et al., 2013; Chekan et al., 2017; Ludewig et al., 2018). Mechanistically, these enzymes first cleave off a C-terminal sequence (the follower peptide) from the ribosomally synthesized precursor peptide, forming an acyl enzyme intermediate which is then resolved by the intramolecunucleophilic attack of the free amino terminus of the peptide substrate to form the macrocycle. The macrocyclase domain of PatG belongs to the S8 family of subtilisin-type serine peptidases (MEROPS database of known proteases and their inhibitors, https://www.ebi.ac.uk/merops/) and recognizes a four-residue follower peptide and a preceding heterocyclic ring residue (Koehnke et al., 2012; Koehnke et al., 2014; McIntosh et al., 2010). PatG tolerates a wide range of peptide substrates but is slow (Oueis et al., 2016; Emilia Oueis et al., 2017; E Oueis et al., 2017). PCY1 and GmPOPB belong to post-proline cleaving prolyl oligopeptidases (POP) of the S9A family of serine peptidases. GmPOPB is a dual function macrocyclase as it sequentially catalyses the removal of a leader and follower sequence from the same peptide substrate (precursor peptide) (Czekster et al., 2017; Czekster & Naismith, 2017; Luo et al., 2014). PCY1 relies on an additional protease to first remove the leader sequence to create the substrate for macrocyclization (Barber et al., 2013). Structural and biophysical analysis shows that both PCY1 and GmPOPB recognise the residues in the follower region for substrate binding and catalysis (Chekan et al., 2017; Ludewig et al., 2018). Butelase 1, a fast and promiscuous cyclase from the asparaginyl endopeptidase (AEP) family, has been reported to recognize the C-terminal Asn/Asp-His-Val motif of its peptide substrate, to cleave off C-terminus and ligate the exposed Asn/Asp to the N-terminus of substrate (Nguyen et al., 2015; Nguyen et al., 2014).

A considerable fraction of naturally occurring macrocyclic peptides is backbone N-methylated at one or multiple amino acids. The function of backbone N-methylation of peptides is to enhance their conformational rigidity, membrane permeability, protease resistance and – most relevant in their use as drugs – oral availability (Chatterjee et al., 2008; Ovadia et al., 2011). GmPOPB and PCY1 have been shown to catalyse the macrocyclization of singly backbone N-methylated linear peptides (Ludewig et al., 2018; Sgambelluri et al., 2018) but no enzyme has been characterised, which can efficiently macrocyclize multiply backbone N-methylated peptides. Synthesis and macrocyclization of peptides with multiple backbone N-methylations by entirely chemical means (Marsault & Peterson, 2011; Marti-Centelles et al., 2015; Yu & Sun, 2013) is challenging. Yet, they have enormous potential, as demonstrated by the most well-known example, cyclosporin A, an orally available natural product of the insect pathogenic fungus *Tolypocladium inflatum* (Olarte et al., 2019) that is used as an immunosuppressant during organ transplantation (Cohen et al., 1984). The macrocycle consists of 11 amino acids, 7 of which are backbone N-methylated and 3 are non-proteinogenic (Dreyfuss et al., 1976). The oral availability is a consequence of its chameleon-like property, where the molecule reversibly switches between a non-polar arrangement for membrane transit and a polar arrangement for water solubility (Alex et al., 2011; El Tayar et al., 1993). The conformational flipping depends on the methylated backbone. Cyclosporin A is synthesised by an NRPS pathway with methylations preinstalled on the building blocks prior to peptide bond formation and ring closure by condensation (Dittmann et al., 1994; Lawen & Zocher, 1990).

The peptide natural product at the center of this study is omphalotin A, a 12-residue macrocyclic RiPP with 9 backbone N-methylations (**Figure 1a**). The molecule is produced by the wood-degrading fungus *Omphalotus olearius* and has anti-nematode activity (Mayer et al., 1997). Two posttranslationally modifying enzymes, a peptide α-N-methyltransferase, OphMA, and a serine peptidase of the S9A family, OphP, contribute to its biosynthesis (Ramm et al., 2017; van der Velden et al., 2017). OphMA catalyses the iterative backbone N-methylation of the amide bonds within the core peptide located at its own C-terminus. The enzyme uses SAM as cofactor and has promiscuity towards proteinogenic hydrophobic, aromatic and small hydrophilic amino acid residues, and even accepts non-proteinogenic residues (Song et al., 2021; Song et al., 2020; Song & Naismith, 2020). Coexpression of OphMA and OphP in the yeast *Pichia pastoris* yielded omphalotin A and swapping of the core peptide in OphMA with sequences for lentinulin A and dendrothelin A led to the production of the respective backbone N-methylated macrocycles (**Figure 1b**), consistent with a promiscuous and functional biosynthetic pathway of the two enzymes (Matabaro, Kaspar, et al., 2021; Ramm et al., 2017) (**Figure 1a**). Based on these data and the sequence similarity between the two enzymes, it was hypothesized that OphP, like GmPOPB (Czekster et al., 2017), possesses two catalytic activities: (1) proteolytic cleavage of the OphMA protein at the N-terminus of the backbone N-methylated core peptide, and (2) proteolytic cleavage of the released precursor peptide at the C-terminus of the core peptide and formation of an acyl-enzyme intermediate between the core peptide and OphP. This intermediate is then resolved by the nucleophilic attack of a water molecule or the N-terminus of the core peptide, leading to the release of the linear or macrocyclized (omphalotin A) core peptide, respectively (**Figure 1a**) (Matabaro, Kaspar, et al., 2021; Matabaro, Song, et al., 2021; Ramm et al., 2017).

**Figure 1.**
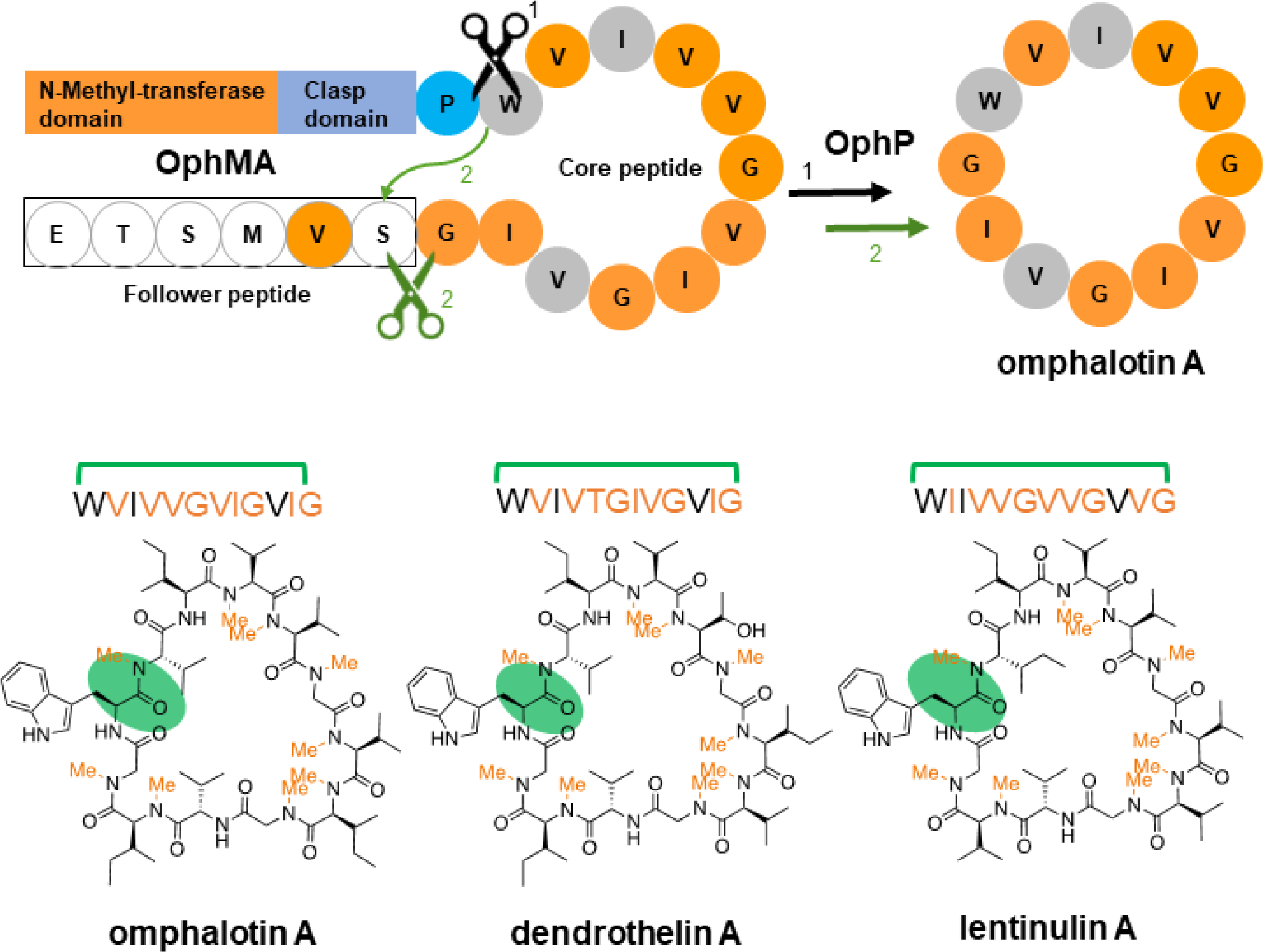
Biosynthesis of omphalotin A inferred from the co-expression of OphMA and OphP in *Pichia pastoris*. (**a**). Proposed biosynthetic pathway of omphalotin A, where scissors indicate proteolytic cleavage (black) and macrocyclization sites (green). (**b**). Sequences and chemical structures of omphalotin A and homologous peptide natural products dendrothelin A and lentinulin A shown with methylation sites in orange and the cyclization site in green. These peptide macrocycles were produced in *Pichia pastoris* by expressing OphP together with OphMA variants with respective C-termini.

In this study, we purified OphP from *P. pastoris* and assessed its *in vitro* activity towards a variety of substrates. We found no evidence for OphP being able to process intact OphMA but detected a robust proteolytic and macrocyclase activity of the enzyme towards multiply backbone N-methylated peptides derived from OphMA. In contrast to other macrocyclase systems, the enzyme appears to be insensitive to the sequence of the follower peptide but have a preference for hydrophobic, multiply backbone N-methylated peptides and recognize a backbone N-methylated glycine residue within the OphMA core peptide as P1 site for proteolytic cleavage. These findings are supported by the structural analysis of enzyme-substrate complexes. This knowledge may enable the exploitation of the enzyme for the biotechnological production of new-to-nature multiply backbone N-methylated peptide macrocycles for applications in agriculture or medicine.

## Results

### OphP proteolytically cleaves and macrocyclizes OphMA-derived, multiply backbone N-methylated peptides *in vitro*

OphP was purified from *P. pastoris* as an N-terminal His_8_SUMO* fusion protein (**Figure 1-figure supplement 1**) and confirmed to be catalytically active using a standard chromogenic substrate for POPs, benzyloxycarbonyl-Gly-Pro-p-nitroanilide (Z-Gly-Pro-4-pNA) (Nagatsu et al., 1976) (**Figure 1-figure supplement 2**). OphP showed optimal activity at 30°C and pH 6.0. The reaction was dependent on enzyme and substrate concentration. OphP activity was reduced by Z-Pro-prolinal (ZPP), a common covalent inhibitor for POPs, but the enzyme was not inhibited as strongly as other members of the family (**Figure 1-figure supplement 2**) (Yoshimoto et al., 1985). Mutation of the putative active site Ser residue at position 580 to Ala (S580A) abolished the detectable enzymatic activity of the fusion protein *in vivo*. Co-incubation of catalytically active OphP with intact OphMA protein showed no detectable peptide product (LC-MS) nor did we observe any mass loss of the protein (ESI-MALDI-TOF) (**Figure 1-figure supplement 3**). Intact OphMA was expressed as an N-terminal His_8_-fusion protein in *E. coli* for 72 hours and purified as previously described (Song et al., 2020; Song et al., 2018; van der Velden et al., 2017). The purified protein is a mixture of different species regarding the degree of backbone N-methylation with the predominant protein species containing 10 methylations (includes one residue in the follower peptide) (**Figure 1a**). The experiment was repeated by purifying OphMA with a lower degree of methylation (predominantly zero, one or two methylations) and again we detected no turnover by OphP. Addition of SAM *in vitro*, which led to conversion of OphMA to the completely methylated state over time, to the reaction, did not change this negative result. Based on these data, we conclude that OphP is not able to process the intact OphMA protein.

To characterize the presumed peptidase and/or macrocyclization activity of OphP, we tested a variety of backbone N-methylated, OphMA-derived C-terminal peptides, consisting of clasp domain, core peptide and follower peptide sequences (**Figure 1a**), for *in vitro* processing by OphP. Since, in our hands, the chemical synthesis of methylated or unmethylated OphMA-derived C-terminal peptides failed (an issue we attributed to their highly hydrophobic character), the peptides were generated by cleaving purified N-terminally His_8_-tagged recombinant wildtype OphMA using trypsin (Oph-30mer), and variants thereof containing cleavage sites for the tobacco etch virus (TEV) protease in the unmethylated clasp domain at various distances from the core peptide, using TEV protease. Some TEV-cleavable OphMA variants also contained deletions and mutations in the C-terminal core and follower peptide sequences. The OphMA variants were expressed for a sufficient period in *E. coli* to allow backbone N-methylation to proceed, purified using metal affinity chromatography and processed by trypsin or TEV protease. The proteolytically released peptides were purified by preparative HPLC (**Figure 2a**). Peptides produced in this way (Oph-30mer, Oph-24mer, Oph-21mer, Oph-18mer, Oph-15mer, Led-21mer and Dbi-21mer), included between three and 12 residues of the unmethylated clasp domain preceding the core peptide. Whereas all other peptides contained the complete 12-residue core and 6-residue follower peptide, Oph-18mer^Tr^ and Oph-15mer contained a truncated core peptide lacking the three C-terminal residues and a truncation of the entire follower peptide, respectively. Led-21mer and Dbi-21mer, contained the 12-residue cores and the follower peptides (SVVSSA) of the lentinulin A and dendrothelin A precursor proteins, LedMA and DbiMA1 (Matabaro, Kaspar, et al., 2021; Matabaro, Song, et al., 2021), respectively (**Figure 2a, Figure 2-figure supplement 1**). MS analysis indicated that the Oph-30mer, Oph-24mer, Oph-21mer, Dbi-21mer, and Led-21mer peptides contained up to ten methylations, both the Oph-18mer^Tr^ had up to 8 methylations and the Oph-15mer carried 7 methylations (**Figure 2-figure supplements 1, 2**). All our attempts to produce and purify non-methylated peptide substrates, using either the catalytically inactive R72A mutant of OphMA or maltose-binding protein fusions of C-terminal parts of OphMA, failed due to the insolubility of the peptides.

**Figure 2.**
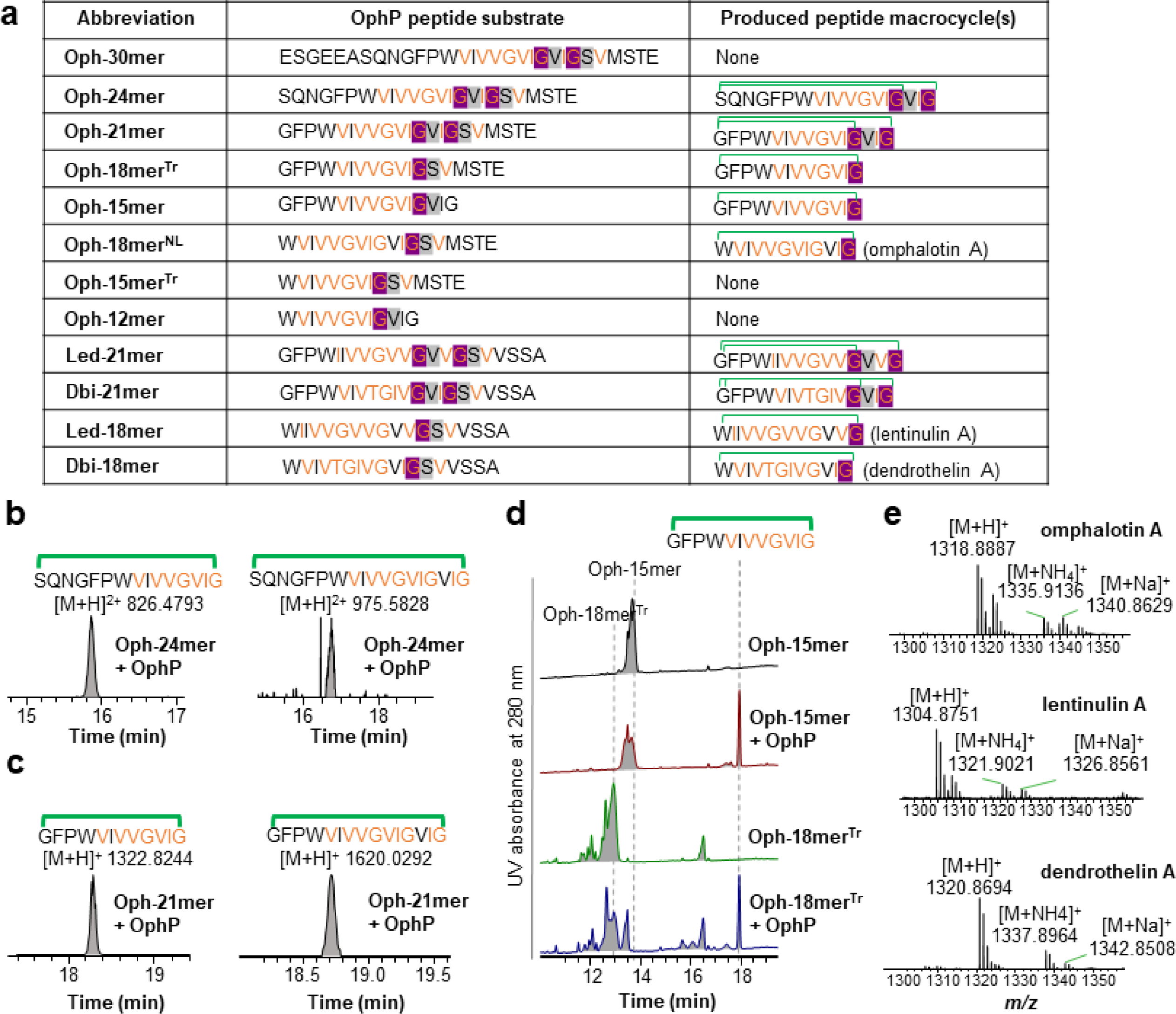
Macrocyclization of OphMA-derived peptide substrates by OphP *in vitro*. (**a**) Sequences of peptides produced in this work, with α-N-methylated residues highlighted in orange. Note that methylation by OphMA proceeds from N- to C-terminus and that only the completely methylated forms of the peptides are displayed. The P1 residues recognized by OphP are highlighted in purple and the P1’ residues in grey. Leader peptide removal was achieved by incubating TEV protease-cleavable OphMA variants with TEV protease and GmPOPB or ProAlanase. Macrocyclization of the peptides by OphP is indicated by green brackets above the peptide sequence. Note that only the macrocyclic products are shown. (**b and c**) Extracted ion chromatograms (EICs) of macrocyclic peptides produced by incubation of Oph-24mer (**b**) or Oph-21mer (**c**)with OphP. (**d**) RP-HPLC UV absorption profiles at 280 nm for Oph-15mer or Oph-18mer^Tr^ in the absence and presence of OphP showing the production of the same macrocyclic peptide. (**e**) MS-analysis of omphalotin A and its orthologues lentinulin A and dendrothelin A produced from Oph-18mer^NL^, Led-18mer^NL^ and Dbi-18mer^NL^ using OphP. The MS chromatograms show masses of both protonated cyclic peptides and corresponding sodium and ammonium adducts.

With the exception of Oph-30mer, where no product could be detected, incubation of all these peptides with OphP led to the formation of peptide macrocycles where the unmethylated residues of the clasp domain that precede the methylated core peptide in OphMA, were incorporated into the product (**Figure 2a, 2b**). The Oph-24mer peptide thereby yielded a mixture of the 15-residue macrocycle cyclo(SQNGFPW^me^VI^me^V^me^V^me^G^me^V^me^I^me^G) ([M+H]^+^=1651.958) and the 18-residue macrocycle cyclo(SQNGFPW^me^VI^me^V^me^V^me^G^me^V^me^I^me^GV^me^I^me^G) ([M+H]^+^=1949.163) (^me^ denotes backbone N-methylation (**Figure 2b, Figure 2-figure supplement 3**). Similarly, the Oph-21mer peptide yielded the 12-residue macrocycle cyclo(GFPW^me^VI^me^V^me^V^me^G^me^V^me^I^me^G) ([M+H]^+^=1322.824) and the 15-residue macrocycle cyclo(GFPW^me^VI^me^V^me^V^me^G^me^V^me^I^me^GV^me^I^me^G) ([M+H]^+^=1620.0293) (**Figure 2c, Figure 2-figure supplement 3**). The Oph-18mer^Tr^ and −15mer peptides both produced the 12-residue macrocycle cyclo(GFPW^me^VI^me^V^me^V^me^G^me^V^me^I^me^G) (**Figure 2d**). These results suggest that OphP is able to *in vitro* macrocyclize OphMA-derived, multiply backbone N-methylated peptides at two different sites within the methylated core but unable to remove the unmethylated clasp domain residues resulting in peptide macrocycles including these residues.

In order to test whether OphP is also able to produce macrocycles consisting only of the methylated core peptides, we exploited the fact that the clasp domain ends with a proline residue in OphMA, LedMA and DbiMA1 and treated derived peptides with the proline-specific endopeptidases recombinant GmPOPB (Czekster et al., 2017) or ProAlanase (Promega, Madison WI, USA). In detail, these were Oph-21mer (leading to 18mer^NL^; ^NL^ denotes ‘<u>n</u>o leader’), Oph-18mer^Tr^ (leading to Oph-15mer^Tr^), Oph-15mer (leading to Oph-12mer), Led-21mer (leading to Led-18mer) and Dbi-21mer (leading to Dbi-18mer). Treatment of the purified peptides with these enzymes removed the unmethylated residues preceding the methylated core peptides (**Figure 2a, Figure 2-figure supplement 4**). Subsequent incubation of Oph-18mer^NL^, Led-18mer and Dbi-18mer with OphP led to the formation of omphalotin A, lentinulin A and dendrothelin A, respectively (**Figure 2e**). Noteworthy, in all reactions where peptide macrocycles were detected, also linear peptide products were found (e.g. **Figure 2-figure supplement 2, 4 and 6**). This is typical for peptide macrocyclases and a consequence of the inherent endopeptidase activity of these enzymes (Czekster et al., 2017). Incubation of Oph-15mer^Tr^ and Oph-12mer with OphP only produced linear peptides (W^me^VI^me^V^me^V^me^G^me^V^me^I^me^G), suggesting OphP cannot form a 9-residue macrocycle. These results are in accordance with the detection of linear 9-residue core peptides in a recent *in vivo* study (Matabaro, Kaspar, et al., 2021).

To test whether OphP could process known peptide substrates for POPs, GmPOPB peptide substrates, namely the 25mer amatoxin precursor (AMA1) and the 17mer phalloidin precursor peptide (PHA1) were chemically synthesized. Consistent with the previous report (Czekster et al., 2017; Luo et al., 2014), GmPOPB converted both substrates and produced cyclic octapeptide from AMA1 and linear peptides in the case of PHA1 (**Figure 2-figure supplement 5**). In contrast, neither OphP nor its homolog LedP was able to process these non-native peptides. These results are in agreement with the specificity of OphP (and LedP) for α-N-methylated peptide substrates.

Taken together, our results show that OphP is able to proteolytically cleave and macrocyclize OphMA-derived peptides of 12 to 24 residues containing a multiply backbone N-methylated core of at least nine residues. The OphP-mediated macrocyclization reaction involves the proteolytic clipping of three to six unmethylated or partially methylated residues following the methylated core peptide and produces peptide macrocycles of 12 to 18 residues that can include stretches of up to six unmethylated residues preceding the methylated core peptide. On the other hand, OphP appears to be unable to remove unmethylated residues preceding the methylated core peptide or process intact OphMA.

### OphP peptide cleavage and macrocyclization activity has a strong preference for an α-N-methylated glycine residue at the P1 site

A closer analysis of the data revealed that peptide processing by OphP depended on the presence of an α-N-methylated glycine (^me^Gly) residue at the P1 site and an unmethylated residue at the P1’ site. This preference is particularly evident on the processing of Oph-18mer^NL^ and Oph-15mer by OphP (**Figure 3a**). MS analysis of purified Oph-18mer^NL^ showed predominantly 10 methylations with 3- to 11-fold methylated species being present in lower amounts (11^th^ methylation of the Met414 residue in the follower peptide) whereby the methylation pattern of the various species can easily be inferred since methylation of the core peptide by OphMA occurs sequentially from the N- to the C-terminus (van der Velden et al., 2017). Incubation of Oph-18mer^NL^ with OphP showed that OphP consumed all species but the 3-, 5- and 6-fold methylated ones. The peptide species consumed by OphP, all have a ^me^Gly followed by an non-methylated residue while the three non-processed ones do not. These results were confirmed by the *in vitro* processing of a 1:1 mixture of 6- and 7-fold Oph-15mer. It should be noted that 7-fold methylation is the maximal degree of methylation that is achievable for a OphMA lacking the 6-residue follower peptide (Song et al., 2020). While the 7-fold methylated peptide species was readily consumed by OphP, the 6-fold methylated one was hardly diminished (**Figure 3a and Figure 3-figure supplement 1**). These results were confirmed by analyzing the processing of another batch of Oph-15mer where purified OphMA was incubated with SAM *in vitro* before cleavage by TEV protease and HPLC-purification of the peptide (**Figure 3-figure supplement 2**). This procedure is known to increase the degree of core peptide methylation (Song et al., 2018) and, accordingly, yielded Oph-15mer with a 1:9 ratio of 6-fold:7-fold methylated peptide. MS analysis of consumption of this batch of Oph-15mer by OphP and the formation of the respective macrocyclic products, cyclo(GFPW^me^VI^me^V^me^V^me^G^me^V^me^I^me^G) and cyclo(GFPW^me^VI^me^V^me^V^me^G^me^V^me^IG), demonstrated that the 7-fold methylated species was strongly preferred over the 6-fold methylated one (**Figure 3b,c and Figure 3-figure supplement 2**). The only difference between the two peptide substrates is the α-N-methylation of the glycine residue at the P1 site of the proteolytic cleavage.

**Figure 3.**
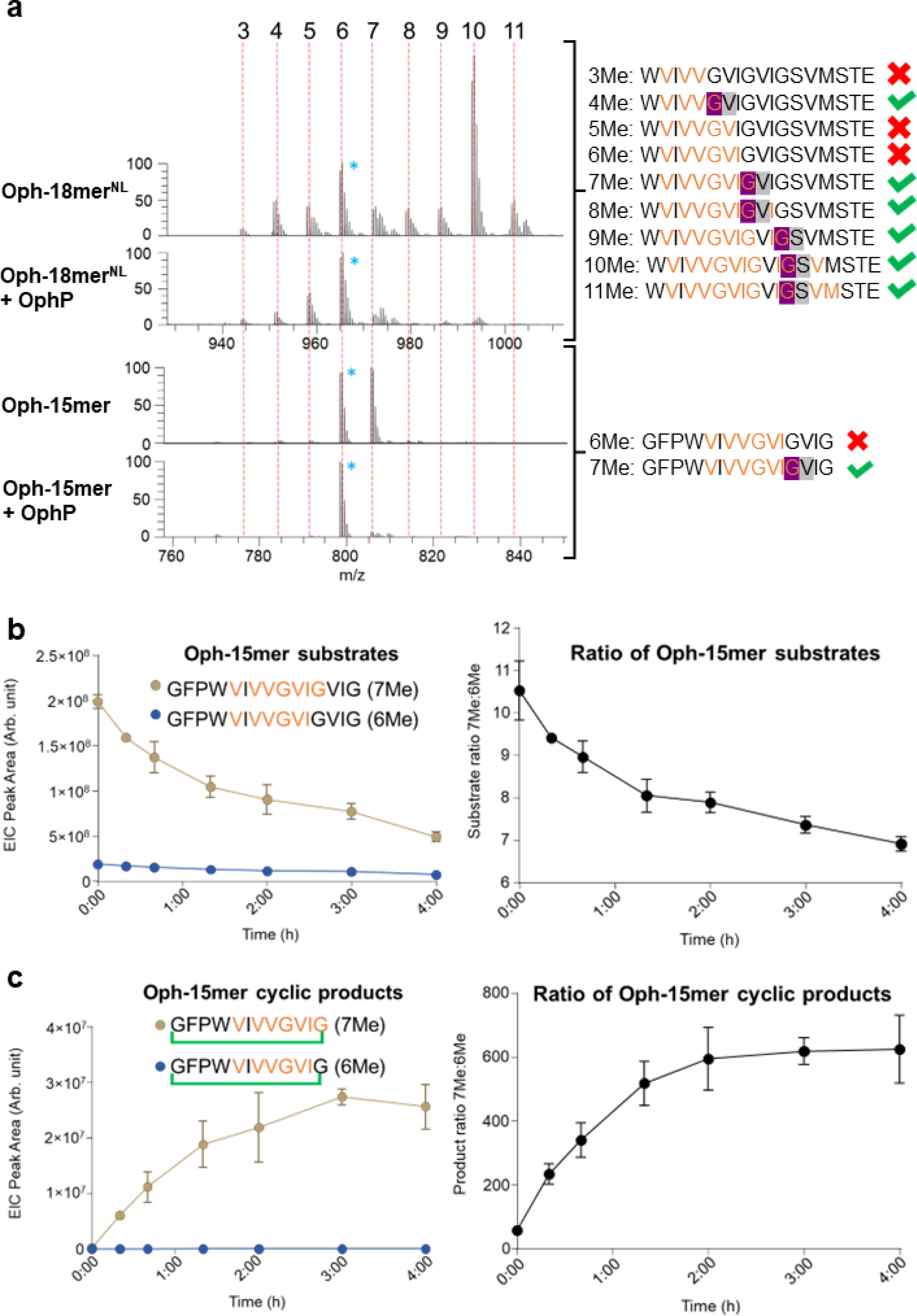
Substrate preference of OphP. (**a**) Mass spectrometric analysis showing ion chromatograms of peptide substrates Oph-15mer and Oph-18mer^NL^ before and after processing by OphP. Red dashed lines indicate the various methylation states of the individual peptides. For comparison, the peak height of the most abundant non-reactive species was drawn at the same level (6-fold methylated species of Oph-15mer and Oph-18mer^NL^). These species were marked with blue asterisks. The sequence of the various peptide species and their processing by OphP is indicated on the right. Green ticks stand for processing and red crosses for lack of processing by OphP. (**b,c**) Time course of *in vitro* reaction between 10µM OphP and 100µM total Oph-15mer Substrate (mixture of 6Me and 7Me species) in HEPES buffer at pH 7.0. EIC peak areas of 6- and 7-fold methylated peptide substrate (**b**) and product (**c**) species were measured at 7 time points over 4 hours. Left panels show the absolute EIC peak areas and the right panels the ratio between the 7- to 6-fold methylated peptide species over the 7 time points.

### OphP adopts a typical POP-fold with an unusual hydrophobic tunnel

OphP shares 35% and 38% sequence identity to PCY1 from *Saponaria vaccaria* and GmPOPB from *Galerina marginata*, respectively, including conservation of the putative catalytic triad residues Ser580, Asp665 and His701 (**Figure 4-figure supplement 1**). Despite these similarities, OphP has two remarkable features that distinguish it from these and other members of the POP family: its preference for multiply backbone N-methylated substrates including a ^me^Gly residue (rather than a proline residue) at the P1 site, and the apparent lack of a site for the recognition of the substrate C-terminus (follower peptide) (**Figures 2 and 3; Figure 4-figure supplement 2**). In order to elucidate the structural basis of these differences, we determined the crystal structures of OphP alone (apo-structure) and in complex with substrates and inhibitors.

We first determined the structure of the putatively catalytically inactive OphP variant S580A. The structure of the apo-form was determined in space group P1 (eight monomers in the asymmetric unit) to 1.9 Å by molecular replacement using GmPOPB structure (PDB entry 5N4C) as the search model. Visual analysis and the PISA server suggest OphP is a monomer in the crystal and other POP enzymes. The eight monomers differ only in flexible loops some of which are involved in crystal contacts. Similar to other POPs, the monomer has two domains, an α/β hydrolase domain (residues 1–82, 453–738) and the seven-bladed β-propeller domain (residues 83–452) (**Figure 4a**). All the monomers in the asymmetric unit adopt a “closed” conformation with the two domains packed against each other which closes off the direct route to the active site. OphP is most similar to GmPOPB (5N4B) with a root-mean-square-deviation (RMSD) of 1.5 Å over 686 residues and closely related to PCY1 (5O3W, RMSD of 1.7 Å over 650 residues) with a more distant relation to porcine muscle POP (1QFS, RMSD of 1.7 Å over 650 residues). Examination of the surface of OphP revealed a large and neutral, hydrophobic tunnel in the middle of the β-propeller domain that connects to the putative active site with its key residue S580 (**Figure 4b**). His701 of OphP sits in a flexible loop region (Leu697–Thr707) which is only resolved in two chains (chain A and chain H) where it adopts different conformations (**Figure 4-figure supplement 3**). In both subunits His701 is around 15 Å away from Ser580 and, thus, a structural rearrangement is required to form the canonical catalytic triad.

**Figure 4.**
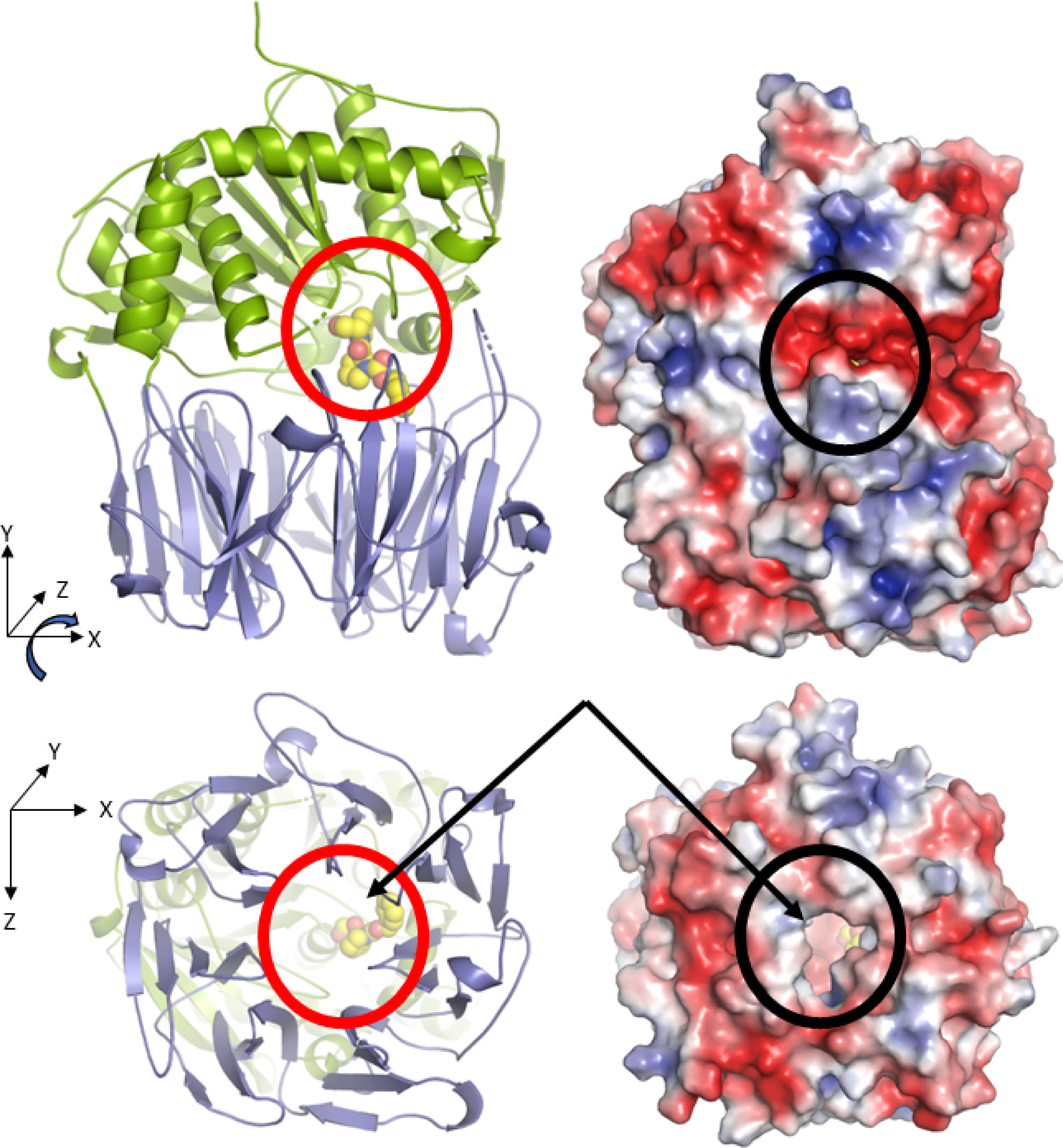
Crystal structures of OphP and possible substrate tunnel. (**a**) In all structures in this study OphP is found with the two domains close together (so called closed state). The β-propellor domain is in blue and the N-terminal domain in green. The active site is located at the interface of the domains. A molecule of the inhibitor ZPP (carbon atoms yellow, oxygen red, and nitrogen blue) shown in sphere form has been placed at the active site circled in red to aid visualization. Experimental details of the ZPP complex structure are shown **Figure 4-figure supplement 3**. On the right is the electrostatic surface calculated APBS-PDB2PQR software suite(Jurrus et al., 2018) under the default setting and drawn in PyMol. The electrostatic potential is set at ± 5 kT/e. There is no access to the active site (black circle) from the side of the structure in the closed state. (**b**) Rotation of 90° around the X-axis shows there is a tunnel through the β-propellor domain (circled in red in cartoon). On the right is the is electrostatic surface calculated as above, the tunnel allows access to the active site (black circle). The yellow carbon atoms of ZPP can be seen through the pore in both representations.

The structure of the covalent complex between OphP and ZPP (**Figure 4-figure supplement 3)** was obtained by soaking wild-type OphP crystals with 5 mM ZPP overnight. The structure of the complex was determined at 2.0 Å resolution. Additional electron density, which we fitted as ZPP covalently linked to Ser580 was seen in two subunits (**Figure 4-figure supplement 3**). Crystal packing would appear to limit the ability of the domains to have separated sufficiently to allow ZPP to access the active site. The calculated solvent content of the crystals is below 40 % consistent with visual observation that there are no large solvent channels that would allow large domain movements that do not disrupt crystal packing. The benzene ring of ZPP is located in an aromatic hydrophobic pocket. The other two subunits had weak electron density suggesting low occupancy. The overall structure of the OphP:ZPP complex is largely unchanged from the apo-structure of OphP(S580A) (0.5 Å over 716 Cα atoms. There are differences in loops Leu139–Ala146, Ser164–Met171, Ser179–Met195, Pro222–Gly230, Leu696–Ser706 and Ala622–Tyr664. As a result, His701 has moved closer (7.6 Å to serine) to both Asp665 and Ser580 to form the catalytic triad (**Figure 4-figure supplement 3**). These results confirm Ser580 as the key catalytic residue and support our suggestion of structural rearrangement of loops is required to form the catalytic triad.

### ^me^Gly at the P1 site of the peptide substrate is coordinated by a specific binding pocket

To characterize the interactions of OphP with its presumed natural substrate, crystals of OphP(S580A) were soaked with mainly 10-fold methylated Oph-18mer^NL^ (W_1_^me^V_2_I_3_^me^V_4_^me^V_5_^me^G_6-_^me^V_7_^me^I_8_^me^G_9_V^me^I_11_^me^G_12_S_13_^me^V_14_M_15_S_16_T_17_E_18_) and mainly 7-fold methylated Oph-15mer peptide substrate (G_1_F_2_P_3_W_4_^me^V_5_I_6_^me^V_7_^me^V_8_^me^G_9_^me^V_10_^me^ I_11_^me^G_12_V_13_I_14_G_15_) (**Figure 5a-c and Figure 5-figure supplement 1, respectively**) (both substrates yield dodecameric peptide macrocycles with the underlined residues; the numbering of the residues eases the visualization of the structural arrangement of the peptide in the active site of the protein displayed below). The Oph-18mer^NL^ complex was determined to 2.0 Å and the Oph-15mer complex at 2.5 Å resolution. The protein structures superimpose well with the apo structure with an RMSD of 0.4 Å over 715 Cα atoms (Oph-18mer^NL^) and 0.8 Å over 714 Cα atoms (Oph-15mer). The peptide substrates are located at the interface between the two domains with its N-terminus extending into the β-propeller domain and its C-terminus contacting the hydrolase domain (**Figure 5a; Figure 5-figure supplement 1**). Substrate binding displaced both glycerol and water molecules found in the apo-structure.

**Figure 5.**
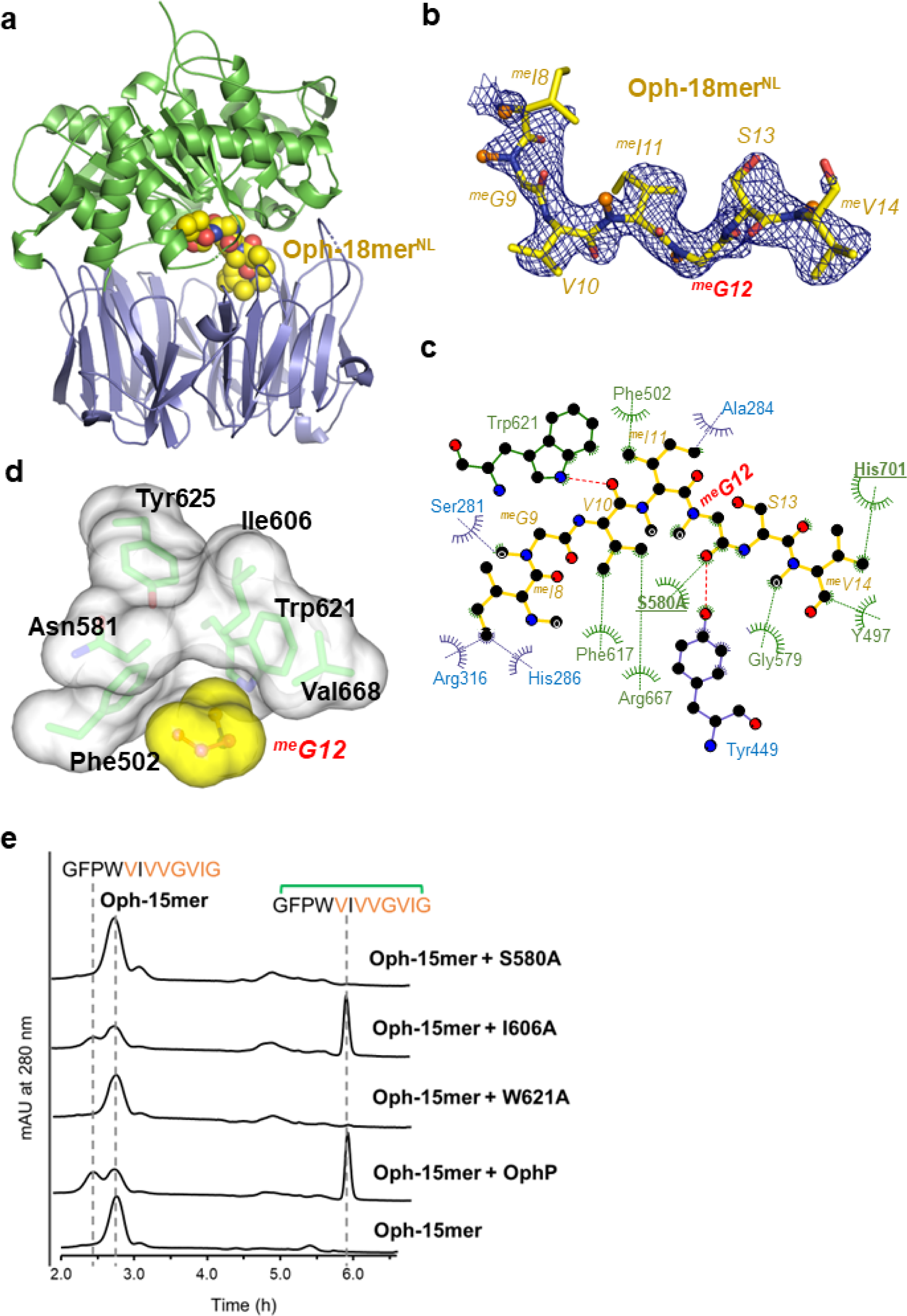
^me^Gly12 recognition site of OphP. (**a**) OphP S580A complex to peptide substrate Oph-18mer^NL^ is shown in spheres. (The color schemes is as Figure 4a) **(b)** The σ_A_-weighted 2mFo-DFc map (contoured at 1σ) for Oph-18mer^NL^ (**c**) LigPlot diagram (Laskowski & Swindells, 2011) showing interactions of the 18mer^NL^ with OphP. **(d)** Surfaces of ^me^Gly12 and its surrounding hydrophobic pocket in Oph-18mer^NL^ complex. The pocket is composed of Tyr625, Trp621, Val668, Phe502, Asn581 and Ile606. (**e**) HPLC analysis of assays using almost pure 7-methylated Oph-15mer as substrate for OphP and its variants. The control reaction without OphP shows the absorption profile of the starting material. While the W621A mutation abolished macrocyclization, I606A showed an activity comparable to wild type OphP. The reactions were performed at 25 °C in buffer containing 50 mM HEPES pH 7.0, 100 mM NaCl and 5 mM DTT at a concentration of 20 µM enzymes and 200 µM Oph-15mer.

For Oph-18mer^NL^, the clearest density for the substrate was found in subunit A. There is some weak electron density for the loop (698–704) which contains the catalytic histidine His701, but we did not model it. Only residues ^me^I8^me^G9V10^me^I11^me^G12S13^me^V14 were fitted to the electron density whilst the remainder were presumed to be disordered (**Figure 5b**). The side chain of ^me^I8 packs against Arg316 whilst ^me^G9 makes almost no contact; the side chain of V10 interacts strongly with the aromatic ring of Phe617, ^me^I11 makes a similar interaction with Phe502 whilst its N-methyl group points to solvent (**Figure 5c**). The P1 binding pocket that accommodates ^me^G12 can be decomposed into two components (amide and side chain) for ease of discussion. The N-methyl group sits in a pocket formed by Phe502, Ile606 and Trp621 (indole makes van der Waal contact) (**Figure 5d**). The Cα of ^me^G12 is surrounded by Ile606, Val668 and Ser580 (< 5 Å). Further away from the Cα of ^me^G12 are the indole of Trp621 (6 Å) and the hydroxyl of Tyr625 (7 Å) which complete the pocket. The peptide bond between ^me^G12 and S13 is positioned for nucleophilic attack by Ser580. The hydroxyl of Tyr499 and the backbone amide of Asn581 are well-positioned to function as the canonical oxyanion hole pair to stabilize the tetrahedral intermediate formed by the nucleophilic attack of Ser580. A different conformer of the side chain of Arg667 would contribute to the stabilization of the negatively charged intermediate. Both Tyr499 and Arg667 are found in many other members of the POP family where they play a similar role^52^.

In the Oph-15mer complex, residues G1F2P3W4^me^V5I6^me^V7^me^V8^me^G9^me^V10^me^I11 were located in the electron density in subunit D (**Figure 5-figure supplement 1)**. The peptide bond of ^me^V8 is closest to Ser580 but its orientation is not consistent with catalysis (**Figure 5-figure supplement 1)**. We conclude this complex is not directly relevant to understanding catalysis but rather is evidence for substrate binding plasticity.

To further investigate the role of Ile606 and Trp621 in substrate recognition and catalysis, we produced OphP variants I606A and W621A. Incubation with Oph-15mer (in almost exclusively 7-fold methylated form) showed that I606A is only slightly less active than the native enzyme while W621A had no activity (**Figure 5e**). We hypothesised that Ile606 might play a role in limiting the volume of the P1 pocket thus introducing selectivity. To test this, we explored whether bulkier side chains at P1 could be processed by the I606A variant of OphP. Unfortunately, we were unable to purify G12V and G12L substrate variants of Oph-15mer and the variant G12A resulted in a complex mixture that prevented further analysis.

## Discussion

The multiply backbone N-methylated peptide macrocycle omphalotin A from *Omphalotus olearius* originates from a RiPP pathway involving the peptide auto-α-N-methyltransferase OphMA and the serine peptidase OphP (Matabaro, Kaspar, et al., 2021; Ramm et al., 2017; van der Velden et al., 2017). Similar macrocyclic peptides, lentinulin A and dendrothelin A, both with nine-fold backbone N-methylations have been isolated from *Lentinula edodes* and *Dendrothele bispora* harboring biosynthetic gene clusters containing OphMA and OphP homologs (Matabaro, Kaspar, et al., 2021) (**Figure 1a**). The simplest model of the biosynthetic pathway is that OphP represents a dual function macrocyclase, that releases a peptide intermediate (core plus follower peptide) from OphMA (function 1) and cleaves off the follower peptide and concomitantly macrocyclizes the core peptide to yield the final product omphalotin A (function 2).

### OphP is not bifunctional

Our *in vitro* experiments with various backbone N-methylated peptide substrates showed oligopeptidase and macrocyclase activity of purified OphP (**Figures 2 and 3 including respective figure supplements**). In contrast, OphP showed no proteolytic activity towards full-length OphMA, irrespective of its methylation state (**Figure 1-figure supplement 3**). OphP was unable to remove unmethylated residues preceding the methylated core peptide, rather in some cases these additional residues were incorporated into the resulting macrocycles (**Figure 2**). We therefore conclude that OphP is not bifunctional and that other enzyme(s) are required to cleave OphMA into the peptide intermediate that then can be macrocyclized by OphP.

Since no candidate for such an endoproteinase has been identified in the gene clusters reported for the biosynthesis of omphalotin A in *O. olearius* (or homologs in *L. edodes* and *D. bispora* (Quijano et al., 2019; Ramm et al., 2017; van der Velden et al., 2017)), the most likely explanation for these results is the existence of a cellular endoproteinase that cleaves OphMA in the clasp region close to the core peptide in order to create the OphP substrate. It has previously been shown that the presumed cleavage site in OphMA is flexible and accessible (Song et al., 2018). In the pathway for the peptide macrocycle segetalin A, an oligopeptidase (OLP1) cleaves off a leader peptide from the precursor peptide to generate the substrate for the POP macrocyclase PCY1(Barber et al., 2013). Thus there is clear precedent for the requirement of separate enzymes. If the cellular endoproteinase responsible for the cleavage of OphMA does not cut after Pro399, additional aminopeptidases will be needed to remove eventual unmethylated residues preceding the omphalotin core peptide and expose Trp400 for macrocyclization of the core peptide by OphP. Based on the sensitivity of the *in vivo* but not *in vitro* production of omphalotin A to proteasomal inhibitors (**Figure 2-figure supplement 6**), the proteasome is a possible candidate for both activities. In agreement with this hypothesis, omphalotin A could be produced by heterologous expression of OphMA and OphP in the eukaryotic host *P. pastoris* (Matabaro, Kaspar, et al., 2021; Matabaro, Song, et al., 2021; Ramm et al., 2017) but not the prokaryotic host *E. coli* (**Figure 2-figure supplement 7**). Further experiments are needed to verify this hypothesis. Since no partially methylated omphalotin A macrocycles have been identified in nature or in expression hosts (Matabaro, Kaspar, et al., 2021; Mayer et al., 1997; Ramm et al., 2017), a critical question arises as to how it is ensured that only the completely methylated OphMA is processed.

### Substrate is routed to the active site most likely through a central hydrophobic channel

The entry of the substrate into active site of POP family enzymes has been proposed to occur by different mechanisms. In one, a hinge between the two domains gives rise to an open (substrate binding/product release) and closed (catalysis) conformation of the enzyme (Canning et al., 2013; Czekster et al., 2017; Kaushik et al., 2014; Li et al., 2010; Shan et al., 2005). Apo OphP(S580A) crystallized in a closed conformation in the absence of ligand (**Figure 4a**). We obtained ZPP, Oph-18mer^NL^ and Oph-15mer co-complexes by soaking the peptide substrates into crystals, with only some small changes in active site loops but no large domain movements. Crystal packing would seem to prevent the large closed to open to closed conformational changes required by the “hinge” mechanism. We cannot exclude the possibility of such gross movements and annealing in the crystal, but it would be unusual. Thus, we rather considered two alternative routes for large peptides to reach the active site. The first is an entrance between the two α/β and β-propeller domains (the side cavity) (**Figure 4a)**. Such side entry has been observed previously, in dipeptidyl peptidase IV, a prolyl-specific exopeptidase (Aertgeerts et al., 2004; Shan et al., 2005; Thoma et al., 2003). However, this side cavity was much larger than that seen in apo OphP. In OphP, the disordered loops Pro222–Gly230 and Leu697–Gly704 would be expected to further constrict the side entry. Furthermore, the protein surface for this route is polar which does not match the nature of the substrate (**Figure 4b)**. We cannot exclude this means of entry but do not consider it as very likely. A second and more plausible route is via the central hydrophobic tunnel (**Figure 4b**) that leads to the cleft. Compared to the other POP enzymes (1NU6, 5O3U and 5N4F), the tunnel in OphP is wider (**Figure 4-figure supplement 4)**. The apparent lack of conformational change, the tunnel width match to substrate size, and the low polarity of this route make the tunnel an attractive route of substrate entry (**Figure 4b**). Entry by this route that has not been observed in any member of the POP family. Interestingly, in a study of porcine muscle prolyl oligopeptidase (Fülöp et al., 1998; Fülöp et al., 2000), the authors proposed the opening of the central tunnel of the β-propeller domain with blades 1 and 7 acting as the gate to allow substrate to gain access to the active site.

### The tight P1 pocket controls selectivity

Detailed analysis of the products derived from the various substrates revealed a ^me^Gly residue as the point of macrocyclization (P1 site) (**Figures 2 and 3 including respective figure supplements**). The requirement for ^me^Gly contrasts the requirement for Pro at P1 which defines the POP family (^me^Gly resembles Pro in the sense that both are tertiary amides). Structural analysis shows no specific interaction of OphP with residues C-terminal to the core peptide (**Figure 5 including respective figure supplements**). Accordingly, the six C-terminal (follower) residues show no binding to OphP (**Figure 4-figure supplement 2**) and there is no conservation in length or amino acid sequence of the follower residue of peptides processed by OphP (**Figure 2a**). We conclude that OphP does not utilize the C-terminal follower residues as a recognition sequence, in contrast to GmPOPB and PCY1 (Chekan et al., 2017; Czekster & Naismith, 2017; Ludewig et al., 2018; Luo et al., 2014).

OphP tolerated some sequence variations within the core peptide to produce omphalotin A, lentinulin A and dendrothelin A (Matabaro, Kaspar, et al., 2021; Ramm et al., 2017) (**Figure 2e**). The enzyme also makes macrocycles of different lengths, incorporating additional residues at the N-terminus (**Figure 2a**). This plasticity is consistent with the observation that the interactions between OphP and the peptide substrate side chains (**Figure 5c)** are dominated by unspecific hydrophobic contacts. Indeed, comparison of the structures of OphP complexed with Oph-15mer and −18mer reveals different side chains of the peptide substrates in the same protein location consistent with recognition plasticity. Furthermore, the different conformations of the catalytic triad residues observed in different structures, indicate considerable flexibility of the pocket and thus capability to adapt to different substrates.

In the Oph-18mer^NL^ complex, ^me^G12 is located at the P1 site consistent with the expected product. The active site configuration creates an appropriately positioned oxy-anion hole for catalysis. ^me^Gly12 binds to a hydrophobic pocket formed by Trp621, Tyr625, Val668, Asn581, Phe502 and Ile606 (**Figure 5d**). The N-methyl group interacts with Trp621 and the mutant W621A is inactive (**Figure 5e**). These favourable hydrophobic interactions are not possible for an unmethylated amide. N-methylation profoundly reduces (Simon et al., 1992) the conformational flexibility of glycine (Lovell et al., 2003; Ramachandran et al., 1963) and, thus, the entropic penalty for binding (^me^Gly12 has Φ and Ψ dihedral angles that are similar to the prolines (P1 site) in structures of PCY1 (PDB 5O3U) and GmPOPB (5N4C)). We suggest this combination of enthalpy and entropic factors drives the preference of OphP (and its relatives) for tertiary amide substrates.

Except for Ile606, the other residues that could interact with a side chain at P1 (Phe502, Val668, Trp621, Tyr625) are strictly conserved across all POP enzymes to date (**Figure 4-figure supplement 1, Figure 5-figure supplement 2**). Isoleucine is almost exclusively found in OphP and closely related methylated peptide macrocyclases (**Figure 4-figure supplement 1, Figure 5-figure supplement 2)**, POP enzymes have the smaller Val in this position. A simple model based on the Oph-18mer^NL^ complex which places a proline residue at P1 result in a clash (distance < 2.5 Å) with Ile606 (**Figure 5-figure supplement 2**), We propose that Ile606 selects against recognition of Pro. In support of this argument, the covalent OphP:ZPP complex, which has a proline group at the P1 site, results in Ile606 adopting an unfavorable conformation (**Figure 5-figure supplement 2**). This correlates with the lower-than-expected inhibitory potency of ZPP against OphP (**Figure 2-figure supplement 6**). The ZPP result does show the selection is not absolute and the pocket can undergo conformational change. Gymnopeptides are members of the borosin family that includes the omphalotins (Quijano et al., 2019) and they possess ^me^Val at the P1 position. The valine side chain would clash not only with Ile606 but also with Val668. The sequences of the OphP homologues of the producing fungus *Gymnopus fusipes* are not known but we would predict their P1 pocket has to be altered from OphP to to accomodate the larger side chain at the P1 position.

In summary, we show that OphP is not a dual function proteinase/macrocyclase and that *in vivo* a third enzyme is almost certainly required for the production of ribosomally synthesized, backbone N-methylated peptide macrocycles from auto-α-N-methylating precursor proteins like OphMA. OphP recognises an N-methylated glycine rather than a proline residue at the P1 site of macrocyclization using a hydrophobic pocket that selects against residues larger than alanine whilst selecting for methylated amides. The hydrophobic and spacious nature of the substrate-binding pocket results in few specific contacts endowing OphP with some substrate promiscuity and the unique ability to cyclize multiply methylated substrates. These unique properties of OphP may turn this enzyme into a valuable tool for producing highly α-N-methylated cyclic peptides for pharmaceutical applications.

## Materials and methods

### Cells, antibodies, and reagents

ProAlanase (mass spec grade) was ordered from Promega. Expression plasmids for recombinant GmPOPB (plasmid PMA 1237: pJ414-GmPOPB) and TEV protease (pRK793_MBP-TEV site-His6-TEV-C4) were previously described (Czekster et al., 2017) and provided by Alvar Gossert (ETH Zürich, Switzerland), respectively. Phusion high-fidelity DNA polymerase was ordered from ThermoFisher Scientific (USA). Peptides AMA1 and PHA1 were chemically synthesized by GenScript Biotech (USA). The chromogenic substrate Gly-Pro-pNA and the POP inhibitor Z-Pro-prolinal (ZPP) were ordered from Sigma-Aldrich. Follower and leader (clasp domain) peptides were synthesized by Genscript.

### Plasmid construction

The plasmids expressing OphP or LedP were constructed from pPIC3.5K-strepII-SUMO*-TEVcs-OphP (PMA1382) and pPIC3.5K-StrepII-SUMO*-TEVcs-LedP (PMA1379) that were previously used in omphalotin A production in *P. pastoris* (Matabaro, Kaspar, et al., 2021). The His_8_-tag was fused to SUMO* by PCR amplification using PMA1379 as a template and His-SUMOSTAR-FW and His-SUMOSTAR-RV as primers. The obtained fragment was cloned into the *Spe*I and *BamH*I restriction sites of plasmids PMA1382 and PMA1379 to generate pPIC3.5K-His_8_-SUMO*-TEVcs-OphP (PMA1596) and pPIC3.5K-His_8_-SUMO*-TEVcs-LedP (PMA1595), respectively. The plasmid coding for His_8_-SUMO*-TEVcs-OphP(S580A) was constructed by exchanging the *Not*I-*Spe*I restriction fragment of PMA1596 by the respective fragment of PMA1374 (pCDFDuet-sMBP-OphP(S580A)). For most of the His_8_-OphMA-TEVcs constructs, one common primer OphMA-FW was used. For plasmids expressing OphMA-TEVcs-ΔVIG, OphMA-TEVcs-DbiCterm and OphMA-TEVcs-LedCterm, primers OphMA-ΔVIG-RV, OphMA-DbiCterm-RV, OphMA-LedCterm-RV were used together with OphMA-FW on PMA1488, respectively. The obtained PCR fragments were digested with *Not*I and *Nd*eI and inserted in the respective sites on the pET24b vector. For the amplification of the coding region of His_8_-OphMA-TEVcs-24mer, primer OphMA-24mer-RV was used on PMA1004. The PCR fragment was inserted into *Nde*I and *Nco*I of PMA1488. In all cases, the plasmids were amplified in *E. coli* DH5α. Plasmid sequences were confirmed by Sanger sequencing. Plasmids encoding His_8_-SUMO*-TEVcs-OphP(I606A), His_8_-SUMO*-TEVcs-OphP(W721A), His_8_-OphMA-TEVcs-ΔC6(G408A), His_8_-OphMA-TEVcs-ΔC6(G408V) and His_8_-OphMA-TEVcs-ΔC6(G408L) were created using published protocols (Liu & Naismith, 2008). Plasmids PMA1488 (His_8_-OphMA-TEVcs-21mer) and PMA1304 (His_8_-OphMA-TEVcs-15mer) were part of our plasmid library. They were used to generate Oph-21mer and −15mer peptides, respectively. PMA1304 and PMA1488 were derived from plasmid PMA1004 (pET24-His_8_-OphMA-cDNA) described earlier (van der Velden et al., 2017). Used primers and plasmids are listed in Supplementary File 1-Tables S1 and S2, respectively.

### Production of recombinant OphP in *P. pastoris*

*Pichia pastori*s GS115 cells were transformed by electroporation using the plasmids pPIC3.5K-His_8_-SUMO*-TEVcs-OphP or pPIC3.5K-His_8_-SUMO*-TEVcs-LedP, linearized by *Bsp*EI for the integration into the *HIS4* locus. Transformants were selected first on a standard SD-HIS minimal medium and then on 250 µg/ml geneticin for high-copy number screening. To produce OphP, cells were cultured at 20 °C in a standard methanol-containing complex medium (BMMY) for three days following the recommendations for *Pichia* protein expression(Matabaro, Song, et al., 2021). For harvesting, cells were separated from the culture medium by centrifugation, and cell pellets were washed with cold lysing buffer (2xPBS) or double-distilled water (dd H_2_O) and centrifuged again at 8000 *× g* for 10 min at 4 °C. Cell pellets were resuspended in cell lysis buffer [2xPBS, 10% glycerol, 1 mM DTT, 20 mM imidazole, and protease inhibitor cocktail (PIC) (Roche)]. Cells lysis was performed on a planetary mill (Pulverisette 7) as previously described (Matabaro, Song, et al., 2021). Cell debris were removed by spinning down the cell lysate at 16, 000 *x g* for 30 min at 4 °C. The cell lysate supernatant was collected and subjected to purification via a Ni^2+^-based immobilized metal affinity chromatography (Ni-IMAC) system using the washing buffer (50 mM HEPES pH 8.0, 10% glycerol and 20 mM imidazole), and elution buffer (50mM HEPES pH 8.0, 10% glycerol and 400 mM imidazole). The purified proteins were stored in storage buffer (50 mM HEPES pH 8.0 + 10% glycerol + 5 mM DTT) at −80 °C. The His_8_SUMO* tag was cleaved off by incubation with TEV protease (1:10, w/w) in assay buffer [300 mM NaCl, 50 mM TRIS pH 8.0, 1mM DTT and 10% Glycerol] overnight, at 25 °C. The reaction mixture was reloaded on a Ni-IMAC system to remove the His_8_-SUMO* tag. The fractions containing OphP were collected and concentrated using an Amicon filter column (50 kDa cut-off). OphP was further polished across an Äkta FPLC system for size exclusion chromatography (SEC) equipped with a Superdex 200 Increase column. For the this final purification step, HEPES buffer [50 mM HEPES pH 8, 1 mM DTT, 10% glycerol (v/v)] was used, as previously described (Matabaro, Song, et al., 2021).

### Protein expression in *E. coli*

OphMA, OphMA-TEV, GmPOPB, and TEV protease were produced in *E. coli* BL21 as previously described (Czekster et al., 2017; Matabaro, Song, et al., 2021). Briefly, chemically competent BL21 cells were transformed with respective expression plasmids and selected on appropriate antibiotics. A pre-culture was incubated overnight in 10 ml at 37 ⁰C, 180 rpm. 5 ml of the preculture was used to inoculate 1 L of terrific broth (TB) medium. The culture was maintained at 37⁰C, 160 rpm until OD_600_ of 1.5 to 2.0. The culture was chilled on ice for 30 min before the addition of 200 µM isopropyl β-D-thiogalactopyranoside (IPTG) and maintained at 16 ⁰C, 160 rpm for 2 days (GmPOPB and TEV) and 3-5 days for OphMA and OphMA-TEV mutants. Cells were harvested by centrifugation at 8000 *× g* for 20min, at 4⁰C. Cells pellets were resuspended in 100 mL ice-cold lysis buffer [300 mM NaCl, 50 mM Tris pH 8.0, 10% glycerol, 1 mM DTT, DNase I, 20 mM Imidazole and protease inhibitor cocktail (Roche)]. For OphMA and OphMA-TEV mutants, NaCl and Tris in the buffer above, were replaced by 50mM HEPES pH 8.0, 0.1% Triton x-100. Cells were lysed by a cold French press, two passages, and clearing of cell debris by centrifugation at 12 000 rpm, 30 min at 4 ⁰C. The cell lysate supernatant was subjected to protein purification either by Ni-IMAC systems (for TEV and GmPOPB) or by Ni-NTA beads (Macherey Nagel). Elution and wash buffers and the SEC method were the same as for OphP purification above. The protein concentration was determined by BCA assay (Thermo Scientific). To confirm the protein quality, 10 µg of each protein was loaded on a 12% SDS gel and run by electrophoresis and stained by coomassie brilliant blue. The purified proteins were shock-frozen in liquid nitrogen and then stored at −80 ⁰C.

### Peptide production and purification

To produce OphMA-C-terminal fragments, purified and TEV-cleavable OphMA proteins were mixed with TEV protease (1:10) to a concentration of 10 mg/ml in assay buffer [300 mM NaCl, 50 mM TRIS pH 8.0, 1 mM DTT and 10 % glycerol]. The reaction was mixed well by pipetting and kept overnight at 25 °C. The reaction mixture was loaded on a pre-equilibrated C8 or C18-SepPak cartridge following the user instruction and recommendations. The peptides were eluted in 1-3 ml 100% methanol depending on the concentration. 5 µl of the eluate was used for HPLC-MS/MS analysis before further purification on an Agilent 1260 infinity preparative HPLC harboring a C18 column (Luna 5 µm C18(2) 100Å, 250X10 mm. The mobile phase consisted of a gradient between water (A) and acetonitrile (B), both supplemented with 0.1% formic acid. The flow rate was set at 1.2 ml/min, and one fraction each minute. The method started with a system equilibration for 5 min at 5% B, followed by a linear gradient up to 40 % B in 10 min, then increased to 98% B in 30 min. The column was finally washed with 98% B for 5 min before gradually dropping to 5% B in a 5 min span. Fractions corresponding to the expected products were collected, and 5 µl of each fraction are tested by HPLC-MS/MS for the presence of peptides. To determine the concentration, samples were analysed by an Agilent 1100 series UV-HPLC calibrated to the bovine albumin serum (BSA) as an internal standard, at λ = 210 nm. To produce Oph-30mer, recombinant His_8_-OphMA lacking a TEV protease cleavage site was digested with trypsin. For this purpose, 100 µg of the purified protein was incubated with 1.25 mg trypsin in 25 µl HEPES buffer pH 8.0, 37 °C, 650 rpm, overnight. 3 µl of the reaction was used for MS analysis. To purify the peptide, C18 cartridges (Sep-Pak C18 1cc (100mg), Waters) were used. The cartridge was first flushed with 8 ml 100% methanol and then equilibrated with 8 ml water. The trypsin reaction mixture was then loaded on the cartridge, followed by 3 washes with 1 mL water supplemented with 1% methanol. The peptides were eluted in 1 mL 100% methanol. The sample was concentrated by evaporating the solvent using a speed vac, and the resulting peptide pellets were resuspended in the assay buffer.

### Protein sequences (the portion used after TEV protease cleavage site (cs) is underlined, the TEV protease cleavage site is in bold)

#### His_8_OphMA-

**TEVcs(21mer)**MEHHHHHHHHTSTQTKAGSLTIVGTGIESIGQMTLQALSYIEAAAKVFYCVIDPATEAFILTKNKNCVDLYQYYDNGKSRLNTYTQMSELMVREVRKGLDVVGVFYGHPGVFVNPSHRALAIAKSEGYRARMLPGVSAEDCLFADLCIDPSNPGCLTYEASDFLIRDRPVSIHSHLVLFQVGCVGIADFNFTGFDNNKFGVLVDRLEQEYGAEHPVVHYIAAMMPHQDPVTDKYTVAQLREPEIAKRVGGVSTFYIPPKARKASNLDIIRRLELLPAGQVPDKKARIYPANQWEPDVPEVEPYRPSDQAAIAQLADHAPPEQYQPLATSKAMSDVMTKLALDPKALADYKADHRAFAQSVPDLTPQERAALELGDSWAIRCAMKNMPSSLLDAARESG**ENLYFQ<u>G</u>**<u>FPWVIVVGVIGVIGSVMSTE</u>

#### His_8_OphMA-TEVcs-ΔVIG(18mer)

MEHHHHHHHHTSTQTKAGSLTIVGTGIESIGQMTLQALSYIEAAAKVFYCVIDPATEAFILTKNKNCVDLYQYYDNGKSRLNTYTQMSELMVREVRKGLDVVGVFYGHPGVFVNPSHRALAIAKSEGYRARMLPGVSAEDCLFADLCIDPSNPGCLTYEASDFLIRDRPVSIHSHLVLFQVGCVGIADFNFTGFDNNKFGVLVDRLEQEYGAEHPVVHYIAAMMPHQDPVTDKYTVAQLREPEIAKRVGGVSTFYIPPKARKASNLDIIRRLELLPAGQVPDKKARIYPANQWEPDVPEVEPYRPSDQAAIAQLADHAPPEQYQPLATSKAMSDVMTKLALDPKALADYKADHRAFAQSVPDLTPQERAALELGDSWAIRCAMKNMPSSLLDAARESG**ENLYFQ<u>G</u>**<u>FPWVIVVGVIGSVMSTE</u>

#### His_8_OphMA-TEVcs-ΔC6(15mer)

MEHHHHHHHHTSTQTKAGSLTIVGTGIESIGQMTLQALSYIEAAAKVFYCVIDPATEAFILTKNKNCVDLYQYYDNGKSRLNTYTQMSELMVREVRKGLDVVGVFYGHPGVFVNPSHRALAIAKSEGYRARMLPGVSAEDCLFADLCIDPSNPGCLTYEASDFLIRDRPVSIHSHLVLFQVGCVGIADFNFTGFDNNKFGVLVDRLEQEYGAEHPVVHYIAAMMPHQDPVTDKYTVAQLREPEIAKRVGGVSTFYIPPKARKASNLDIIRRLELLPAGQVPDKKARIYPANQWEPDVPEVEPYRPSDQAAIAQLADHAPPEQYQPLATSKAMSDVMTKLALDPKALADYKADHRAFAQSVPDLTPQERAALELGDSWAIRCAMKNMPSSLLDAARESG**ENLYFQ<u>G</u>**<u>FPWVIVVGVIGVIG</u>

#### His_8_OphMA-TEVcs-24mer

MEHHHHHHHHTSTQTKAGSLTIVGTGIESIGQMTLQALSYIEAAAKVFYCVIDPATEAFILTKNKNCVDLYQYYDNGKSRLNTYTQMSELMVREVRKGLDVVGVFYGHPGVFVNPSHRALAIAKSEGYRARMLPGVSAEDCLFADLCIDPSNPGCLTYEASDFLIRDRPVSIHSHLVLFQVGCVGIADFNFTGFDNNKFGVLVDRLEQEYGAEHPVVHYIAAMMPHQDPVTDKYTVAQLREPEIAKRVGGVSTFYIPPKARKASNLDIIRRLELLPAGQVPDKKARIYPANQWEPDVPEVEPYRPSDQAAIAQLADHAPPEQYQPLATSKAMSDVMTKLALDPKALADYKADHRAFAQSVPDLTPQERAALELGDSWAIRCAMKNMPSSLLDAA**E NLYFQ<u>G</u>**<u>SQNGFPWVIVVGVIGVIGSVMSTE</u>

#### His8-SUMO*-TEVcs-OphP

MGHHHHHHHHGGSDSEVNQEAKPEVKPEVKPETHINLKVSDGSSEIFFKIKKTTPLRRLMEAFAKRQGKEMDSLTFLYDGIEIQADQTPEDLDMEDNDIIEAHREQIGG**ENLYFQG**TSMSFPGWGPYPPVERDETSAITYSSKLHGSVTVRDPYSQLEVPFEDSEETKAFVHSQRKFARTYLDENPDREAWLETLKKSWNYRRFSALKPESDGHYYFEYNDGLQSQLSLYRVRMGEEDTVLTESGPGGELFFNPNLLSLDGNAALTGFVMSPCGNYWAYGVSEHGSDWMSIYVRKTSSPHLPSQERGKDPGRMNDKIRHVRFFIVSWTSDSKGFFYSRYPPEDDEGKGNAPAMNCMVYYHRIGEDQESDVLVHEDPEHPFWISSVQLTPSGRYILFAASRDASHTQLVKIADLHENDIGTNMKWKNLHDPWEARFTIVGDEGSKIYFMTNLKAKNYKVATFDANHPDEGLTTLIAEDPNAFLVSASIHAQDKLLLVYLRNASHEIHIRDLTTGKPLGRIFEDLLGQFMVSGRRQDNDIFVLFSSFLSPGTVYRYTFGEEKGYRSLFRAISIPGLNLDDFMTESVFYPSKDGTSVHMFITRPKDVLLDGTSPVLQYGYGGFSLAMLPTFSLSTLLFCKIYRAIYAIPNIRGGSEYGESWHREGMLDKKQNVFDDFNAATEWLIANKYASKDRIAIRGGSNGGVLTTACANQAPGLYRCVITIEGIIDMLRFPKFTFGASWRSEYGDPEDPEDFDFIFKYSPYHNIPPPGDTVMPAMLFFTAAYDDRVSPLHTFKHVAALQHNFPKGPNPCLMRIDLNSGHFAGKSTQEMLEETADEYSFIGKSMGLTMQTQGSVDSSRWSCVTV

#### Enzyme activity assays

For the chromogenic substrate and the POP inhibitor, stocks of 50 mM and 10 mM, respectively, were prepared in methanol. Before the reaction, the substrate was solubilized at 40 °C and then the required amounts were mixed with the standard assay buffer [50 mM HEPES pH 6.0 + 10 mM DTT]. The enzyme was added last, to make the total reaction volume 50 µL. The reaction mixture was kept at 30°C while shaking at 600 rpm. The reaction was quenched by the addition of 50 µl methanol. The enzymatic activity was monitored by absorbance measurement at 410 nm (for *p*-nitroanilide) on an Infinite 200Pro M Plex spectrophotometer (Tecan). Continuous reaction monitoring was ensured by incubating the 96-well plates at 30°C, with absorbance measurements each 30 min.

For peptide substrates, reactions were also carried out in 50 µl with a standard buffer [50 mM HEPES pH 7.0, 10 mM DTT], at 30°C, 600 rpm unless stated otherwise. The peptide was first mixed with the buffer before adding the enzymes. Both substrate and enzyme concentrations varied up to 100 µM. The reaction was stopped by the addition of 50 µl methanol, then spun down at the highest speed (20,000 *x g*), 4⁰C. For HPLC-MS/MS analysis, 5 µl supernatant was used. For the synthetic peptides, PHA1 and AMA1 were stored in 1 mg/ml. For the reaction, 20 µM substrate was used with 1 µM of protease. For ProAlanase, a 1:50 enzyme/substrate (w/w) was used in a 25 µl reaction volume, in either 50 mM HEPES (pH 6.0) or 50 mM sodium acetate (pH 4.0). These reactions were carried out at 37°C for 4 h and stopped by adding 25 µl methanol. After centrifugation at the highest speed at 4 °C, 5 µl supernatant was used for HPLC-MS/MS analysis.

For the time course experiment with OphP and Oph-15mer, the reactions were performed in buffer containing 20 mM HEPES (pH 7.0), 100 mM KOAc, 2mM Mg(OAc)2, 10% glycerol and 10 mM DTT at a concentration of 100 μM total Oph-15mer (90 μM 7-fold methylation, 10 μM 6-fold methylation) and 10 μM OphP. The reactions were performed in 70 μl reaction volume at 30°C and at each time point, 5 μl were sampled and added to 20 μl of MeOH to stop the reaction. The samples were centrifuged for 5 minutes at 15000 rcf and 10 μl were injected for HPLC-MS/MS analysis.

For the experiment of OphP with Oph-15mer (mixture of 7-fold and 6-fold methylations) and protease inhibitors (**Figure 2-figure supplement 6**), the reactions were performed in buffer containing 20 mM HEPES (pH 7.0), 100 mM KOAc, 2mM Mg(OAc)2, 10% glycerol and 10 mM DTT at a concentration of 100 μM total Oph-15mer (90 μM 7-fold methylation, 10 μM 6-fold methylation) and 10 μM OphP. DMSO stocks of the different inhibitors were diluted in the HEPES buffer and added to the reaction to the indicated final concentration. The highest used DMSO concentration was used in a control reaction without inhibitors The reactions were performed in 5 μl reaction volume at 30°C and 20 μl of MeOH were added after 30 minutes to stop the reaction. The samples were centrifuged for 5 minutes at 15000 xg and 10 μl were injected for HPLC-MS/MS analysis. For the experiment testing for preferred production of 7-fold over 6-fold methylated species (**Figure 3-figure supplement 2**), the reactions were performed as described above, but with varying Oph-15mer substrate concentrations of 100 μ M and 1000 μ M total Oph-15mer substrate.

For assays of OphP variants with Oph-15mer (almost pure 7-fold methylated peptide), the reactions were performed in buffer containing 50 mM HEPES (pH 7.0), 100 mM NaCl, and 5 mM DTT at a concentration of 200 μM Oph-15mer and 20 μM OphP. The reaction mixtures in Eppendorf tubes were incubated at 25 °C and 1000 rpm using a ThermoMixer C for 18 hours. Reactions were quenched by adding an equal amount of methanol and protein precipitation was removed by centrifugation. The supernatant was analysed using Waters Acquity H-Class plus UPLC equipped with SQ, PDA and ELS detectors. Buffer A (water with 0.1% FA) and buffer B (acetonitrile with 0.1% FA) are used as mobile phase. Typically, 10 µL of supernatant was loaded and separated on a ACQUITY UPLC BEH C18 column (1.7 µM, 2.1 x 100 mm) at a flow rate of 0.6 ml/min with following gradient: 0-0.25 min for 5% buffer B, 0.25-0.4 min from 5% to 40% buffer B, 0.4-3.6 min from 40% to 55% buffer B, 3.6-3.7 min from 55% to 60% buffer B, 3.7-6.0 min from 60% to 70% buffer B, 6.0-6.15 min from 70% to 95% buffer B, 6.15-7.6 min for 95% buffer B.

#### LC-ESI-MS/MS analysis

Peptide analysis was performed by following previous protocols(Matabaro, Song, et al., 2021) for HPLC-MS/MS analysis of borosins. Briefly, data were generated from Dionex Ultimate 3000 UHPLC HPLC coupled to an MS/MS system (Thermo Scientific Q Exactive classic). A list of expected masses (inclusion list) was always included in the HPLC-MS/MS method. The peptide was applied on a C18 column (Phenomenex Kinetex ®µm XB-C18 100 Å (150 x 2.1 mm)) pre-heated at 50 °C. For UV-vis spectra, the signals were recorded by a diode array detector (DAD) set at 210, 230, 250, and 280 nm. Water (solvent A) and acetonitrile (solvent B) both containing 0.1% FA served as a mobile phase, with a 0.5 ml/min flow rate. The HPLC method started by equilibrating the system for 2 min at 5% B, followed by a linear gradient up to 70 % B in 13 min, and then increased to 100% B in 3 min. The column was rinsed first with 100% B for 1 min and then with 5% B for 5 min. The MS/MS system used the heated electrospray ionization in a positive ion mode. The full MS was set as follows: an automatic gain control (AGC) target of 70,000 [AGC target 1e6, maximum ion trap (IT), 120 ms, and scan range of 200-2500m/z]. Data-dependent (dd) MS/MS was performed at a resolution of 17,500 [AGC target at 1e5, maximum IT 120 ms, and isolation window of 3.0 m/z]. The normalized collision energy (NCE) was set at 30% for macrocycles, and 18% or stepped NCE (16, 18, 22%) for linear peptides.

#### ESI-TOF MS and intact mass measurement

Intact protein mass analysis was performed using a Waters Xevo G2-QS QToF MS coupled to an ACQUITY UPLC I-Class LC system (Waters Corporation). The spectrometer was auto-calibrated throughout the experiment using an internal lock-spray mass calibrant (at 556.2771 m/z of leucine enkephalin, 1 s scan every 120 s interval). The data was acquired in Continuum format, in a 300 – 2500 m/z spectral window with a scan time of 1 s and an interscan time of 0.014 s. The mass spectrometer and ESI ionization source were operated under the following parameters: capillary voltage at 3 kV, cone voltage at 20 V, source offset voltage at 40 V, source temperature at 100 °C, desolvation temperature at 400 °C and desolvation gas (nitrogen) flow at 650 L/h, cone gas flow at 30 L/h. In a typical analysis, 1 µL (30 ng) of the protein sample was injected and desalted on a ProSwift™ RP-1S HPLC (250 x 4.6 mm x 5 μm, Thermo Scientific™) column maintained at 60 °C. The elution was performed using solvent A (water with 0.1% formic acid) and solvent B (acetonitrile with 0.1% formic acid) with a gradient elution using isocratic 95% solvent A (0 - 1 min), linear gradient to 95% solvent B (1.1 – 7 min), and isocratic 95% solvent A (7.1 – 10 min) at a flowrate of 0.2 ml/min. Combined mass spectra from the total chromatographic protein peak were used for intact mass reconstruction using MassLynx (Waters) Max(imum) Ent(ropy) 1 deconvolution algorithm (Resolution: 0.5 Da/channel, Width at half-height: ion series/protein-dependent, Minimum intensity ratios: 33% Left and Right). Spectra were deconvoluted between 40000 and 55000 for OphMA.

#### ITC studies of OphP with peptides

Isothermal titration calorimetry (ITC) experiments were performed using a MicroCal PEAQ-ITC (Malvern). Leader peptides and follower peptides (SVMSTE and VIGSVMSTE) were dissolved in DMSO at sufficient concentration as a stock. A typical ITC experiment was performed by repeated injections 2 µL of peptide solution eighteen times into a cell solution containing 0.4 ml of S580A. The measurements were done at 25 °C at a stirring rate of 750 rpm with a reference power (DP) of 5 mcal/s or 10 mcal/s. Control titrations were performed similarly except for the absence of protein in the cell solution. Data processing and fitting were performed using MicroCal PEAQ-ITC analysis software (Malvern).

#### Crystallization, data collection, reduction, and refinement

Freshly purified OphP and S580A proteins were screened for apo crystals using the sitting drop method using commercial screening kits from Hampton Research or Molecular Dimensions. OphP and S580A were eventually crystallized at 40-50 mg/ml by mixing 0.3 µL of proteins with 0.3 µL of the reservoir (0.2 M sodium acetate, 0.1 M Bis-Tris propane (pH 6.0-6.5) and 28% PEG 3350). Crystals appeared within eight days and grew to full size within three weeks at room temperature. Crystals were fished, transferred to cryo-protectant (reservoir solution supplemented with 40% PEG 3350) and flash-frozen in liquid nitrogen. The structure of the OphP-ZPP complex was determined by soaking OphP crystals with 5 mM Z-Pro-prolinal overnight at room temperature. Crystals were in similar cryoprotectant with 1 mM ZPP. For the S580A:Oph-15mer complex, apo S580A crystals were soaked with 1 mM purified Oph-15mer overnight at room temperature. Due to the poor solubility of Oph-18mer^NL^, apo OphP(S580A) crystals were soaked with a cloudy reservoir solution containing 18mer overnight to obtain the OphP(S580A)-18mer^CF^ complex. More than 50 crystals dataset were collected for each complex and screened based on the density.

All the X-ray diffraction data were recorded at Diamond Light Source beamlines (I03, I04, I04-1, and I24). The diffraction images were reduced, integrated, and scaled using xia2(Winter, 2010), DIALS(Winter et al., 2018), Apo structures of S580A were determined by molecular replacement using a Phaser(McCoy et al., 2007) with GmPOPB structure (PDB entry 5N4C) as the initial model. Autobuilding was performed using Buccanner from the molecular replacement solutions and further refined using Refmac5(Murshudov et al., 1997) and manually built with Coot(Emsley et al., 2010), and improved using PDB-redo(Joosten et al., 2012). Simulated omit maps for the ligands were calculated using Phenix(Liebschner et al., 2019). All the other structures were solved by molecular replacement using apo S580A structure as initial models, refined, manually built and improved similarly. All crystallographic figures were generated using Pymol (Schrödinger LLC), CCP4mg (McNicholas et al., 2011) or Chimera(Pettersen et al., 2004). Structure-based sequence alignment was created using MUSCLE (Edgar, 2004) and ESPript 3.0 (Robert & Gouet, 2014). Full crystallographic details are given in Table 3.

## Supporting information

Fig2SOI

Fig3SOI

Fig5SOI

Fig4SOI

Tables SOI

Fig1SOI

## Acknowledgements

We would like to thank Prof. Jörn Piel and his staff, in particular, Dr. Clara Chepkirui and Dr. Silke Probst for their support in the HPLC purification, and Philip Rößler for supplying TEV protease and helping with protein purification. We are grateful to Dr. Chris Arter from Rosalind Franklin Institute for the assistance with the MS experiments and Dr. Xiaping Fu for the assistance with the purification of peptides. We would also like to thank Diamond Light source for the access to the synchrotron. This work was funded by the Swiss National Science Foundation (Grant no. 31003A-173097), ETH Zürich and UK BBSRC (BB/R018189/1).

## Data availability

Crystallographic and structural data reported in the paper are available from the RCSB; 7ZB2 (apo-structure), 7ZAZ (ZPP complex). 7ZB0 (Oph-15mer complex) 7ZB1 (Oph-18mer complex). Raw data for the Figures are shown in Supporting Information.

