## Supplementary material for "Molecular insight into the enzymatic macrocyclization of multiply backbone N-methylated peptides": Fig2SOI

**Figure 2-figure supplement 1**. **HPLC-MS analysis of OphP substrates.** (**a, b**) Ion chromatograms of OphMA-derived, backbone N-methylated peptide substrates produced using TEV protease-cleavable OphMA-variants and TEV protease only (a) or a combination of TEV protease and GmPOPB (b). The number of methylated residues is indicated by green numbers and red dashed lines to facilitate a comparison. The different peptides have been aligned by extent of methylation rather than by absolute mass. The core peptide is underlined. Note that the follower peptide is different for the non-omphalotin peptides.

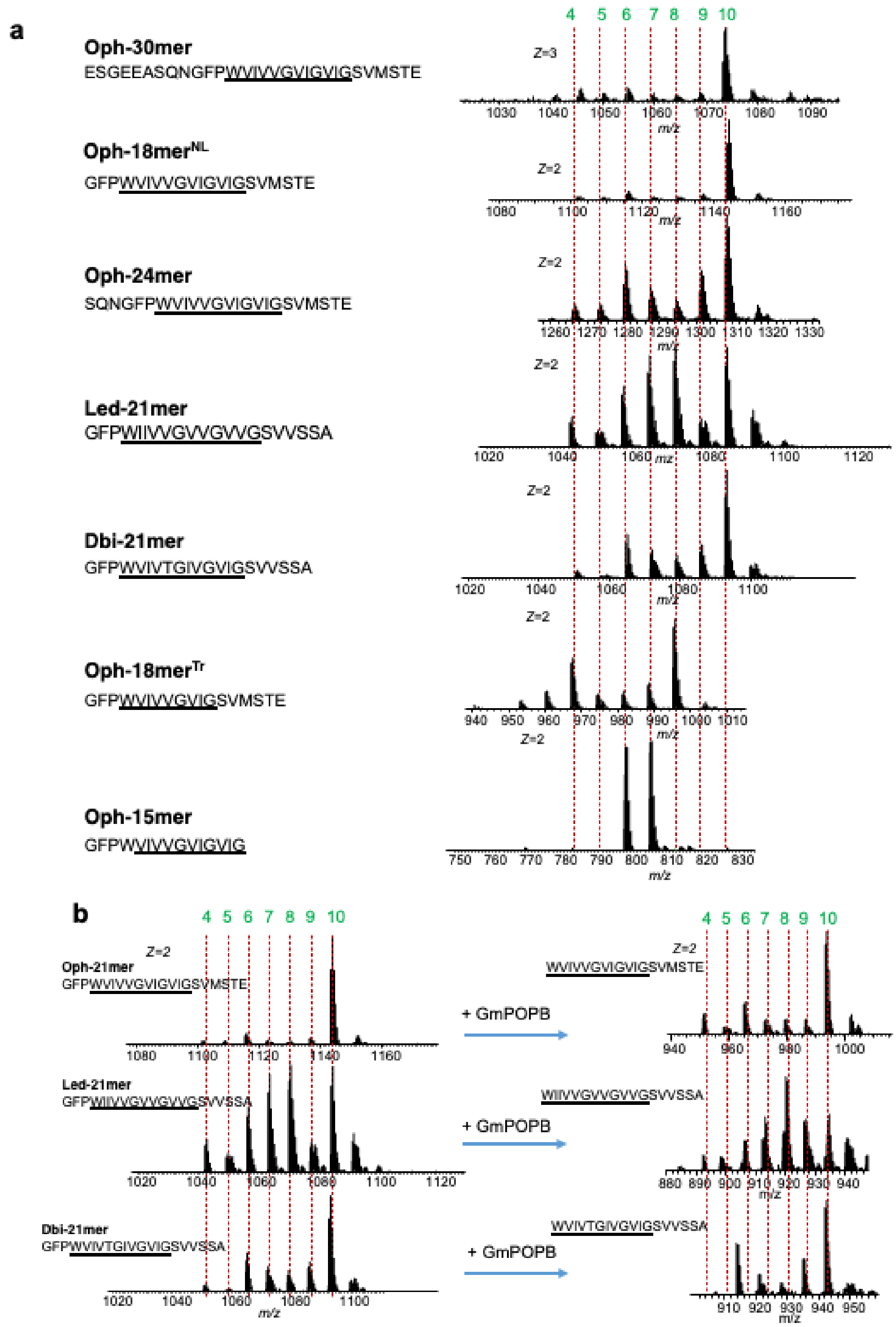

**Figure 2-figure supplement 2**. **LC-MS/MS spectra of OphP peptide substrates and products.** (**a-e**) Peptide substrates obtained by cleaving OphMA-variants with TEV protease: Oph-21mer, Led-21mer, Dbi-21mer, Oph-18mer$^{NL}$, Oph-18mer$^{Tr}$, Oph-24mer and Oph-15mer. (**f-l**) Linear peptide products obtained by incubation of the peptide substrates with OphP: linear 15mer and 12mer derived from Oph-21mer (**f-g**), linear 12mer and 15mer derived from Led-21mer (**h-i**), 15mer derived from Dbi-21mer (**j**), linear 15mer and 18mer derived from Oph-24mer (**k-l**). (**m-o**) Linear omphalotin A, lentinulin A and dendrothelin A intermediates. Methylated residues are highlighted in green color. b-ions and y-ions are shown in blue color. Mass tolerance between observed masses and the theoretical masses was set at 10 ppm. This difference is indicated in parentheses. Mass precision was set at 4 decimals.

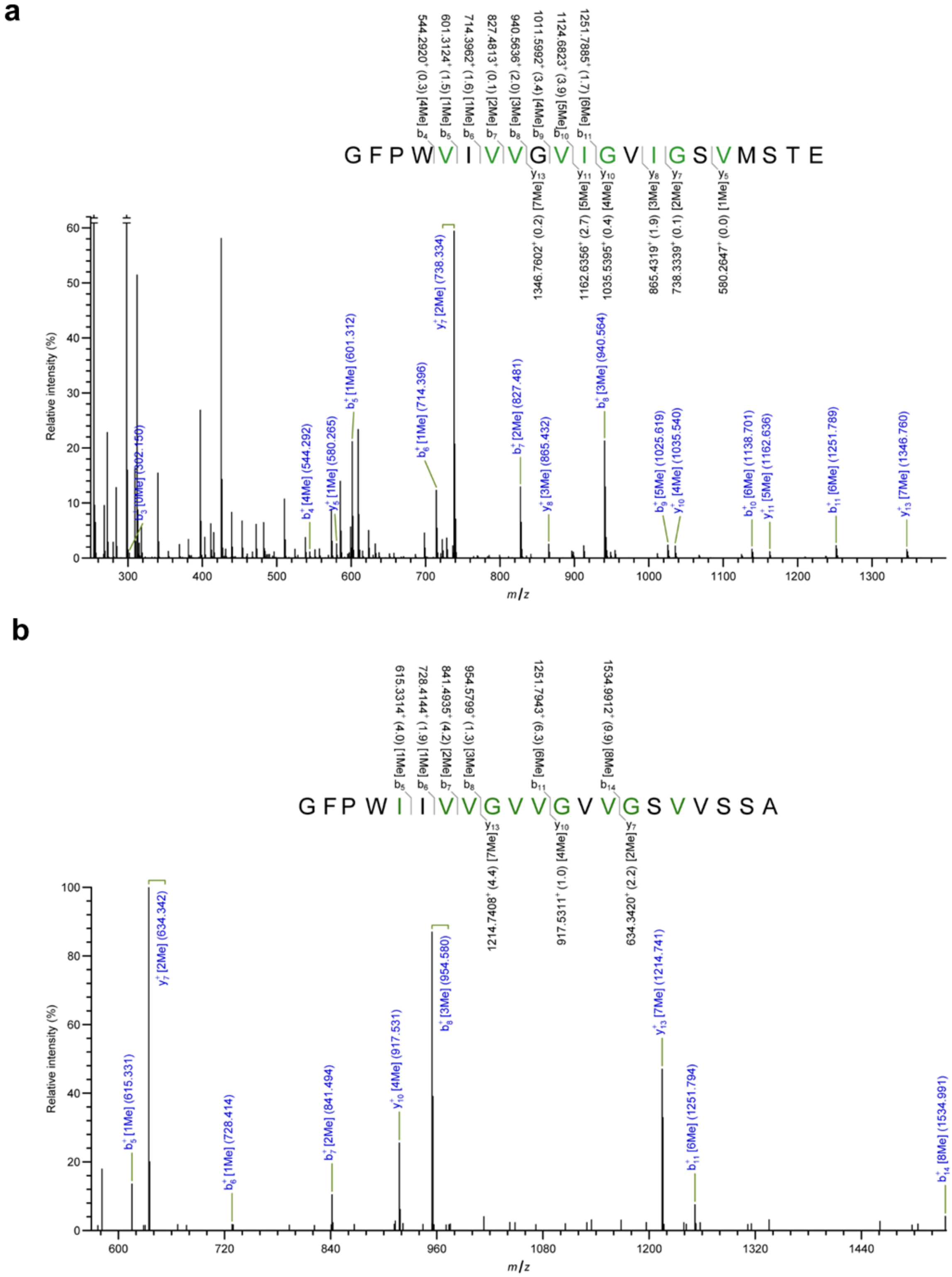

a
GFPWVIVVGVIGVIGSVMSTE
b4 544.2920+ (0.3) [4Me]
b5 601.3124+ (1.5) [1Me]
b6 714.3962+ (1.6) [1Me]
b7 827.4813+ (0.1) [2Me]
b8 940.5636+ (2.0) [3Me]
b9 1011.5992+ (3.4) [4Me]
b10 1124.6823+ (3.9) [5Me]
b11 1251.7885+ (1.7) [6Me]
y13 1346.7602+ (0.2) [7Me]
y11 1162.6356+ (2.7) [5Me]
y10 1035.5395+ (0.4) [4Me]
y8 865.4319+ (1.9) [3Me]
y7 738.3339+ (0.1) [2Me]
y5 580.2647+ (0.0) [1Me]
Relative intensity (%)
m/z
b3+ [0Me] (302.150)
b4+ [4Me] (544.292)
y5+ [1Me] (580.265)
b5+ [1Me] (601.312)
b6+ [1Me] (714.396)
y7+ [2Me] (738.334)
b7+ [2Me] (827.481)
y8+ [3Me] (865.432)
b8+ [3Me] (940.564)
b9+ [5Me] (1025.619)
y10+ [4Me] (1035.540)
b10+ [6Me] (1138.701)
y11+ [5Me] (1162.636)
b11+ [6Me] (1251.789)
y13+ [7Me] (1346.760)
b
GFPWIIVVGVVGVVGSVVSSA
b5 615.3314+ (4.0) [1Me]
b6 728.4144+ (1.9) [1Me]
b7 841.4935+ (4.2) [2Me]
b8 954.5799+ (1.3) [3Me]
b11 1251.7943+ (6.3) [6Me]
b14 1534.9912+ (6.9) [8Me]
y13 1214.7408+ (4.4) [7Me]
y10 917.5311+ (1.0) [4Me]
y7 634.3420+ (2.2) [2Me]
Relative intensity (%)
m/z
b5+ [1Me] (615.331)
y7+ [2Me] (634.342)
b6+ [1Me] (728.414)
b7+ [2Me] (841.494)
y10+ [4Me] (917.531)
b8+ [3Me] (954.580)
y13+ [7Me] (1214.741)
b11+ [6Me] (1251.794)
b14+ [8Me] (1534.991)

c

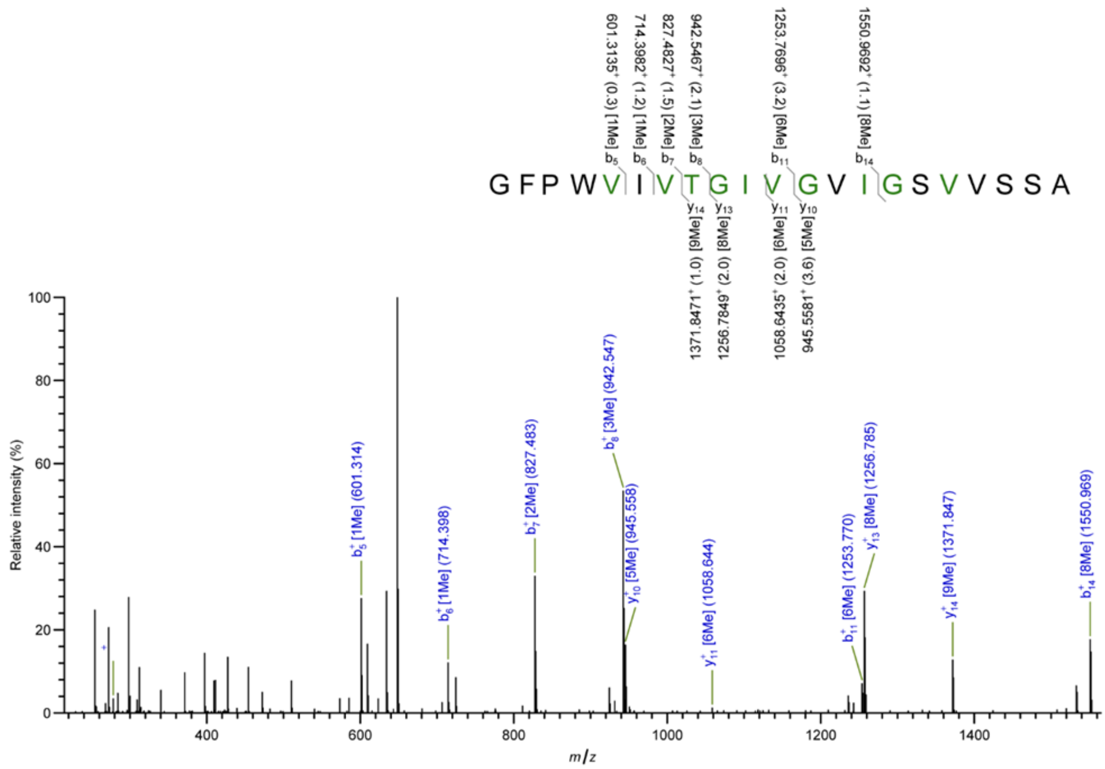

G F P W V I V T G I V G V I G S V V S S A
601.3135+ (0.3) [1Me] b5
714.3982+ (1.2) [1Me] b6
827.4827+ (1.5) [2Me] b7
942.5467+ (2.1) [3Me] b8
1253.7696+ (3.2) [6Me] b11
1550.9692+ (1.1) [8Me] b14
y14 1371.8471+ (1.0) [9Me]
y13 1256.7849+ (2.0) [8Me]
y11 1058.6435+ (2.0) [6Me]
y10 945.5581+ (3.6) [5Me]
b5+ [1Me] (601.314)
b6+ [1Me] (714.398)
b7+ [2Me] (827.483)
b8+ [3Me] (942.547)
y10+ [5Me] (945.558)
y11+ [6Me] (1058.644)
b11+ [6Me] (1253.770)
y13+ [8Me] (1256.785)
y14+ [9Me] (1371.847)
b14+ [8Me] (1550.969)
Relative intensity (%)
m/z


d

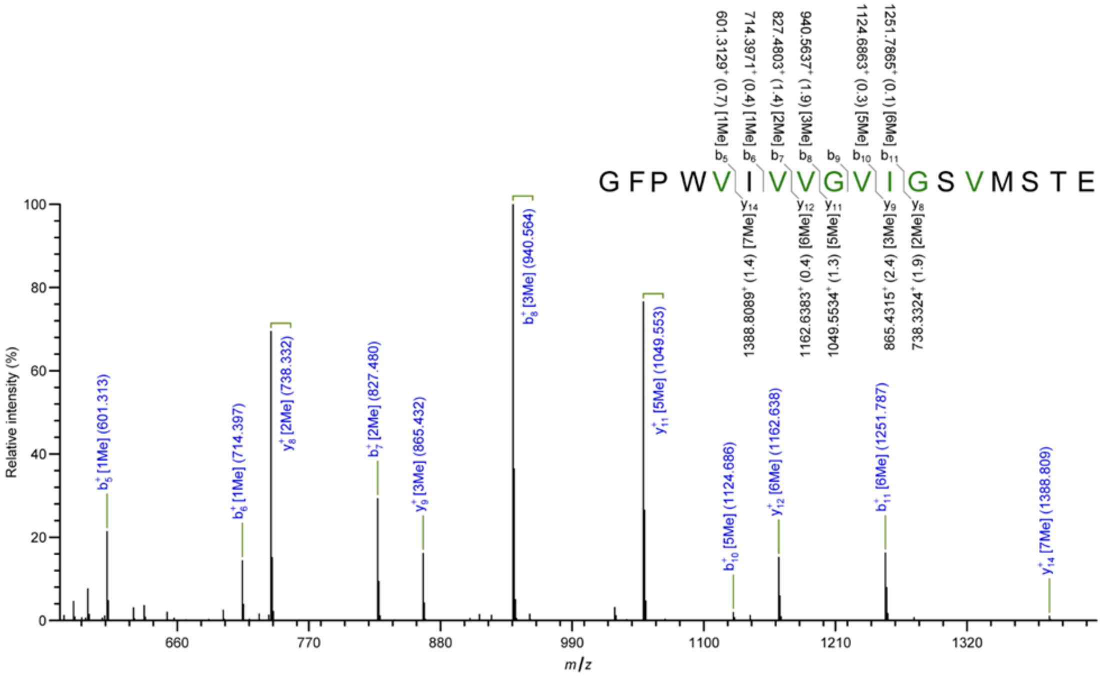

G F P W V I V V G V I G S V M S T E
601.3129+ (0.7) [1Me] b5
714.3971+ (0.4) [1Me] b6
827.4803+ (1.4) [2Me] b7
940.5637+ (1.9) [3Me] b8
b9
1124.6863+ (0.3) [5Me] b10
1251.7865+ (0.1) [6Me] b11
y14 1388.8089+ (1.4) [7Me]
y12 1162.6383+ (0.4) [6Me]
y11 1049.5534+ (1.3) [5Me]
y9 865.4315+ (2.4) [3Me]
y8 738.3324+ (1.9) [2Me]
b5+ [1Me] (601.313)
b6+ [1Me] (714.397)
y8+ [2Me] (738.332)
b7+ [2Me] (827.480)
y9+ [3Me] (865.432)
b8+ [3Me] (940.564)
y11+ [5Me] (1049.553)
b10+ [5Me] (1124.686)
y12+ [6Me] (1162.638)
b11+ [6Me] (1251.787)
y14+ [7Me] (1388.809)
Relative intensity (%)
m/z

**e**

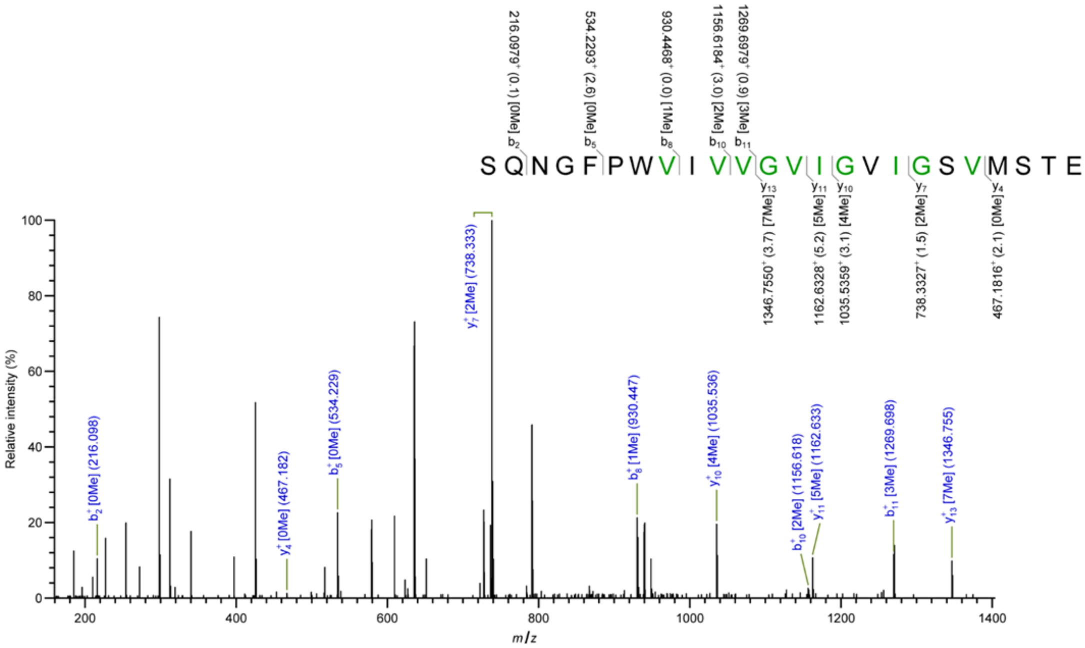

S Q N G F P W V I V V G V I G V I G S V M S T E
216.0979+ (0.1) [0Me]
534.2293+ (2.6) [0Me]
930.4468+ (0.0) [1Me]
1156.6184+ (3.0) [2Me]
1269.6979+ (0.9) [3Me]
1346.7550+ (3.7) [7Me]
1162.6328+ (5.2) [5Me]
1035.5359+ (3.1) [4Me]
738.3327+ (1.5) [2Me]
467.1816+ (2.1) [0Me]
b2+ [0Me] (216.098)
y4+ [0Me] (467.182)
b5+ [0Me] (534.229)
y7+ [2Me] (738.333)
b8+ [1Me] (930.447)
y10+ [4Me] (1035.536)
b10+ [2Me] (1156.618)
y11+ [5Me] (1162.633)
b11+ [3Me] (1269.698)
y13+ [7Me] (1346.755)
Relative intensity (%)
m/z

**f**

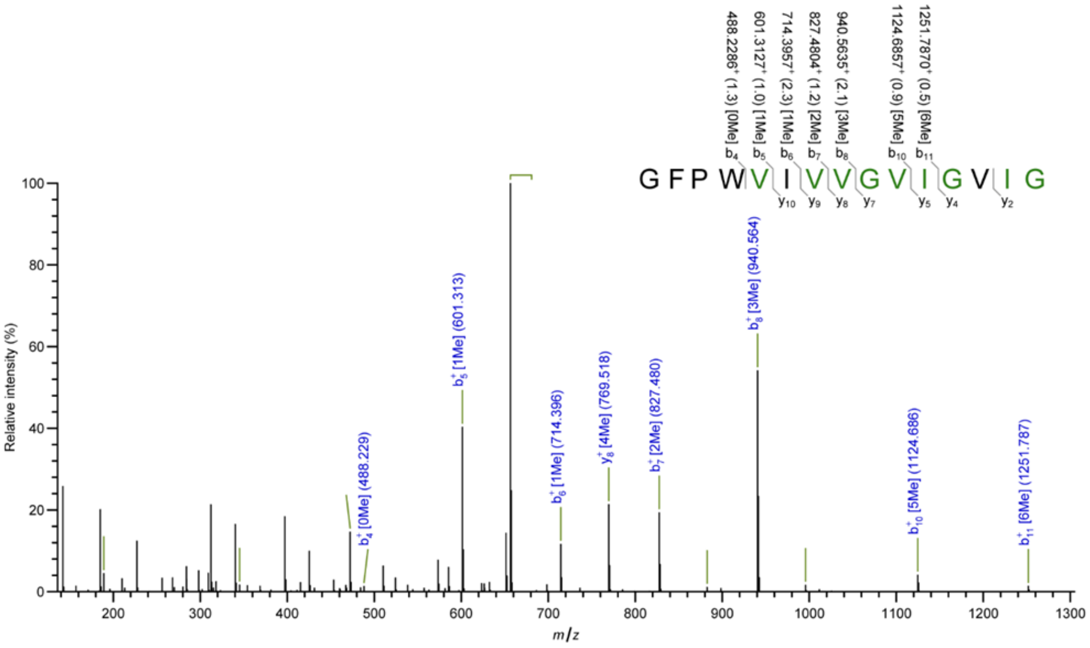
GFPWVIVVGVIGVIG
488.2286+ (1.3) [0Me] b4
601.3127+ (1.0) [1Me] b5
714.3957+ (2.3) [1Me] b6
827.4804+ (1.2) [2Me] b7
940.5635+ (2.1) [3Me] b8
1124.6857+ (0.9) [5Me] b10
1251.7870+ (0.5) [6Me] b11
y10 y9 y8 y7 y5 y4 y2
Relative intensity (%)
m/z
b4+ [0Me] (488.229)
b5+ [1Me] (601.313)
b6+ [1Me] (714.396)
y8+ [4Me] (769.518)
b7+ [2Me] (827.480)
b8+ [3Me] (940.564)
b10+ [5Me] (1124.686)
b11+ [6Me] (1251.787)

**g**

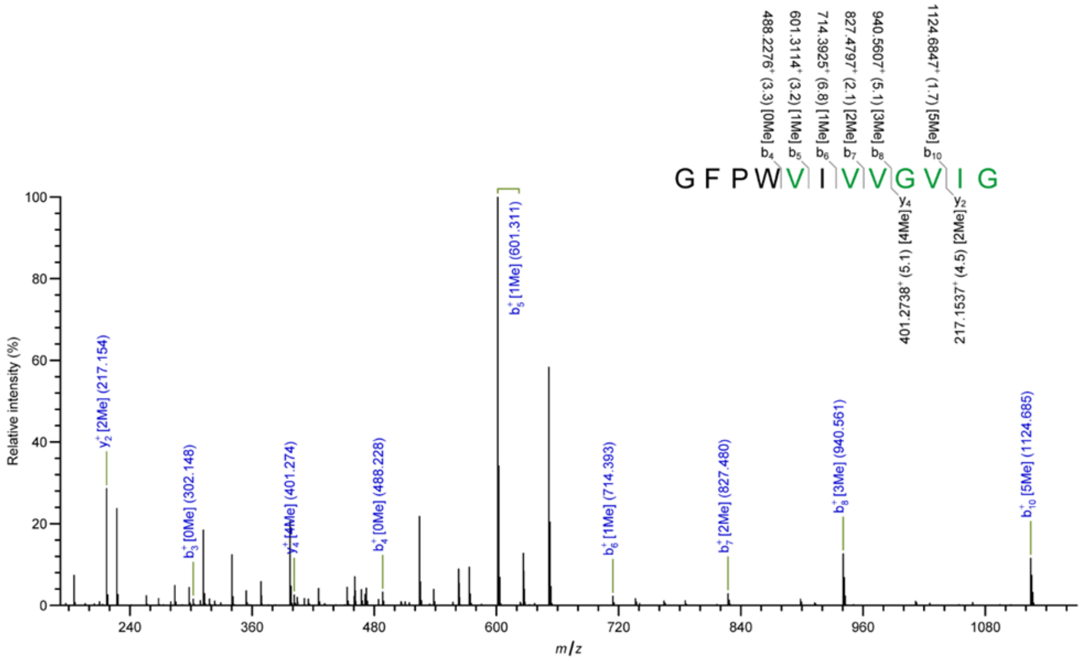
GFPWVIVVGVIG
488.2276+ (3.3) [0Me] b4
601.3114+ (3.2) [1Me] b5
714.3925+ (6.8) [1Me] b6
827.4797+ (2.1) [2Me] b7
940.5607+ (5.1) [3Me] b8
1124.6847+ (1.7) [5Me] b10
y4 [4Me] (5.1) 401.2738+
y2 [2Me] (4.5) 217.1537+
Relative intensity (%)
m/z
y2+ [2Me] (217.154)
b3+ [0Me] (302.148)
y4+ [4Me] (401.274)
b4+ [0Me] (488.228)
b5+ [1Me] (601.311)
b6+ [1Me] (714.393)
b7+ [2Me] (827.480)
b8+ [3Me] (940.561)
b10+ [5Me] (1124.685)

**h**

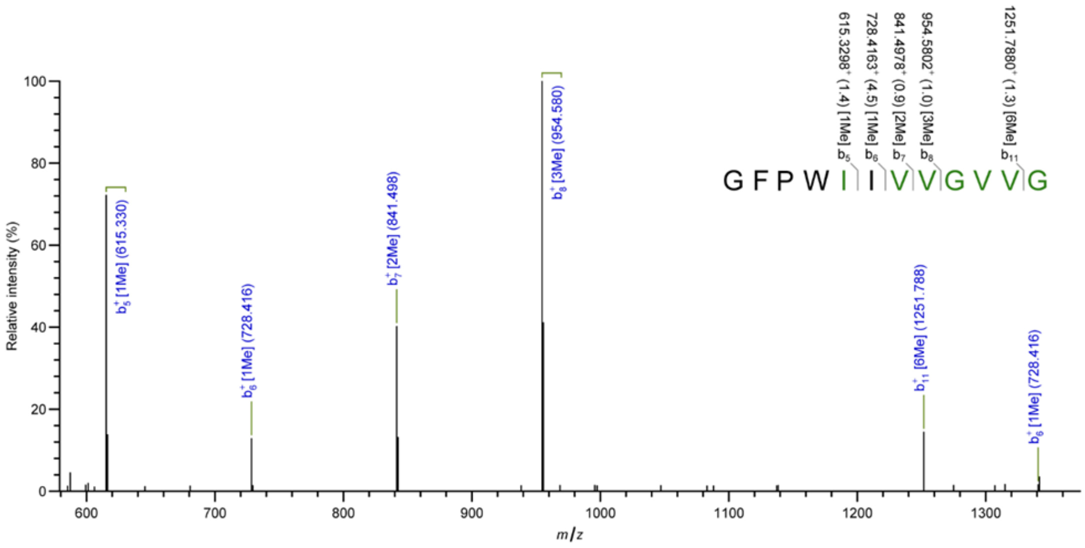
GFPWIIVVGVVG
b5 b6 b7 b8 b11
615.3298+ (1.4) [1Me]
728.4163+ (4.5) [1Me]
841.4978+ (0.9) [2Me]
954.5802+ (1.0) [3Me]
1251.7880+ (1.3) [6Me]
b5+ [1Me] (615.330)
b6+ [1Me] (728.416)
b7+ [2Me] (841.498)
b8+ [3Me] (954.580)
b11+ [6Me] (1251.788)
b6+ [1Me] (728.416)
Relative intensity (%)
m/z

**i**

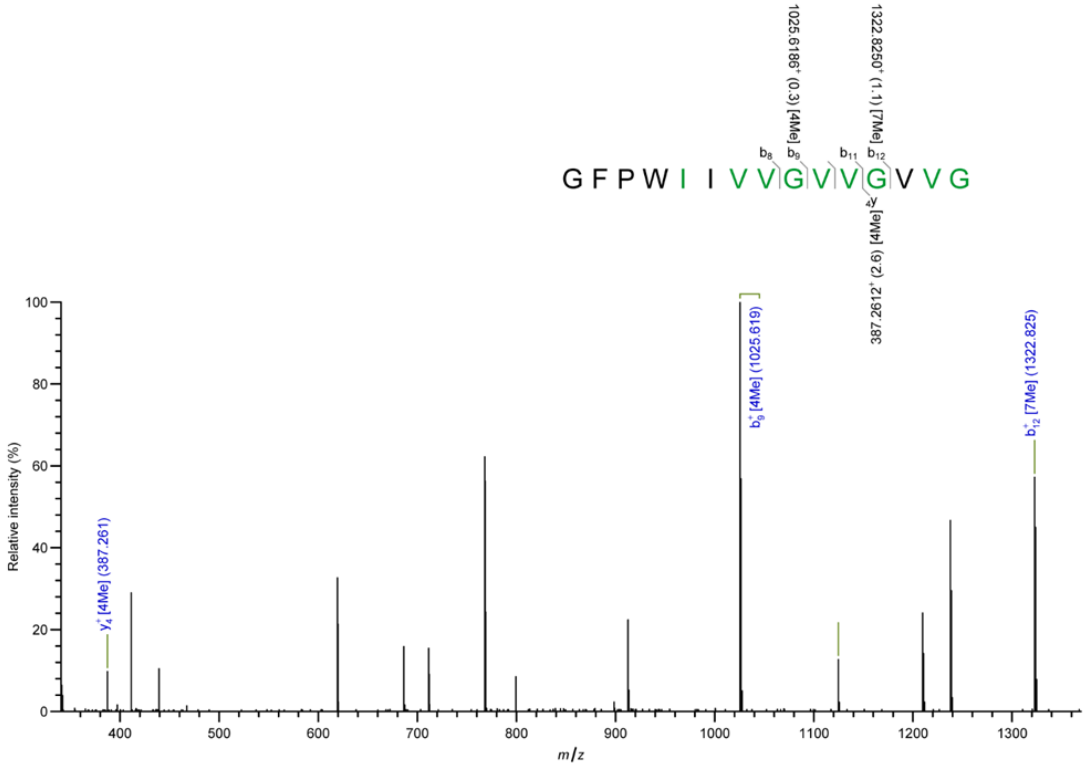
GFPWIIVVGVVGVVG
b8 b9 b11 b12
y4
1025.6186+ (0.3) [4Me]
1322.8250+ (1.1) [7Me]
387.2612+ (2.6) [4Me]
y4+ [4Me] (387.261)
b9+ [4Me] (1025.619)
b12+ [7Me] (1322.825)
Relative intensity (%)
m/z

**j**

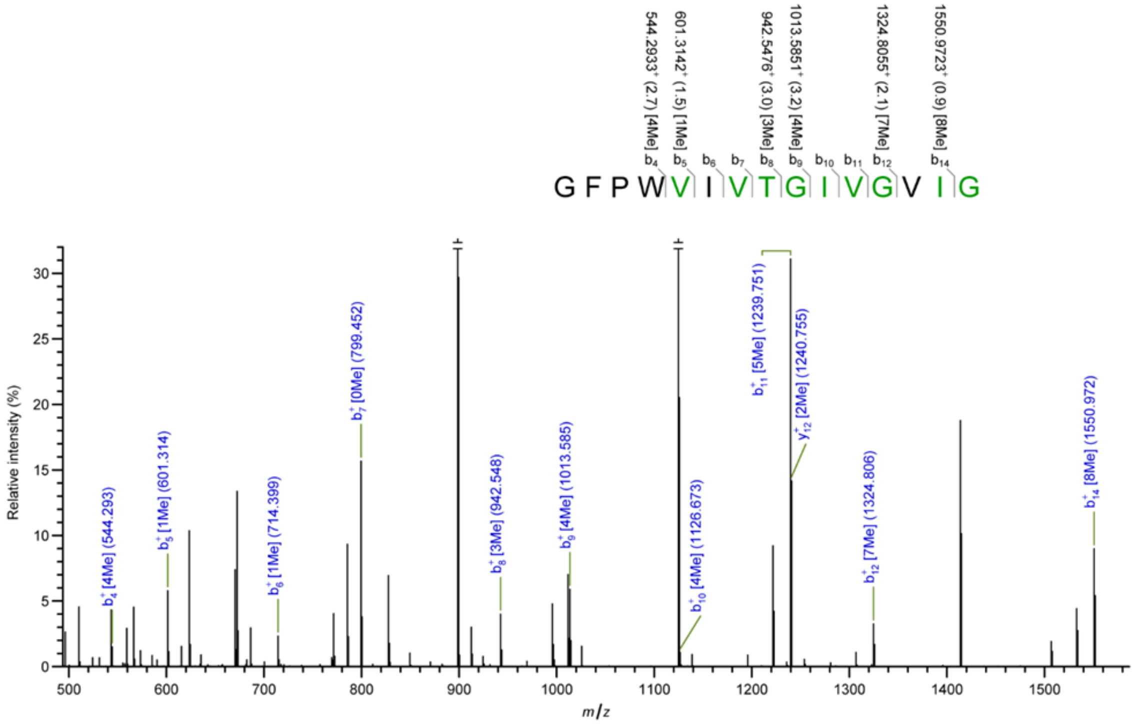

544.2933+ (2.7) [4Me]
601.3142+ (1.5) [1Me]
942.5476+ (3.0) [3Me]
1013.5851+ (3.2) [4Me]
1324.8055+ (2.1) [7Me]
1550.9723+ (0.9) [8Me]
G F P W V I V T G I V G V I G
Relative intensity (%)
m/z
b4+ [4Me] (544.293)
b5+ [1Me] (601.314)
b6+ [1Me] (714.399)
b7+ [0Me] (799.452)
b8+ [3Me] (942.548)
b9+ [4Me] (1013.585)
b10+ [4Me] (1126.673)
b11+ [5Me] (1239.751)
y12+ [2Me] (1240.755)
b12+ [7Me] (1324.806)
b14+ [8Me] (1550.972)


**k**

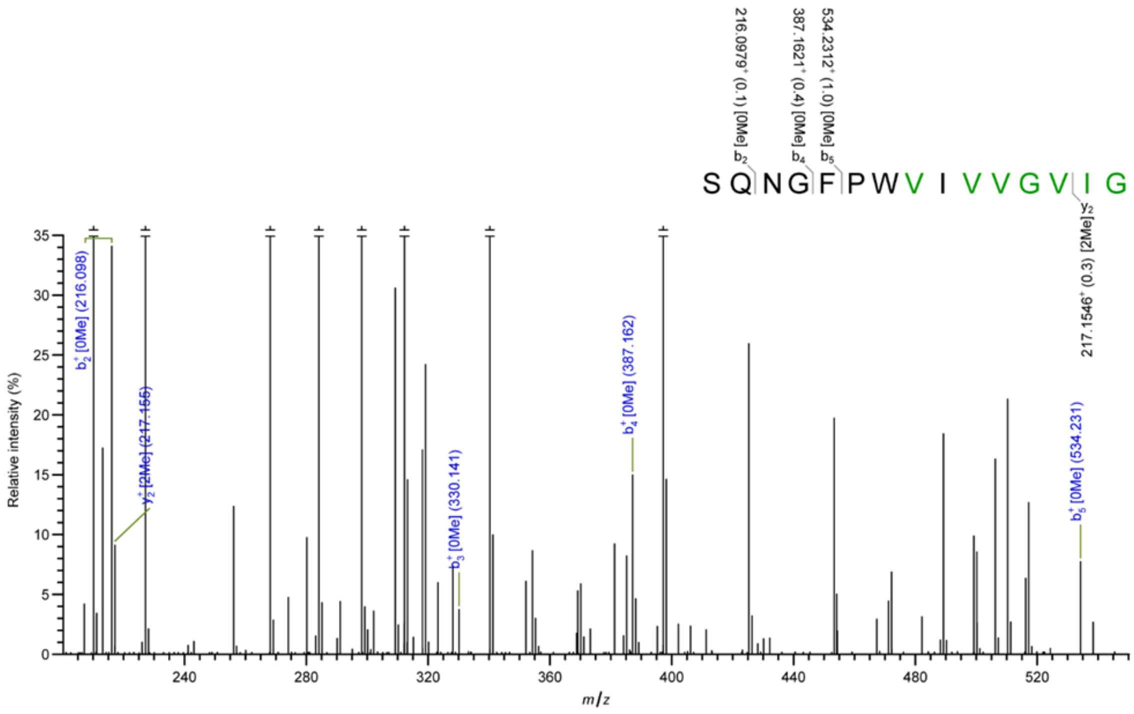

216.0979+ (0.1) [0Me]
387.1621+ (0.4) [0Me]
534.2312+ (1.0) [0Me]
S Q N G F P W V I V V G V I G
217.1546+ (0.3) [2Me]
Relative intensity (%)
m/z
b2+ [0Me] (216.098)
y2+ [2Me] (217.155)
b3+ [0Me] (330.141)
b4+ [0Me] (387.162)
b5+ [0Me] (534.231)

l

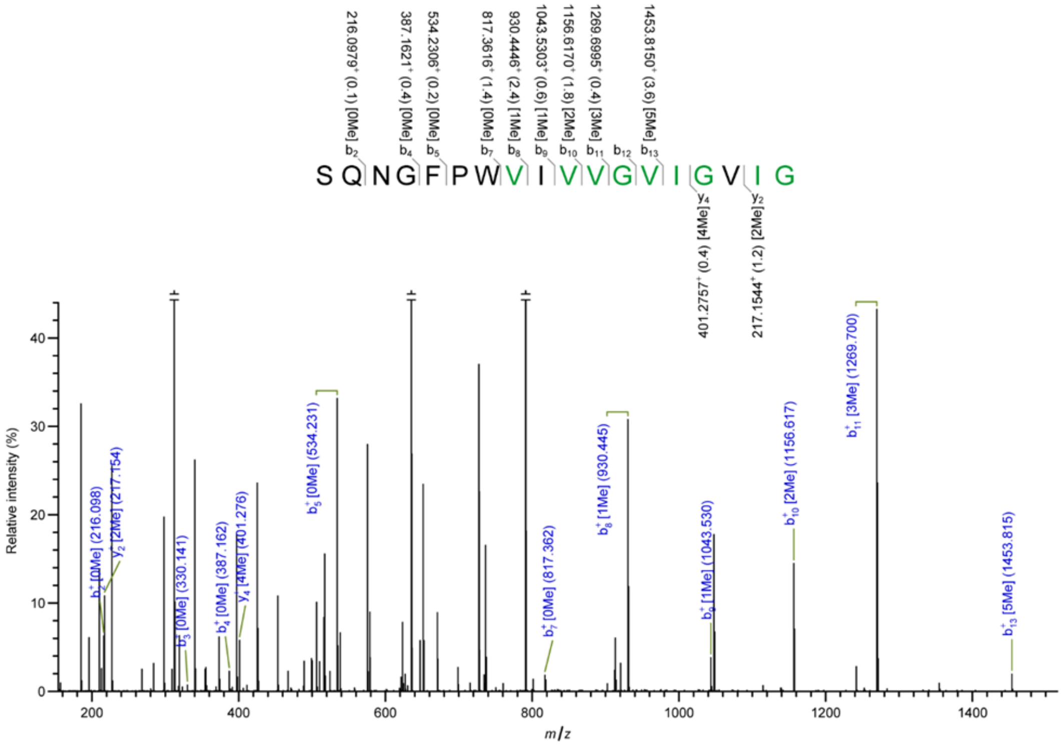

216.0979+ (0.1) [0Me]
387.1621+ (0.4) [0Me]
534.2306+ (0.2) [0Me]
817.3616+ (1.4) [0Me]
930.4446+ (2.4) [1Me]
1043.5303+ (0.6) [1Me]
1156.6170+ (1.8) [2Me]
1269.6995+ (0.4) [3Me]
1453.8150+ (3.6) [5Me]
b2 b4 b5 b7 b8 b9 b10 b11 b12 b13
S Q N G F P W V I V V G V I G V I G
y4 y2
401.2757+ (0.4) [4Me]
217.1544+ (1.2) [2Me]
b2+ [0Me] (216.098)
y2+ [2Me] (217.154)
b3+ [0Me] (330.141)
b4+ [0Me] (387.162)
y4+ [4Me] (401.276)
b5+ [0Me] (534.231)
b7+ [0Me] (817.362)
b8+ [1Me] (930.445)
b9+ [1Me] (1043.530)
b10+ [2Me] (1156.617)
b11+ [3Me] (1269.700)
b13+ [5Me] (1453.815)
Relative intensity (%)
m/z


m

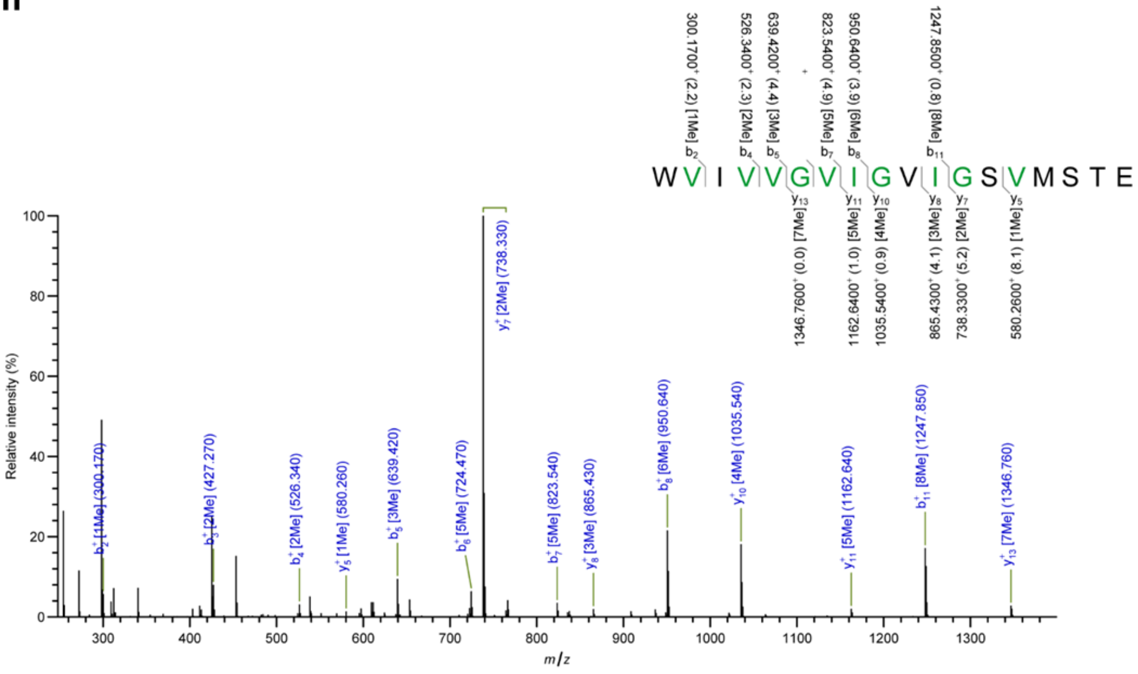

300.1700+ (2.2) [1Me]
526.3400+ (2.3) [2Me]
639.4200+ (4.4) [3Me]
823.5400+ (4.9) [5Me]
950.6400+ (3.9) [6Me]
1247.8500+ (0.8) [8Me]
b2 b4 b5 b7 b8 b11
W V I V V G V I G V I G S V M S T E
y13 y11 y10 y8 y7 y5
1346.7600+ (0.0) [7Me]
1162.6400+ (1.0) [5Me]
1035.5400+ (0.9) [4Me]
865.4300+ (4.1) [3Me]
738.3300+ (5.2) [2Me]
580.2600+ (8.1) [1Me]
y7+ [2Me] (738.330)
b2+ [1Me] (300.170)
b3+ [2Me] (427.270)
b4+ [2Me] (526.340)
y5+ [1Me] (580.260)
b5+ [3Me] (639.420)
b6+ [5Me] (724.470)
b7+ [5Me] (823.540)
y8+ [3Me] (865.430)
b8+ [6Me] (950.640)
y10+ [4Me] (1035.540)
y11+ [5Me] (1162.640)
b11+ [8Me] (1247.850)
y13+ [7Me] (1346.760)
Relative intensity (%)
m/z

**n**

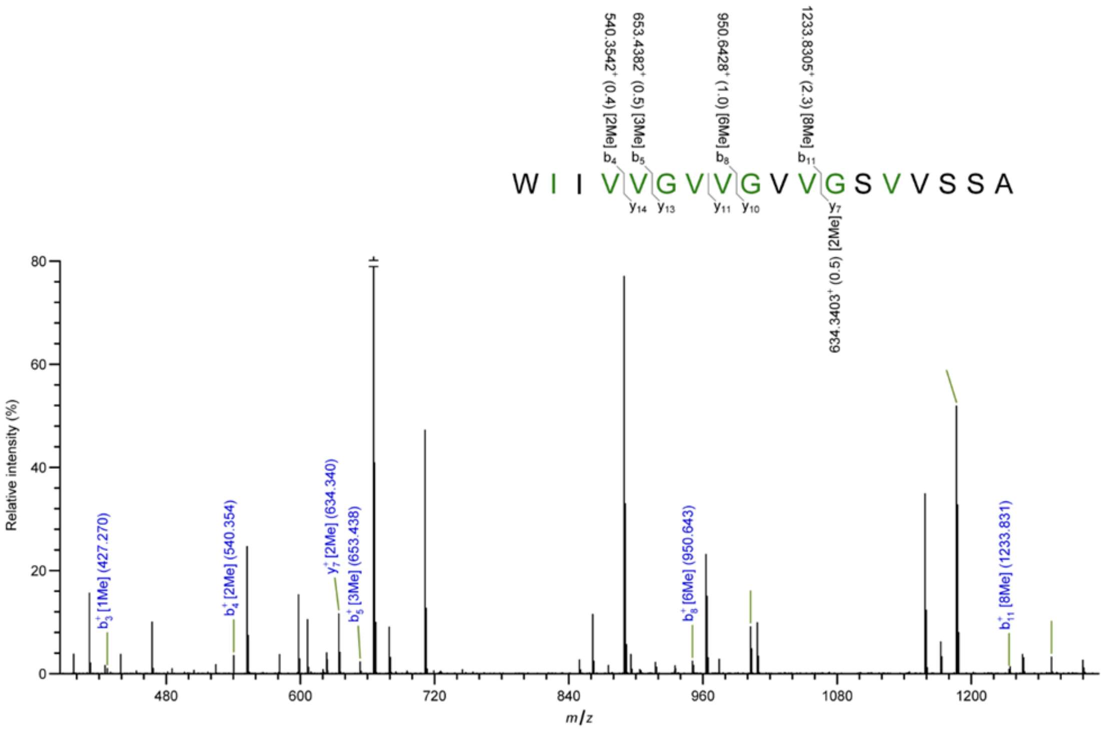

540.3542+ (0.4) [2Me] b4
653.4382+ (0.5) [3Me] b5
950.6428+ (1.0) [6Me] b8
1233.8305+ (2.3) [8Me] b11
W I I V V G V V G V V G S V V S S A
y14 y13 y11 y10 y7
634.3403+ (0.5) [2Me]
b3+ [1Me] (427.270)
b4+ [2Me] (540.354)
y7+ [2Me] (634.340)
b5+ [3Me] (653.438)
b8+ [6Me] (950.643)
b11+ [8Me] (1233.831)
Relative intensity (%)
m/z


**o**

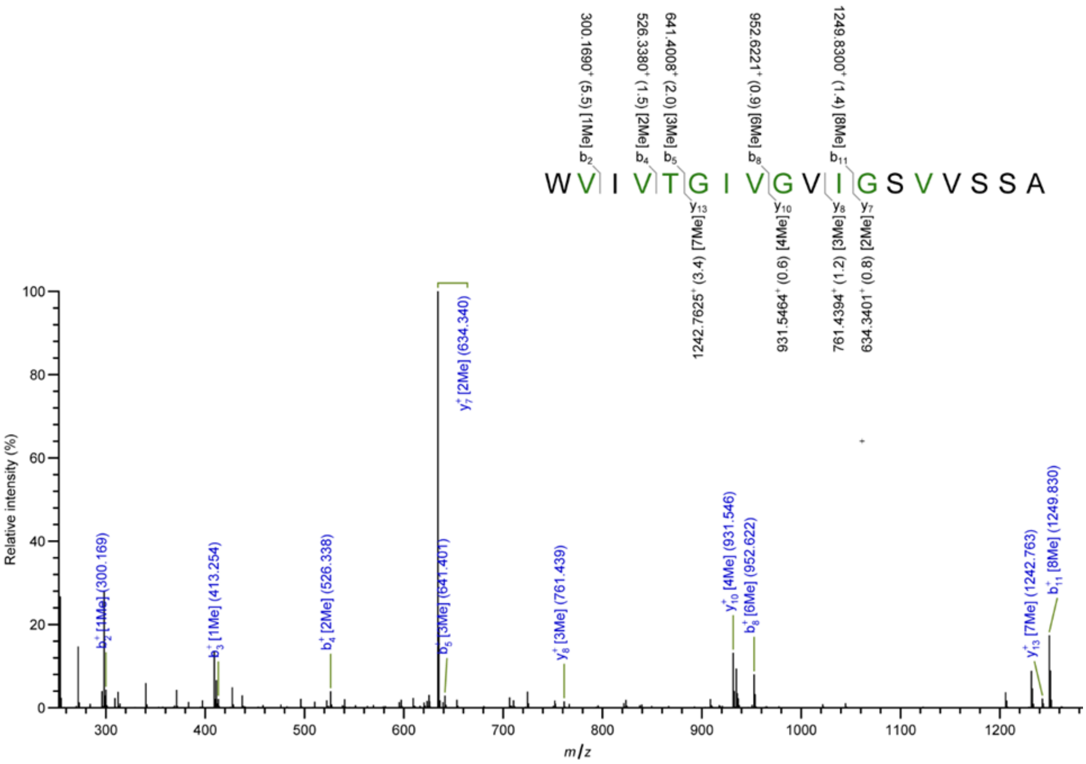

300.1690+ (5.5) [1Me] b2
526.3380+ (1.5) [2Me] b4
641.4008+ (2.0) [3Me] b5
952.6221+ (0.9) [6Me] b8
1249.8300+ (1.4) [8Me] b11
W V I V T G I V G V I G S V V S S A
y13 y10 y8 y7
1242.7625+ (3.4) [7Me]
931.5464+ (0.6) [4Me]
761.4394+ (1.2) [3Me]
634.3401+ (0.8) [2Me]
y7+ [2Me] (634.340)
b2+ [1Me] (300.169)
b3+ [1Me] (413.254)
b4+ [2Me] (526.338)
b5+ [3Me] (641.401)
y8+ [3Me] (761.439)
y10+ [4Me] (931.546)
b8+ [6Me] (952.622)
y13+ [7Me] (1242.763)
b11+ [8Me] (1249.830)
Relative intensity (%)
m/z

**Figure 2-figure supplement 3. HPLC-MS of macrocyclic OphP peptide products.** Extracted ion chromatograms (EICs) of peptide products derived from Led-21mer and Dbi-21mer by the action of OphP. Methylated residues are highlighted in green. Macrocyclization is indicated by a bracket below the peptide sequence.

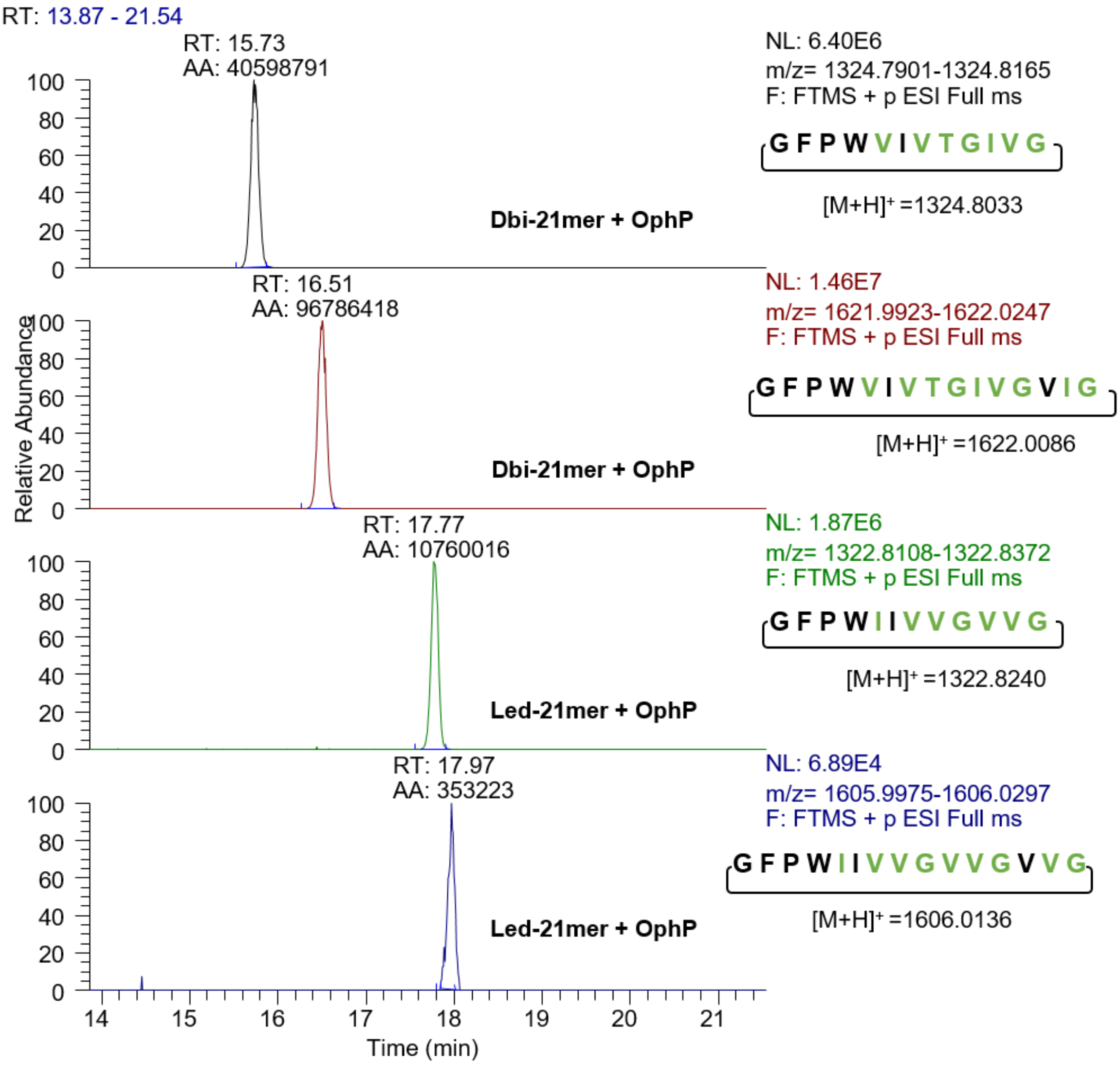

**Figure 2-figure supplement 4. Comparison of endopeptidase/macrocyclase activities towards the Oph-15mer peptide. (a)** GmPOPB compared to OphP and LedP activities. (**b**) ProAlanase activity. The enzymatic activity was assessed by determining the UV absorbance at 280 nm.

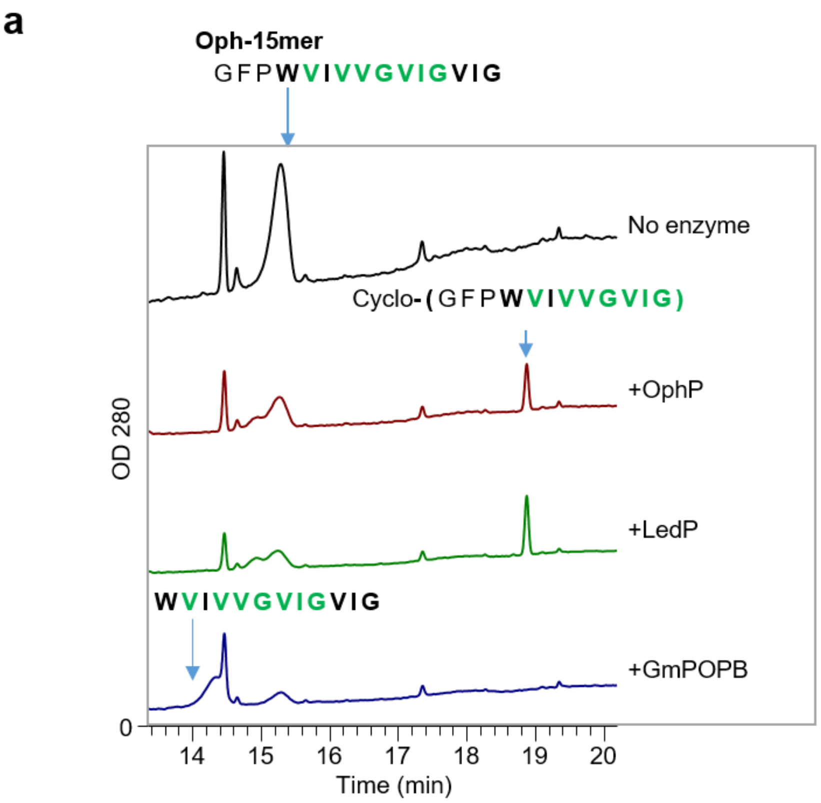


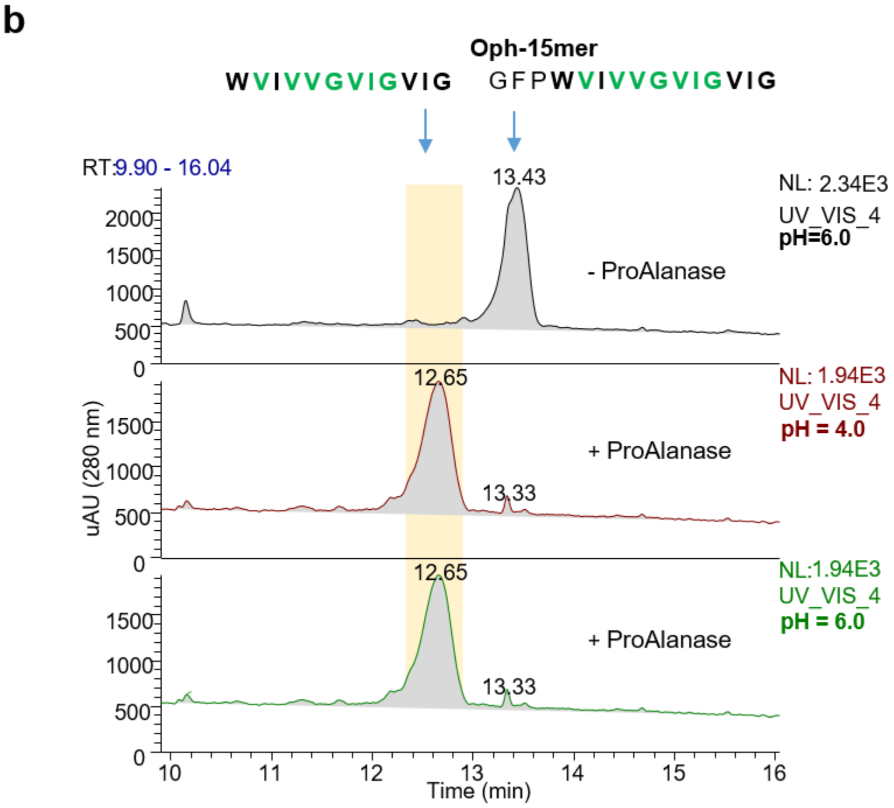

**Figure 2-figure supplement 5. GmPOPB and OphP activity on synthetic peptides PHA1 and AMA1.** PHA1 and AMA1 are known to be processed by GmPOPB protease (Czekster et al., 2017; Luo et al., 2014). The graphs show the HPLC-UV-VIS chromatograms with optical density (OD) at 280nm. The synthetic peptides, PHA1 and AMA1 were stored at 1 mg/ml in phosphate buffer at pH 6.0. For the reaction, 20 µM substrate was used with 1 µM of OphP or GmPOPB in the standard assay buffer [50 mM HEPES pH 6.0 + 10 mM DTT]. The reaction mixture (50 µl) was incubated at 30°C, 600 rpm, overnight. The reaction was quenched by the addition of 50 µl methanol. After centrifugation at the highest speed at 4°C, 5 µl supernatant was used for HPLC-MS/MS analysis. No cyclic peptides were observed for PHA1. The peak around 12 min, marked with an asterisk, is unspecific as it does not correspond to any product. The amanitin and phallacidin core peptides are underlined. The products are indicated by a blue arrow.

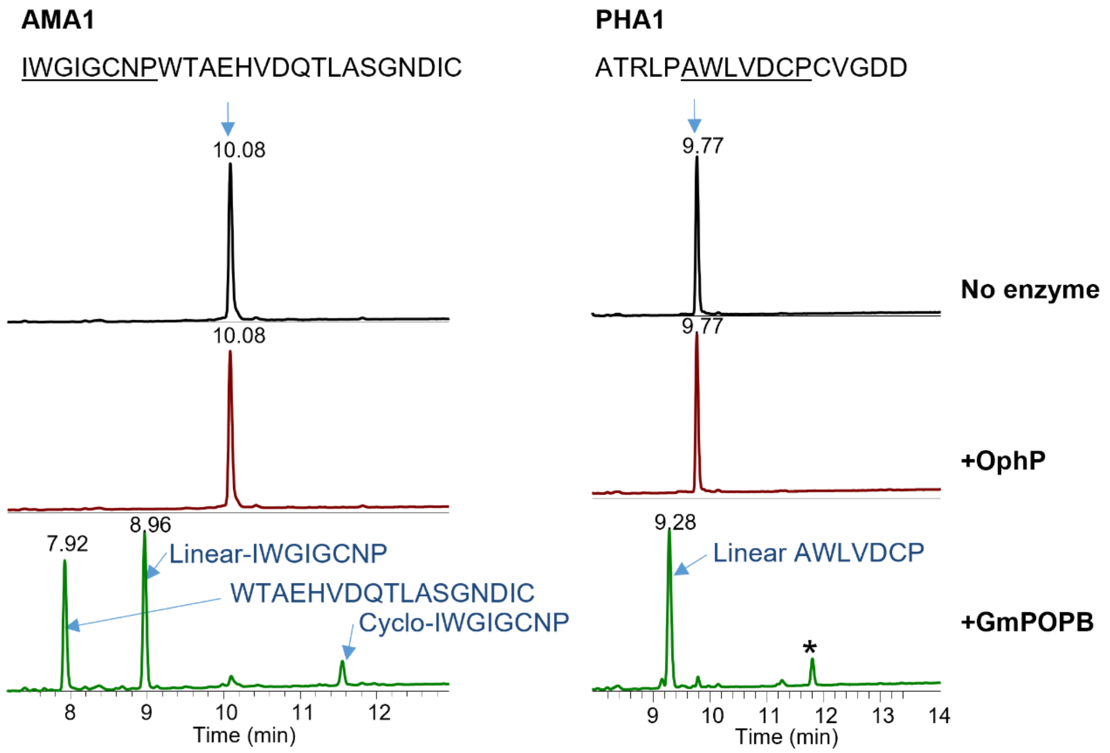

**Figure 2-figure supplement 6. Susceptibility of *in vivo* and *in vitro* production of omphalotin-related peptides towards protease inhibitors.** (**a**) *In vivo* production of omphalotin A in *P. pastoris* co-expressing *ophMA* and *ophP*, in the presence of different protease inhibitors. Omphalotin A concentrations were quantified by integrating the Extracted Ion Chromatogram (EIC) peak area of HPLC-MS measurements of cell lysates of triplicate cultures. (**b**) *In vitro* production of 6- and 7-fold methylated linear and macrocyclic peptide product species with 100 µM total Oph-15mer peptide substrate and 10 µM OphP concentration, in presence of different protease inhibitors. NOTE: The batch of Oph-15mer used for these experiments is the same as used for Figure 3bc and Figure 3-figure supplement 2. The reaction was performed in HEPES buffer at pH 7.0 and stopped after 30 minutes. Product species were quantified using the EIC peak area of HPLC-MS measurements. (**c**) *In vitro* production of 6- and 7-fold methylated linear and macrocyclic peptide product species with 100µM total Oph-15mer peptide substrate and 10µM OphP concentration, in presence of ZPP at two different concentrations. The reaction was performed in HEPES buffer at pH 7.0 and stopped after 30 minutes. Peptide species were quantified by integrating the EIC peak area of the HPLC-MS measurements.

**a**

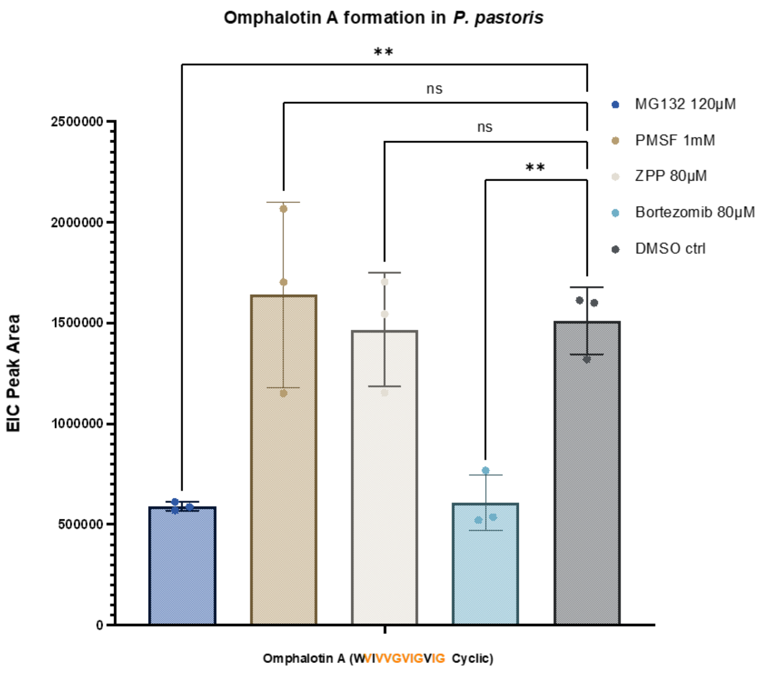

**b**

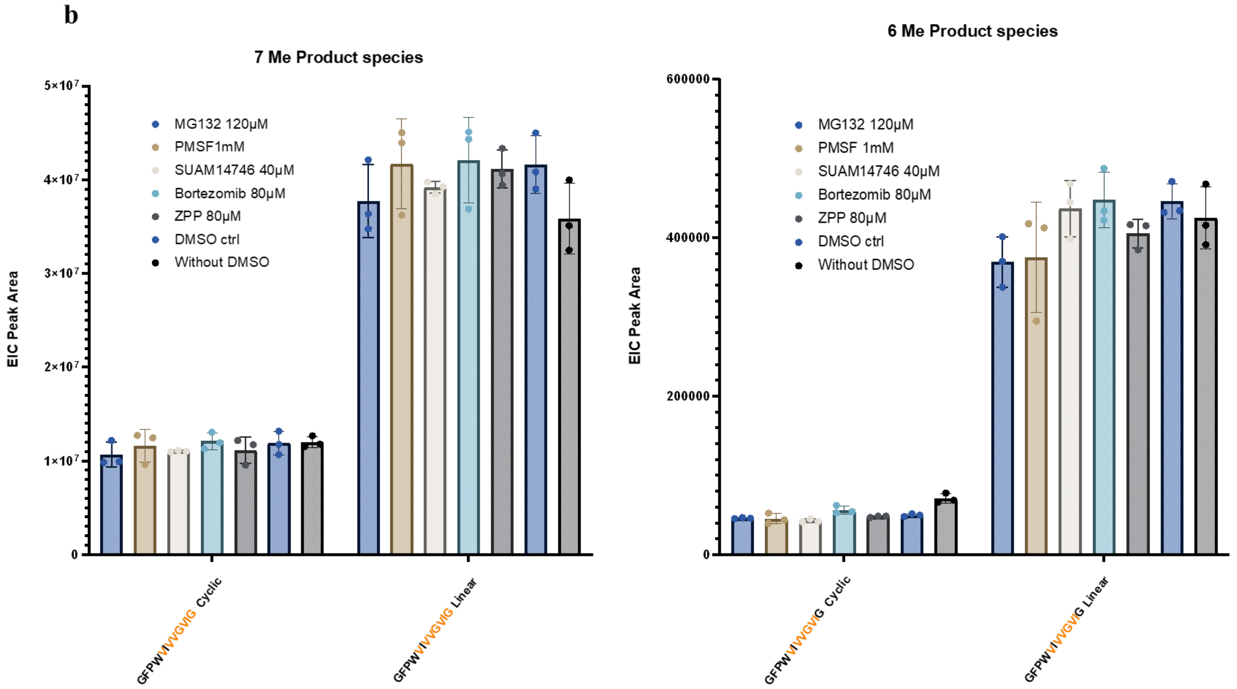
7 Me Product species
EIC Peak Area
5×10⁷
4×10⁷
3×10⁷
2×10⁷
1×10⁷
0
MG132 120μM
PMSF1mM
SUAM14746 40μM
Bortezomib 80μM
ZPP 80μM
DMSO ctrl
Without DMSO
GFPWIVVGVIG Cyclic
GFPWIVVGVIG Linear
6 Me Product species
EIC Peak Area
600000
400000
200000
0
MG132 120μM
PMSF 1mM
SUAM14746 40μM
Bortezomib 80μM
ZPP 80μM
DMSO ctrl
Without DMSO
GFPWIVVGVIG Cyclic
GFPWIVVGVIG Linear

**c**

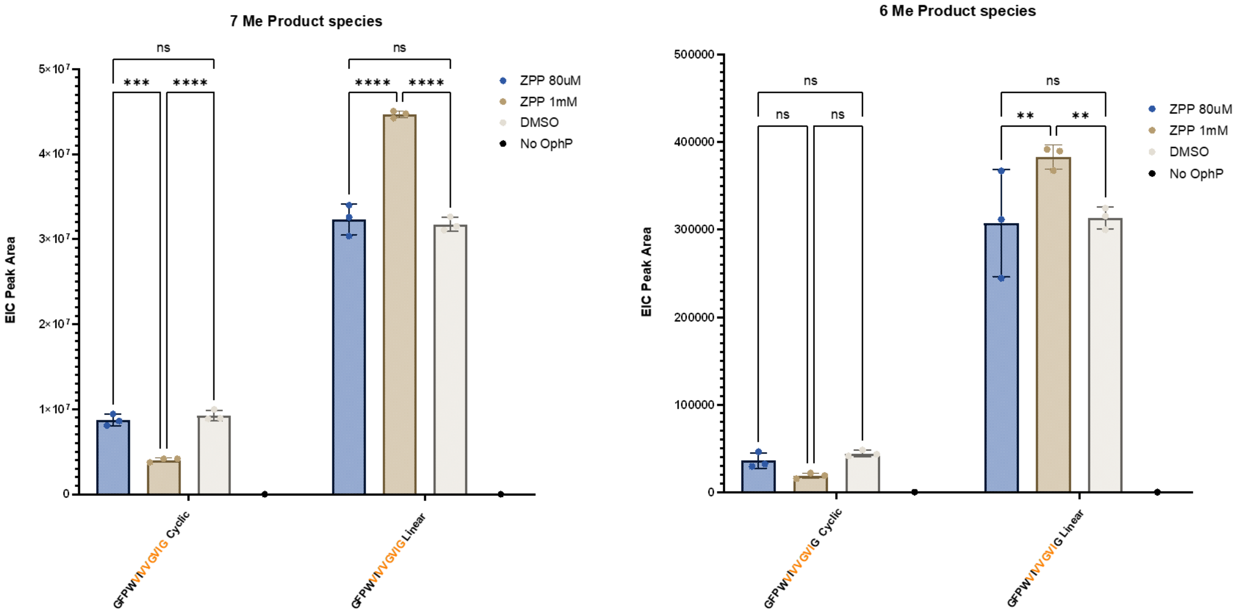
7 Me Product species
EIC Peak Area
5×10⁷
4×10⁷
3×10⁷
2×10⁷
1×10⁷
0
ns
***
****
ns
****
****
ZPP 80uM
ZPP 1mM
DMSO
No OphP
GFPWIVVGVIG Cyclic
GFPWIVVGVIG Linear
6 Me Product species
EIC Peak Area
500000
400000
300000
200000
100000
0
ns
ns
ns
ns
**
**
ZPP 80uM
ZPP 1mM
DMSO
No OphP
GFPWIVVGVIG Cyclic
GFPWIVVGVIG Linear

**Figure 2-figure supplement 7. *In vivo* production of omphalotin A in *P. pastoris* and *E. coli.*** Production of omphalotin A in *P. pastoris* was performed as described previously (Matabaro et al., 2021). In *E. coli*, production of omphalotin A was attempted by co expression of *ophMA* and *ophP* under the control of a T7 promoter. *E. coli* was grown in TB medium until $OD_{600}$ 1.5 and expression was induced by addition of 0.2 mM IPTG. The induced cultures were incubated for two days at 25 °C. The bacterial cells were lysed using 0.1 mm glass beads in a FastPrep device and omphalotin A was extracted using a 1:1 mixture of ethyl acetate and n-hexane. The organic phase was evaporated, the pellets were redissolved in 100 µl of methanol and 10 µl of the solution was subjected to HPLC-MS/MS analysis. The graph shows the extracted ion chromatogram (EIC) peak area of omphalotin A extracted from *P. pastoris* and *E. coli* adjusted to the microbial biomass (wet weight) inferred from the optical density at 600 nm.

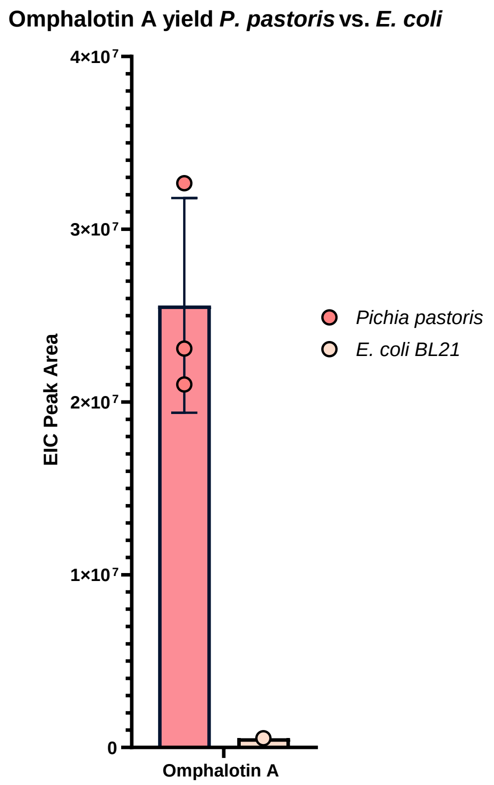
