## Supplementary material for "Molecular insight into the enzymatic macrocyclization of multiply backbone N-methylated peptides": Fig3SOI

**Figure 3-figure supplement 1. Preference of OphP for the highly methylated substrates.** The batch of Oph-15mer used for these experiments was not *in vitro* methylated and, thus, contained a quasi-equimolar ratio of 6- and 7-fold methylated peptide species. The reaction was conducted in the standard buffer [50 mM HEPES pH 7.0, 10 mM DTT] at 30°C, overnight**.** NOTE: The batch of Oph-15mer peptide substrate used for these experiments was the same as used for Figure 3a. In contrast to the batch used for Figure 3bc and Figure 3-Supplement 2, this batch was not *in vitro* methylated and, thus, contained a different ratio of 6- and 7-fold methylated peptide species. (**a**) RP-HPLC UV absorption profile of 100 µM of Oph-15mer with varying concentrations of OphP (0, 1, 2, 5, 10, 20 µM). Linear products resulting from the peptidase activity of OphP can be detected but overlapped with the starting material. 6-fold methylated cyclo(GFPW$^{me}$VI$^{me}$V$^{me}$V$^{me}$G$^{me}$V$^{me}$IG) can only be detected when high concentrations of OphP are used. Dashed lines show the retention times of the various peptide species. (**b**) Mass spectrometric analysis showing ion chromatograms of the consumption of Oph-15mer mixture (100 µM) when subjected to different concentrations of OphP. The 7-fold methylated Oph-15mer is depleted at high concentrations of OphP (10 and 20 µM). After the depletion of this species, OphP started to process the 6-fold methylated species.

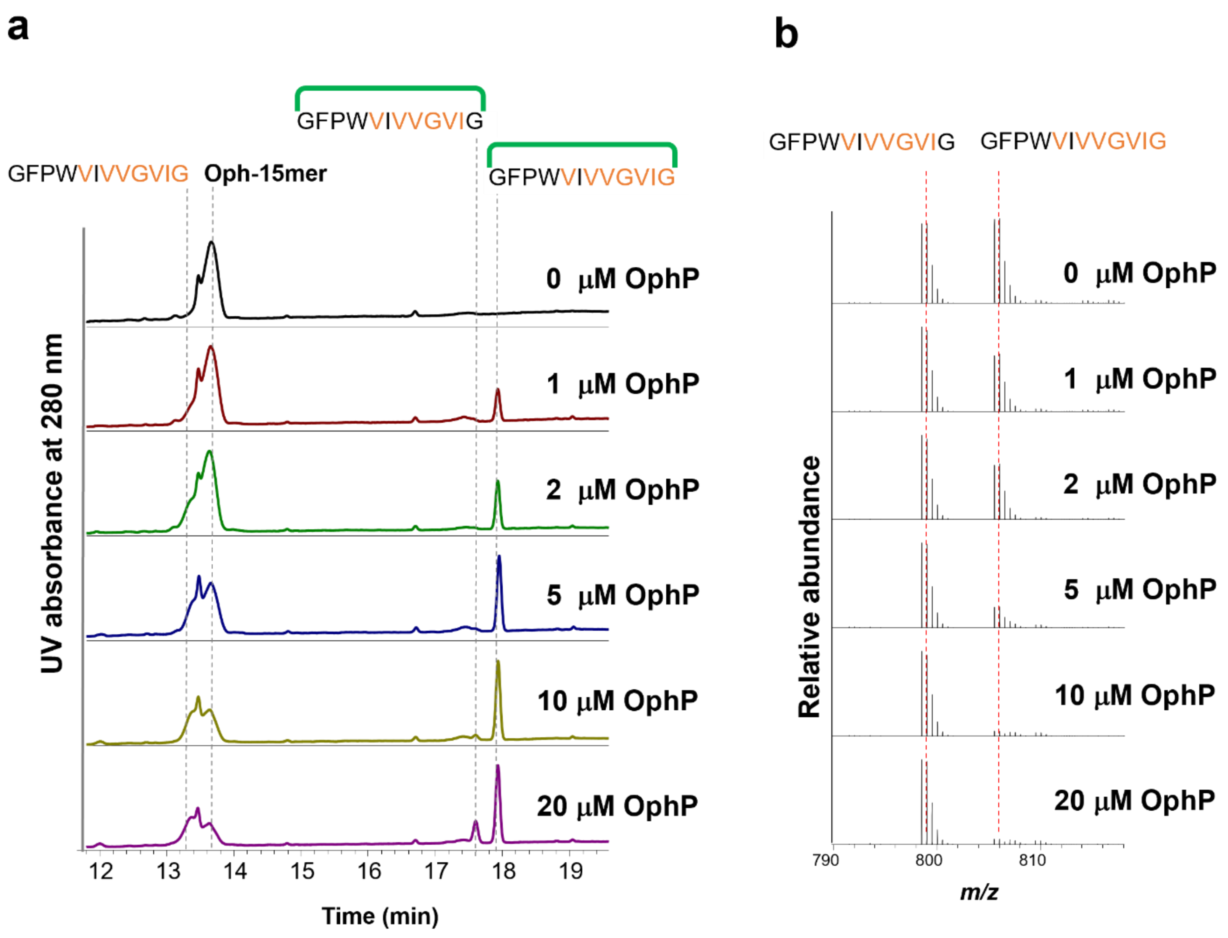

**Figure 3-figure supplement 2. Preferred production of 7-fold linear and cyclic peptide product from a mixture of 6- and 7fold Oph-15mer by OphP.** NOTE: The batch of Oph-15mer peptide substrate used for these experiments was also used for Figure 3, panels b and c. EIC peak area of Oph-15mer substrate and product species of 30 min *in vitro* reaction with 10µM OphP and different Oph-15mer substrate (mixture of 6Me and 7Me) concentrations in HEPES buffer at pH 7.0. (**a**) Comparison of 6- and 7-fold methylated substrate species at 100µM total substrate concentration without addition of OphP. (**b**) Comparison of 6- and 7-fold methylated product species at equimolar (90µM) substrate concentration and 10µM OphP concentration. (**c**) Comparison of 6- and 7-fold methylated product species at equimolar (9µM) substrate concentration and 10µM OphP concentration.

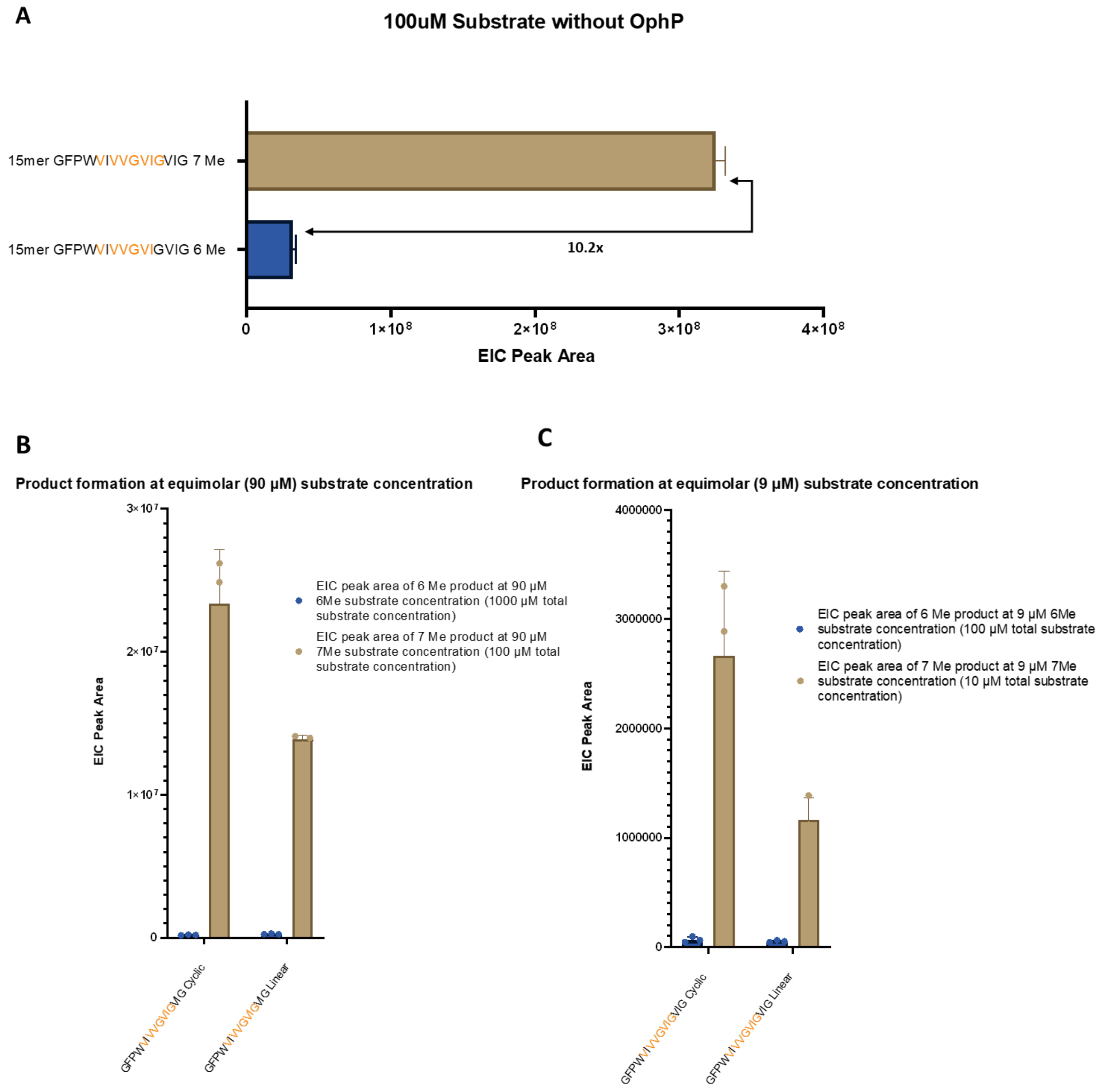
