## Supplementary material for "Molecular insight into the enzymatic macrocyclization of multiply backbone N-methylated peptides": Fig5SOI

**Figure 5-figure supplement 1. Crystal structures of OphP in complex with Oph-15mer.** $\sigma_A$-weighted 2mFo-DFc map (contoured at 1) for Oph-15mer (**a**) complexed with OphP S580A. Active site view of OphP: Oph-15mer (**b**) in the complex structure showing the interactions. Side chains of the active site residues are shown in sticks with carbon green, nitrogen blue, oxygen red and the methyl groups orange. Broken lines show hydrogen bonds.

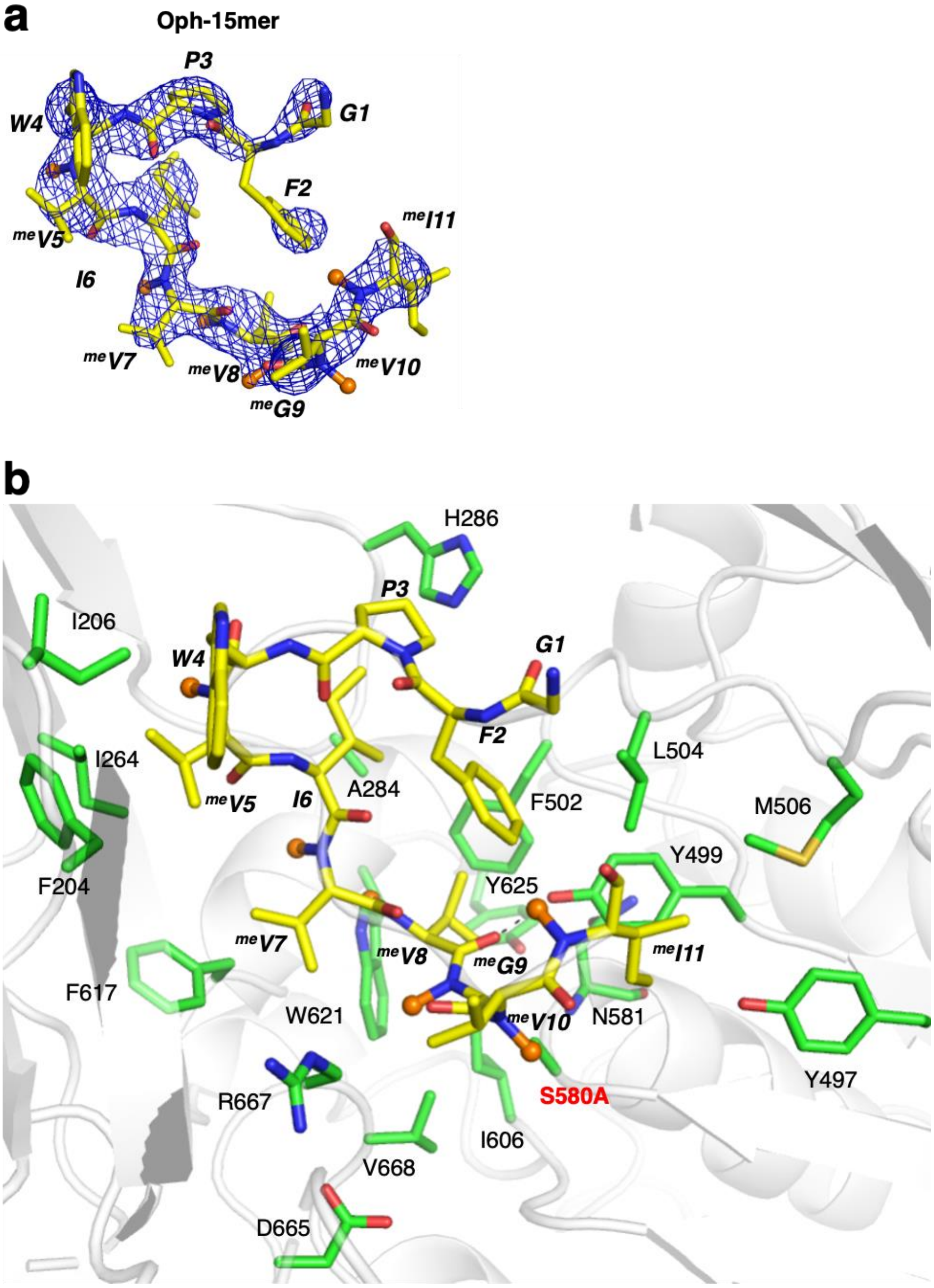

**Figure 5-figure supplement 2**. **Surface of the P1 pocket accommodating the serine acyl intermediate for PCY1 and OphP.** (**a**) The canonical P1 pocket of PCY1 can accommodate proline well at the cleavage P1 site. (**b**) Presence of covalently linked ZPP results in flipped Ile606. P1 pocket becomes larger to accommodate proline of ZPP. Flipped Ile606 clashes with Val584 with a 2.8 Å distance. (**c**) Surface showing clash between the proline of ZPP and Ile606. Cδ of I606 is only 2.3 Å away from Cβ of proline.

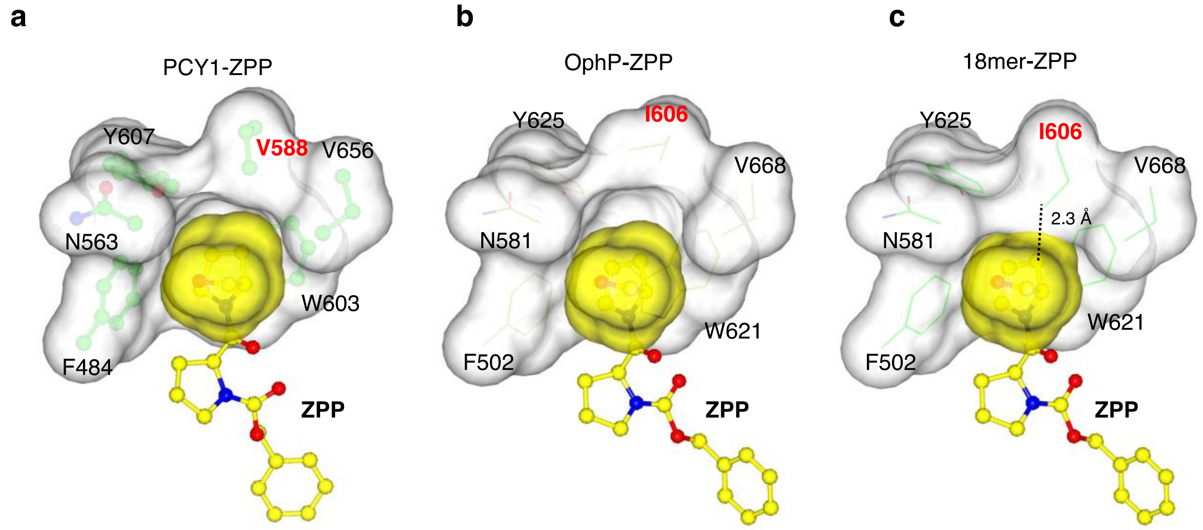
