## Supplementary material for "Molecular insight into the enzymatic macrocyclization of multiply backbone N-methylated peptides": Fig4SOI

**Figure 4-figure supplement 1. Multiple sequence alignment of OphP with post-proline cleaving prolyl oligopeptidases (POPs).** The sequence of OphP is used as a query and blasted (protein-protein BLAST) against the UniProtKB/SwissProt database. Sequences are further selected by removing all the dipeptidyl peptidase family enzyme sequences. Multiple sequence alignment was created with MUSCLE [15] and rendered as white and black with ESPript 3.0[16]. The hydrophobic pocket residues accommodating P1 proline are highlighted by underlying solid red triangles and catalytic triad residues are shown in solid purple triangles. Secondary structure elements of OphP are depicted above the aligned sequence. Black box with white character shows strict identity, bold black character similarity, and black frame similarity across groups. Names, UniProt ID and corresponding organisms are as follows: PCY1, R4P353, *Gypsophila vaccaria*; GmPOPB, H2E7Q8, *Galerina marginate*; GmPOPA, H2E7Q7, *Galerina marginate*; AbPOPB, E2JFG2, *Amanita bisporigera*; AbPOPA, E2JFG1, *Amanita bisporigera*; PfuPOP, Q51714 (PDB entry 5T88), *Pyrococcus furiosus*; AhPOP, Q06903, *Aeromonas hydrophila*; EmPOP1, P27028, *Elizabethkingia meningoseptica*; EmPOP2, P27195, *Elizabethkingia meningoseptica*; DbPOP, Q86AS5, *Dictyostelium discoideum*; MmPOP, Q9QUR6, *Mus musculus*; P48147, *Homo sapiens*; BtPOP, Q9XTA2, *Bos taurus*; SsPOP, P23687 (PDB entry 1QFS), *Sus scrofa*. Sequence of DbiP from *Dendrothele bispora* is retrieved from Joint Genome Institute website (name: gm1.24165_g, protein ID: 871165, location: scaffold_621:21283-24607). The sequence of LedP from *Lentinula* edodes is obtained from NCBI database (GAW09065).

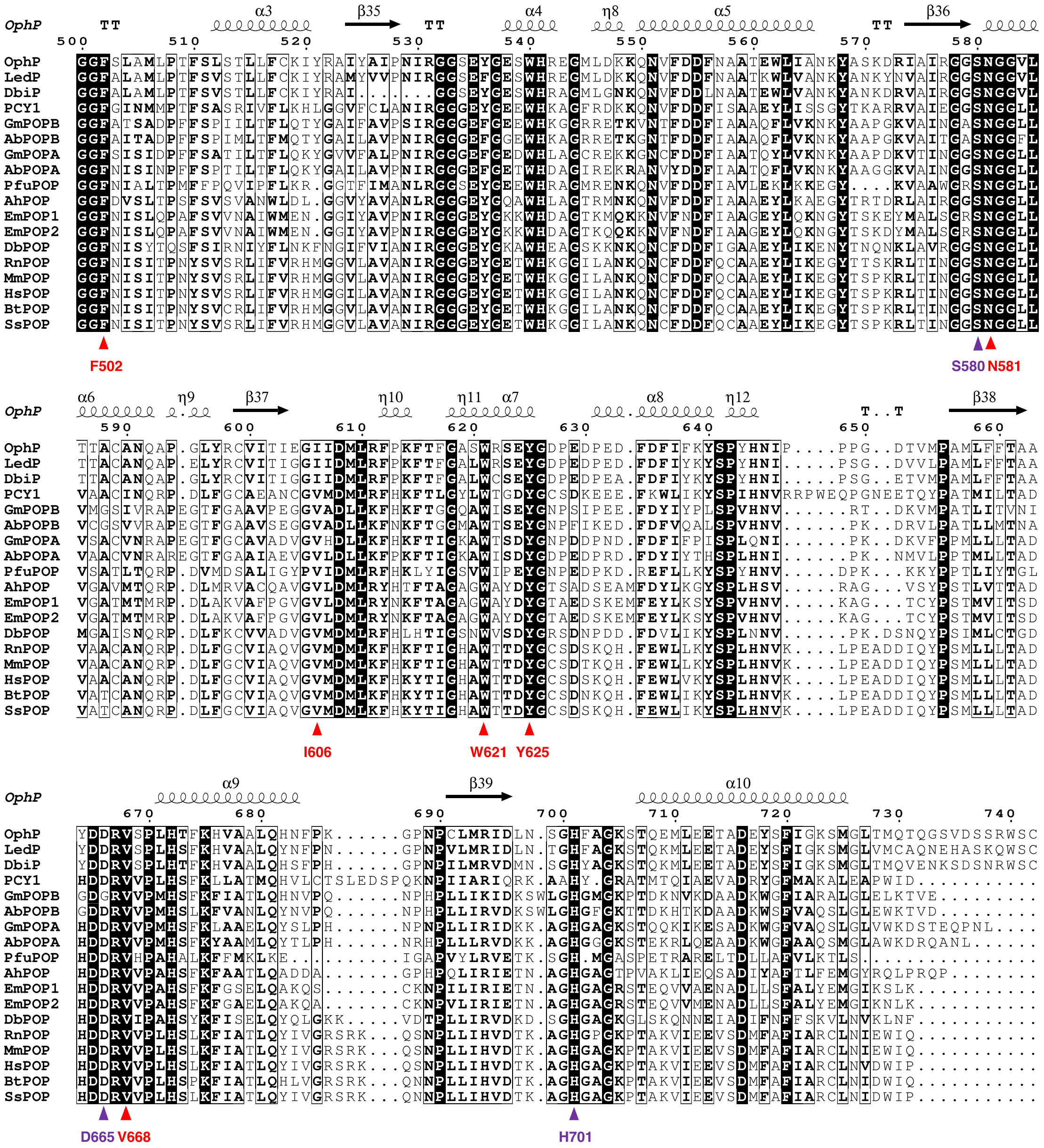

OphP
LedP
DbiP
PCY1
GmPOPB
AbPOPB
GmPOPA
AbPOPA
PfuPOP
AhPOP
EmPOP1
EmPOP2
DbPOP
RnPOP
MmPOP
HsPOP
BtPOP
SsPOP
F502
S580
N581
I606
W621
Y625
D665
V668
H701

**Figure 4-figure supplement 2. ITC experiments between OphP(S580A) and residues from the clasp domain (MPSSLLDAARESGEEASQNGFP), follower peptide (SVMSTE, and VIGSVMSTE), and 15mer.** Residues from the clasp domain (**a**) and follower peptides (**b** and **c**) showed no binding up to 2 mM in the presence of 50 μM of OphP(S580A). 15mer binds to OphP(S580A) with a $K_D$ of 1.5 μM (**d**). The positive value of ΔH showed that the binding is endothermic suggesting the exclusion of water solvent during binding.

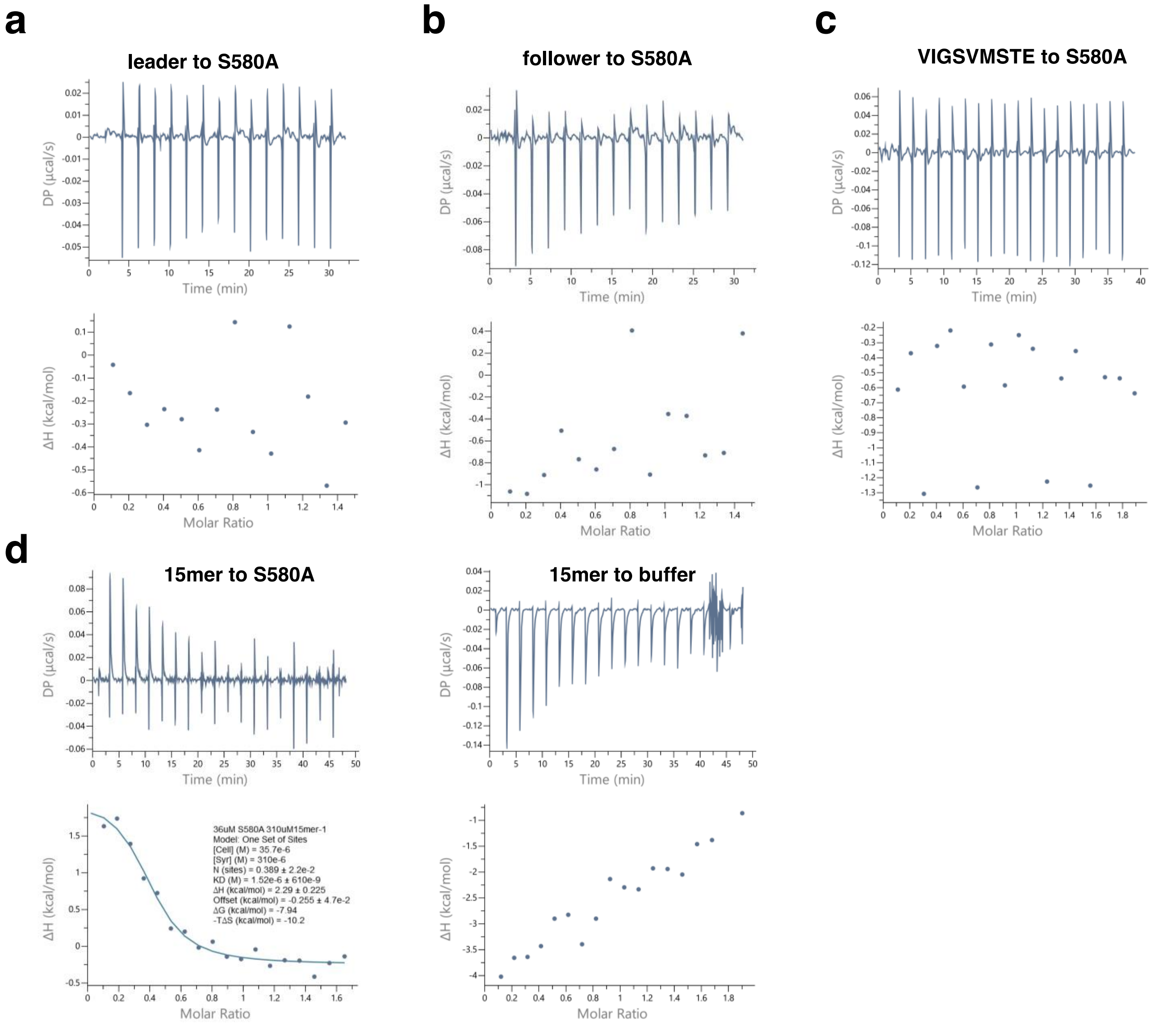

**Figure 4-figrue supplement 3. Crystal structure of the OphP-ZPP covalent inhibitor complex.** (a) Overall structure of the OphP-ZPP complex (b) $\sigma_A$-weighted electron density map (2mFo-DFc map contoured at 1 $\sigma$) showing the S580-acyl intermediate. (c). Superimposition of apo S580A (subunit A green and H grey, both have ordered loop region Leu697-Thr707) with OphP:ZPP complex (subunit A colored in salmon). The black arrows illustrate the large movement of two loops (Ser164-Met171, and Leu697-Thr707) when ZPP is bound. As a result of the 7.6 Å shift of His701 it is now part of the catalytic triad in the OphP:ZPP complex. The movement of Asp156 upon ZPP results in a salt bridge and Arg667 in the OphP:ZPP complex.

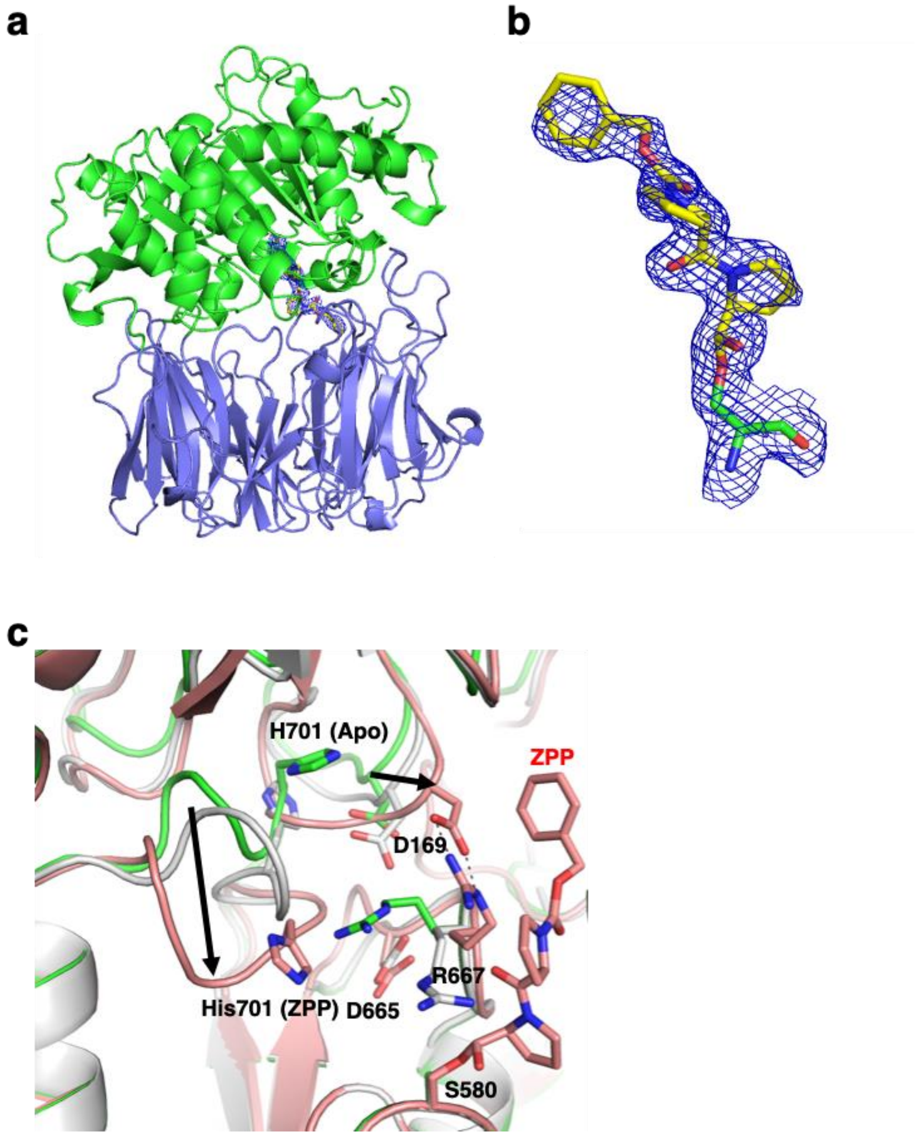

**Figure 4-figure supplement 4. Cartoon and electrostatic surfaces of apo GmPOPB (PDB 5N4F) and apo human dipeptidyl peptidase IV (DPP-IV, PDB 1NU6).** The view through the β-propellor and side views of (**a**) apo GmPOPB with the 35mer substrate (yellow spheres), and (**b**) apo DPP-IV with modelled tripeptide substrate (yellow spheres). Electrostatic calculation was performed using the APBS-PDB2PQR software suite under the default setting and drawn in PyMol. The electrostatic potential for the figures is set at ± 5 kT/e.

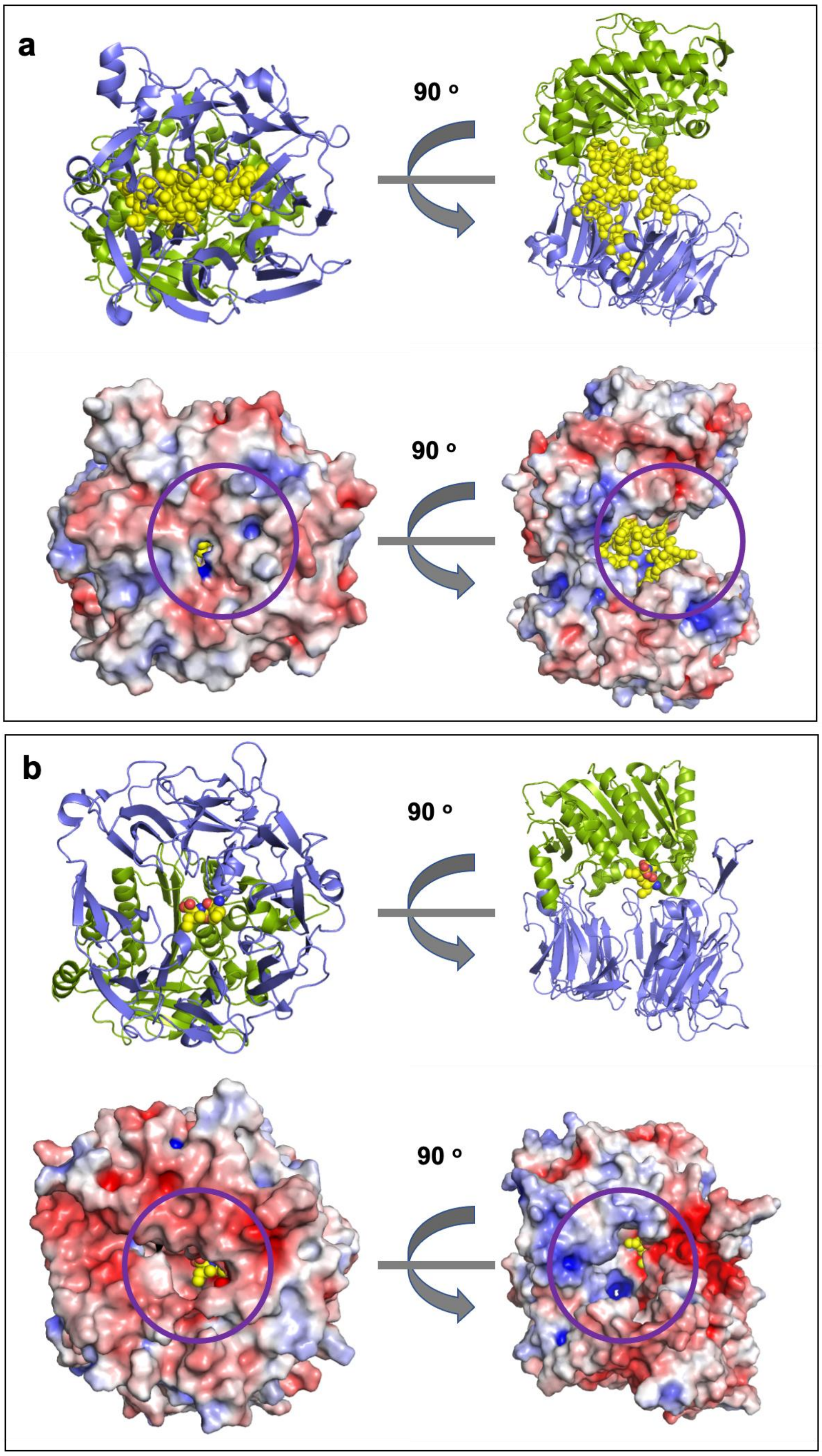
