## Supplementary material for "Molecular insight into the enzymatic macrocyclization of multiply backbone N-methylated peptides": Tables SOI

**Table S1**. List of oligonucleotides

| | |
|---|---|
| His-SUMOSTAR-FW | GAAGGATCCGAAACTATGGGTCATCATCATCATCATCATCATCATGGTGGTTCT GACTCCGAGGTC |
| SUMOSTAR-RV | AGACATACTAGTACCTTGGAAGTACAAGTTTTC |
| OphMA-NotI-RV | GAGTGCGGCCGCTTATTCCGTGCTCATGACTGATC |
| OphP-FW | AAGGTACTAGTATGTCGTTTCCAGGATGGGGACCAT |
| OphMA-FW | ATATACATATGGAGCATCATCATCATCATCATCATCATACTTCCACTC |
| OphMA-DbiCterm-RV | GCTCGAATTCTTACGCGCTGCTAACCACGCTTCCGATGACCCCAACGATACCCG TGACGATGACCCATGGGAAACC |
| OphMA-LedCterm-RV | CTCGAATTCTTACGCGCTGCTAACCACGCTACCAACCACGCCAACCACACCAAC CACAATGATCCATGGGAAACCTTGGAAGTAC |
| OphMA-ΔVIG-RV | GTAGCAGCAACCACAATAGCCAAGTCAGTACTCGTGCCTTATTCGCCGGCGGTC |
| OphMA-24mer-RV | TGACCCATGGGAAACCGTTTTGGGATTGGAAGTACAAGTTTTCACGAGCAGCGT CCAAGAGCGACG |
| OphP-I606A-FW | GAGGGTGCAATTGATATGCTTAGATTTCCTAAGTTTACATTTG |
| OphP-I606A-RV | CATATCAATTGCACCCTCAATTGTGATCACACAACGGTAGAG |
| OphP-W621A-FW | GCTAGCGCACGTTCTGAATATGGCGATCCTGAGGATCCTG |
| OphP-W621A-RV | TCAGAACGTGCGCTAGCGCCAAATGTAAACTTAGGAAATCTA |
| OphMAΔC6-G408A-FW | GTCGTTGGTGTTATCGCTGTCATCGGATAAGCGGCCGC |
| OphMAΔC6-G408A-RV | AGCGATAACACCAACGACGATGACCCATGGGAAACCGTTT |
| OphMAΔC6-G408V-FW | GTCGTTGGTGTTATCGTCGTCATCGGATAAGCGGCCGC |
| OphMAΔC6-G408V-RV | GACGATAACACCAACGACGATGACCCATGGGAAACCGTTT |
| OphMAΔC6-G408L-FW | GTCGTTGGTGTTATCCTCGTCATCGGATAAGCGGCCGC |
| OphMAΔC6-G408L-RV | GAGGATAACACCAACGACGATGACCCATGGGAAACCGTTT |

**Table S2**. List of plasmids

| Name | # | Reference/Source |
|---|---|---|
| pPIC3.5K-strepII-SUMO*-TEVcs-OphP | PMA1382 | (Matabaro et al., 2021) |
| pPIC3.5K-strepII-SUMO*-TEVcs-OphP | PMA1379 | (Matabaro et al., 2021) |
| pCDFDuet-sMBP-OphP(S580A) | PMA1374 | E. Matabaro, unpublished |
| pPIC3.5K-His$_8$-SUMO*-TEVcs-OphP | PMA1596 | This study |
| pPIC3.5K-His$_8$-SUMO*-TEVcs-LedP | PMA1595 | This study |
| pPIC3.5K-His$_8$-SUMO*-TEVcs-OphP(S580A) | PMA1597 | This study |
| pPIC3.5K-His$_8$-SUMO*-TEVcs-OphP(I606A) | | This study |
| pPIC3.5K-His$_8$-SUMO*-TEVcs-OphP(W721A) | | This study |
| pET24-His$_8$-OphMA(cDNA) | PMA1004 | (van der Velden et al., 2017) |
| pET24-His$_8$-OphMA-TEVcs-ΔC6(15mer*) | PMA1304 | (van der Velden et al., 2017) |
| pET24-His$_8$-OphMA-TEVcs-ΔVIG(18mer*) | PMA1637 | This study |
| pET24-His$_8$-OphMA-TEVcs(21mer*) | PMA1488 | (van der Velden et al., 2017) |
| pET24-His$_8$-OphMA-TEVcs-24mer* | PMA1640 | This study |
| pET24-His$_8$-OphMA-TEVcs-ΔC6(G408A) | | This study |
| pET24-His$_8$-OphMA-TEVcs-ΔC6(G408V) | | This study |
| pET24-His$_8$-OphMA-TEVcs-ΔC6(G408L) | | This study |

*Size of peptide released by TEV protease

**Table S3**. Data collection, phasing and refinement statistics for all the structures.

| | Apo S580A | OPhP-ZPP | S580A:Oph-15mer | S580A:Oph-18mer |
|---|---|---|---|---|
| PDB Entry | | | | |
| **Data collection** | | | | |
| Space group | P 1 | P 1 | P 1 | P 1 |
| Cell dimensions | | | | |
| *a*, *b*, *c* (Å) | 69.86, 113.43, 186.32 | 69.68, 102.65, 110.27 | 70.22,106.18, 114.79 | 70.01, 104.61, 110.65 |
| α, β, γ (°) | 83.97, 82.09, 76.93 | 116.23, 101.09, 92.18 | 113.04, 101.67, 93.24 | 115.81, 98.87, 93.79 |
| Wavelength (Å) | 0.9795 | 0.9763 | 0.9763 | 0.9762 |
| Resolution (Å) | 66.00-1.94<br>(1.97-1.94) | 67.71-2.00<br>(2.03 -2.00) | 64.35-2.47<br>(2.51-2.47) | 55.84 -2.00<br>(2.03-2.00) |
| $R_{merge}$ | 0.075 (0.993) | 0.115 (1.099) | 0.120 (1.375) | 0.090 (1.234) |
| $I / \sigma I$ | 7.3 (0.9) | 14.3 (1.2) | 13.0 (0.9) | 15.0 (1.0) |
| Completeness (%) | 97.6 (96.6) | 97.6 (92.3) | 98.6(97.96) | 98.0 (96.9) |
| Redundancy | 2.5 (2.4) | 3.5 (3.4) | 3.6 (3.6) | 3.3 (3.5) |
| No. Unique reflections | 999365 (397483) | 611334 (175661) | 370936 (104483) | 598522 (182773) |
| $CC_{1/2}$ | 0.996 (0.315) | 0.990 (0.580) | 0.993 (0.530) | 0.996 (0.545) |
| **Refinement** | | | | |
| Resolution (Å) | 66.00-1.94<br>(1.97-1.94) | 67.71-2.0<br>(2.03-2.0) | 64.35-2.47<br>(2.51-2.47) | 55.84-2.0<br>(2.03 -2.0) |
| $R_{work}$ / $R_{free}$ | 0.214/0.241 | 0.210/0.245 | 0.227/0.263 | 0.201/0.235 |
| No. atoms | | | | |
| Protein | 46235 | 22799 | 22933 | 23204 |
| Ligand/ion | 89 | 257 | 276 | 251 |
| Water | 898 | 1328 | 53 | 682 |
| *B*-factors (Å$^2$) | | | | |
| Protein | 45.91 | 36.6 | 76.66 | 47.46 |
| Ligand/ion | 60.91 | 56.4 | 103.09 | 76.67 |
| Water | 36.98 | 36.6 | 61.90 | 39.54 |
| R.m.s deviations | | | | |
| Bond lengths (Å) | 0.007 | 0.006 | 0.007 | 0.006 |
| Bond angles (°) | 1.354 | 1.317 | 1.421 | 1.332 |
| Ramachandran | | | | |
| Allowed (%) | 99.75 | 99.72 | 99.79 | 99.83 |
| Outliers (%) | 0.25 | 0.28 | 0.21 | 0.17 |

*Values in parentheses are for highest-resolution shell.
