## Supplementary material for "Molecular insight into the enzymatic macrocyclization of multiply backbone N-methylated peptides": Fig1SOI

**Figure 1-figure supplement 1**. **OphP production in *Pichia pastoris*.** (**a**) OphP was fused to a His8-SUMO* via a cleavage site for TEV protease (in red). Residues Thr and Ser after the TEV cleavage site resulted from the insertion of the restriction site *Spe*I between OphP and the tag. The arrow indicates the cleavage site for TEV protease. (**b**) Purification of OphP. After purification of $His_8$SUMO*-OphP using metal affinity chromatography (lane MA), the tag was cleaved off by incubation of the purified fusion protein with TEV protease at room temperature (lane RT+TEV). We noted that the tag was also slowly cleaved off even without the addition of TEV protease (lane RT). The P1 site of latter cleavage was mapped to residue Thr117 (see panel (**a**)) by whole protein MS. The OphP protein without tag was further purified by size exclusion chromatography (SEC) using an Aekta FPLC system. (**c**) OphP peak from SEC purification consistent with a monomer. mAU: Milli Absorbance Unit. (**d**) Heterologous production of OphP homologs LedP and GmPOPB. Recombinant LedP was produced in *P. pastoris* (this study) and GmPOPB in *E. coli* (Czekster et al., 2017).

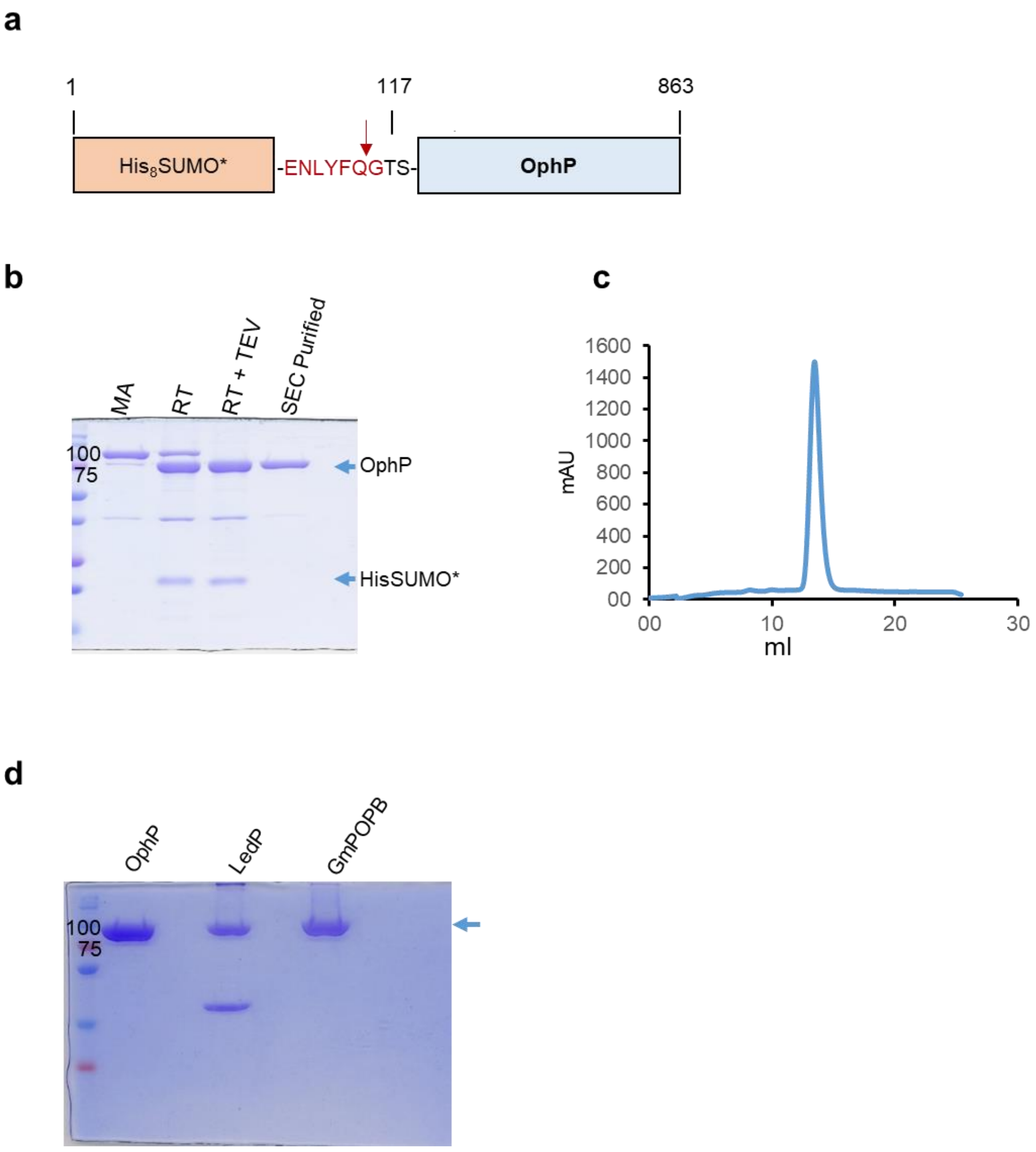

**Figure 1-figure supplement 2. OphP activity on the chromogenic substrate Z-Gly-Pro-pNA.** (**a**) The effect of temperature on OphP activity. The reactions were run in the standard assay buffer (50 mM HEPES pH 6.0 + 10 mM DTT). (**b**) The effect of pH on OphP activity (at 30°C). In both assays (a) and (b), the concentration of the substrate and the enzyme was 1 mM and 8 µM, respectively. All assays were performed in triplicates overnight. The enzymatic activity was monitored by absorbance measurement at 410 nm on an Infinite 200 Pro M Plex spectrophotometer. Error bars are standard errors of the mean absorbance. (**c**) Conversion kinetics of Z-Gly-Pro-pNA by OphP. On the left, the calibration curve of Z-Gly-Pro-pNA, and on the right, enzyme saturation assay are shown. For the substrate calibration curve, the reactions were incubated at 30°C with 8 µM OphP for 22h. A linear curve was fitted to the data points and the resulting linear equation $A_{410} = \varepsilon m \times C + b$ was used to calculate pNA concentration as a function of absorbance ($C = (A - b) / \varepsilon m$, where εm is the molar extinction coefficient of pNA. The curve was fitted with an R-squared value of 0.9831. For the enzyme saturation assay, pNA product was measured at various times for four different OphP concentrations against 1mM of substrate. The curves for the 2x and 5x relative levels of OphP were outside of the linear range after 4h and 10h, respectively. On the subsequent *in vitro* experiments, 1x[E] (8uM) was used. (**d**) The effect of the addition of Z-Pro-prolinal and the S580A mutation on OphP activity. OphP and OphP(S580A) (8 µM) were incubated with Z-Pro-prolinal (ZPP) (67 µM) for 30min at 30°C in the standard assay buffer as stated above, before adding the chromogenic substrate 1 mM) for the indicated time period.

**a**

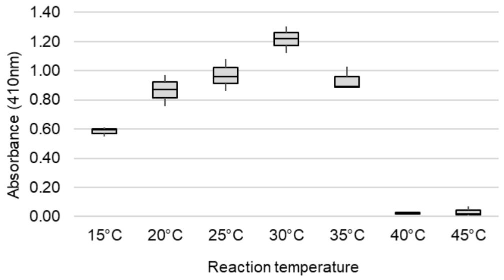


**b**

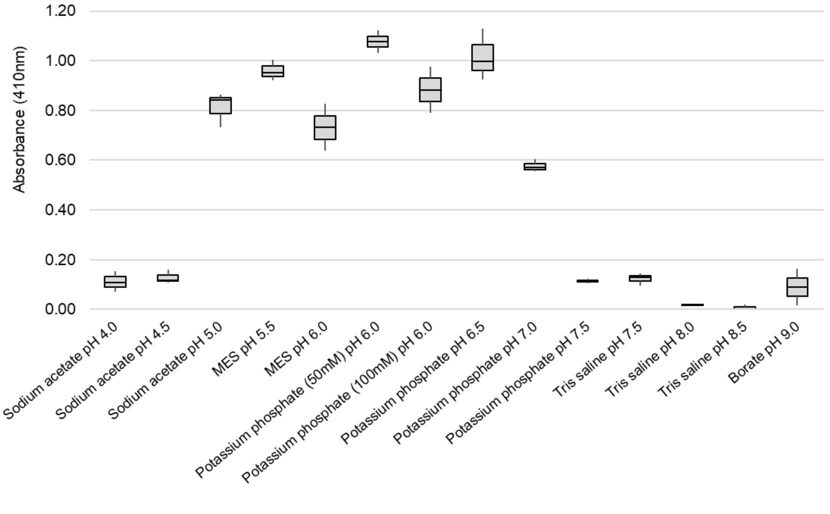

c

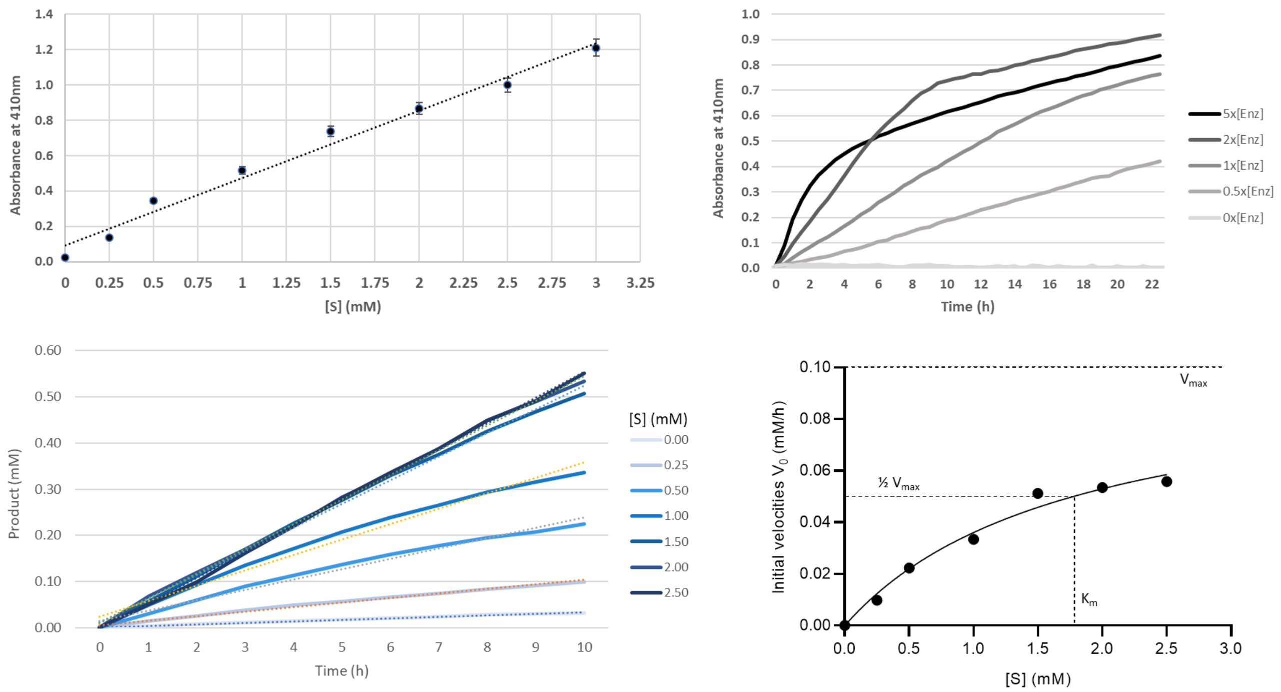
Absorbance at 410nm
[S] (mM)
Absorbance at 410nm
Time (h)
5x[Enz]
2x[Enz]
1x[Enz]
0.5x[Enz]
0x[Enz]
Product (mM)
Time (h)
[S] (mM)
0.00
0.25
0.50
1.00
1.50
2.00
2.50
Initial velocities V0 (mM/h)
[S] (mM)
Vmax
½ Vmax
Km

d

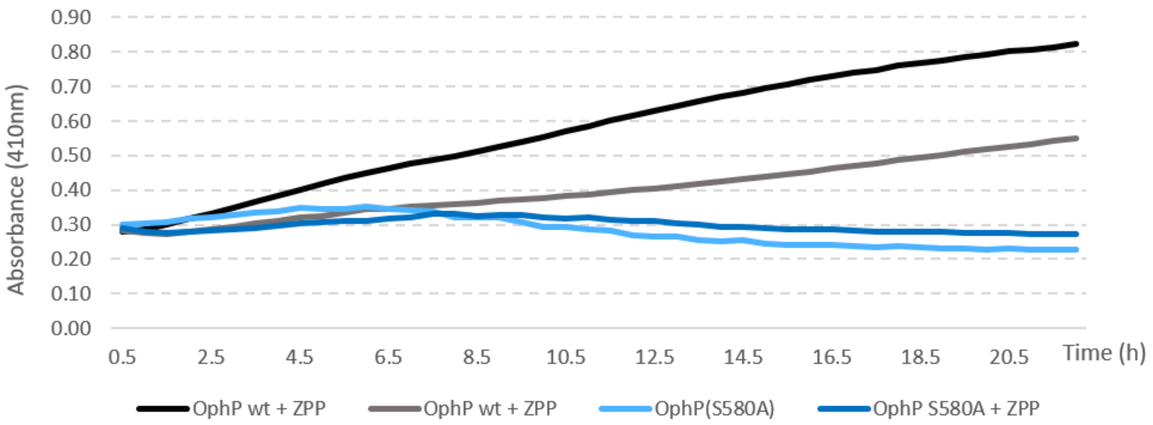
Absorbance (410nm)
Time (h)
OphP wt + ZPP
OphP wt + ZPP
OphP(S580A)
OphP S580A + ZPP

**Figure 1-figure supplement 3**. **Lack of conversion of completely methylated, full-length OphMA by OphP *in vitro***. (**a**) ESI-MALDI-ToF analysis of purified OphMA co-incubated with methyl donor substrate SAM and purified OphP *in vitro*. Full spectra can be found in panels b, c and d. Dashed red lines indicate the various methylation states of OphMA. (**b**) Intact mass of full length (fl) $His_8$OphMA expressed in *E. coli* for 2 h at 16$^{o}$C. (**c**) Intact mass of full length $His_8$OphMA-2h when incubated at 4 mg/ml with 5 mM SAM at room temperature in 50 mM Tris pH 8.0, 100 mM NaCl and 1 mM TCEP. (**d**) Intact mass of full length $His_8$OphMA-2h incubated at 4 mg/ml with 5 mM SAM and 0.9 mg/ml OphP (10 mM) at room temperature in 50 mM Tris pH 8.0, 100 mM NaCl and 1 mM TCEP shows no evidence of cleavage. The molecular mass of OphP (after TEV cleavage) is 84611 Da and the peak at 42303 Da corresponds to doubly charged OphP. In panels b, c, d the y axis is the % of the highest signal and a the representation is the deconvoluted spectrum (thus mass not m/z is used). The raw data from experimental are shown in red above the spectra.

**a**

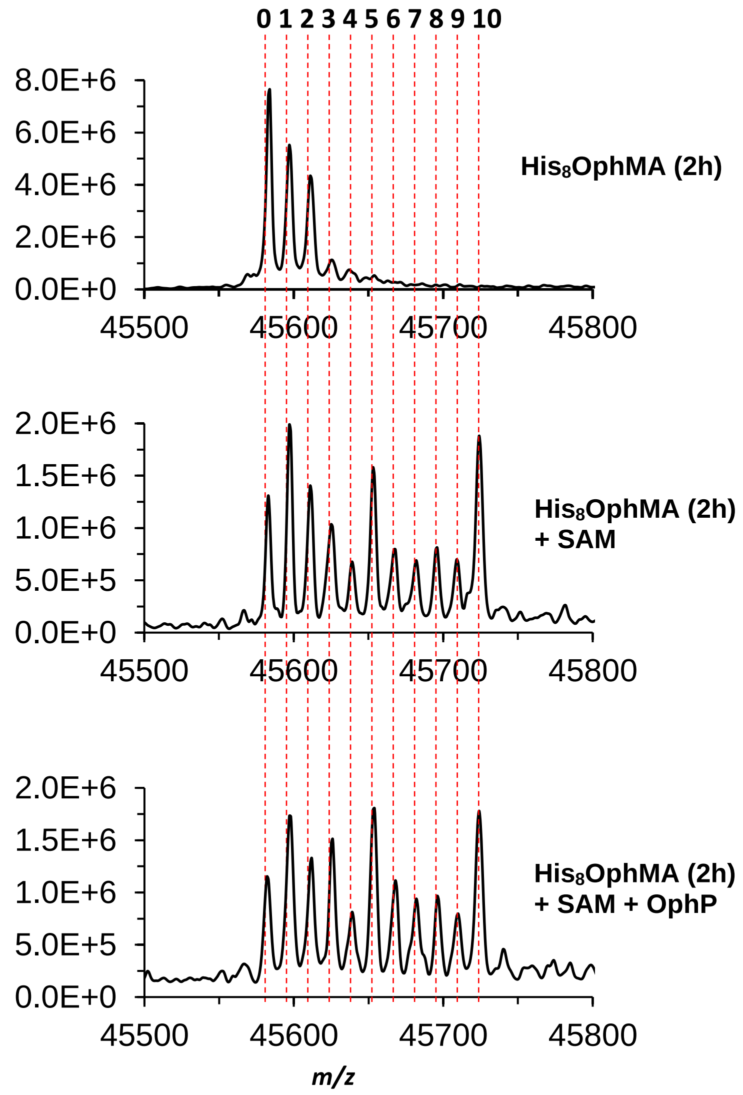

**b**

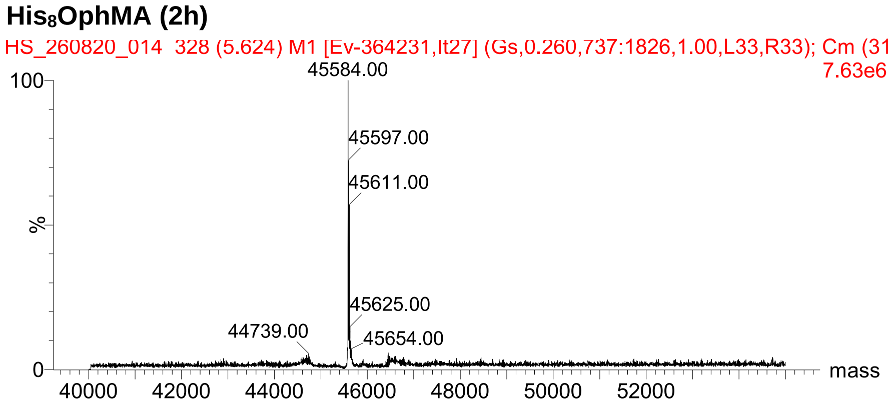


**c**

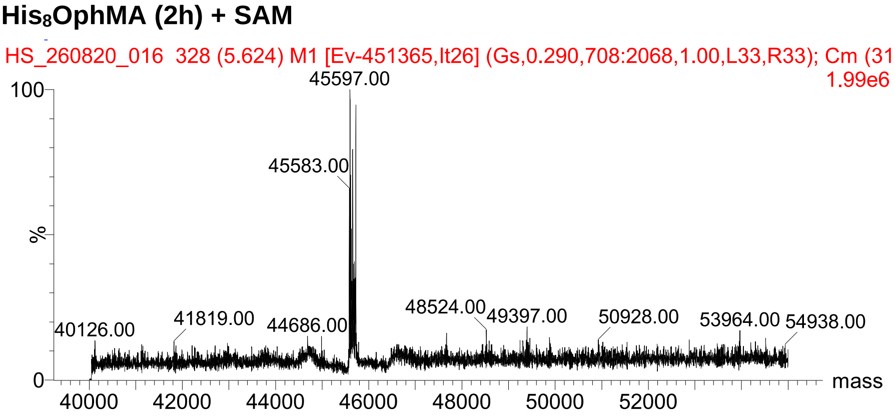


**d**

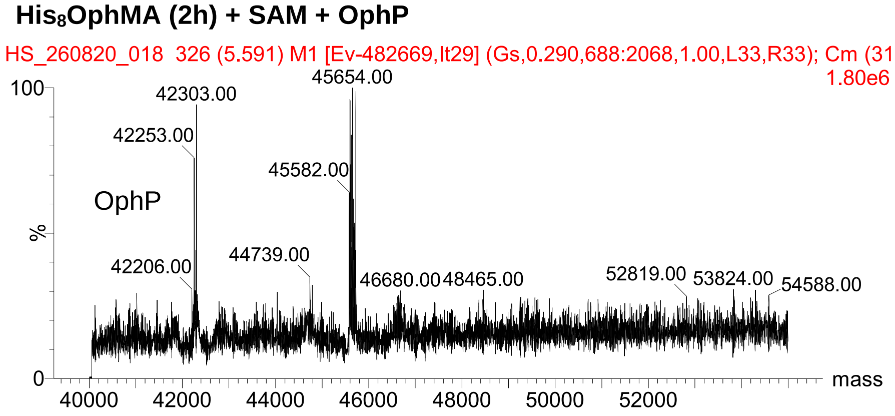
